# Conformational Landscapes of a Class I Ribonucleotide Reductase Complex during Turnover Reveal Intrinsic Dynamics and Asymmetry

**DOI:** 10.1101/2024.06.16.599213

**Authors:** Da Xu, William C. Thomas, Audrey A. Burnim, Nozomi Ando

## Abstract

Understanding the structural dynamics associated with enzymatic catalysis has been a long-standing goal of biochemistry. With the advent of modern cryo-electron microscopy (cryo-EM), it has become conceivable to redefine a protein’s structure as the continuum of all conformations and their distributions. However, capturing and interpreting this information remains challenging. Here, we use classification and deep-learning-based analyses to characterize the conformational heterogeneity of a class I ribonucleotide reductase (RNR) during turnover. By converting the resulting information into physically interpretable 2D conformational landscapes, we demonstrate that RNR continuously samples a wide range of motions while maintaining surprising asymmetry to regulate the two halves of its turnover cycle. Remarkably, we directly observe the appearance of highly transient conformations needed for catalysis, as well as the interaction of RNR with its endogenous reductant thioredoxin also contributing to the asymmetry and dynamics of the enzyme complex. Overall, this work highlights the role of conformational dynamics in regulating key steps in enzyme mechanisms.

## Introduction

Ribonucleotide reductases (RNRs) provide the sole *de novo* synthesis pathway for the building blocks of DNA in nature by catalyzing the conversion of ribonucleotides to 2′-deoxyribonucleotides^1^. Despite wide evolutionary diversity^2,3^, all RNRs share a common catalytic mechanism that is initiated by the formation of a thiyl radical on a conserved active-site cysteine^4^. The oxygen-dependent class I RNRs^5,6^ are used by aerobic life, including most eukaryotes, many prokaryotes, and associated viruses^2,3^. These RNRs have long fascinated researchers for their relevance as drug targets and for their dynamic quaternary structure involving two subunits (Figure 1A): 1) a small ferritin-like subunit (β) that houses a stable radical cofactor, and 2) a large catalytic subunit (ɑ) that contains the nucleotide-binding sites, including the active site, an allosteric site that controls substrate specificity, and additional allosteric sites involved in regulating overall enzyme activity.

**Figure 1.**
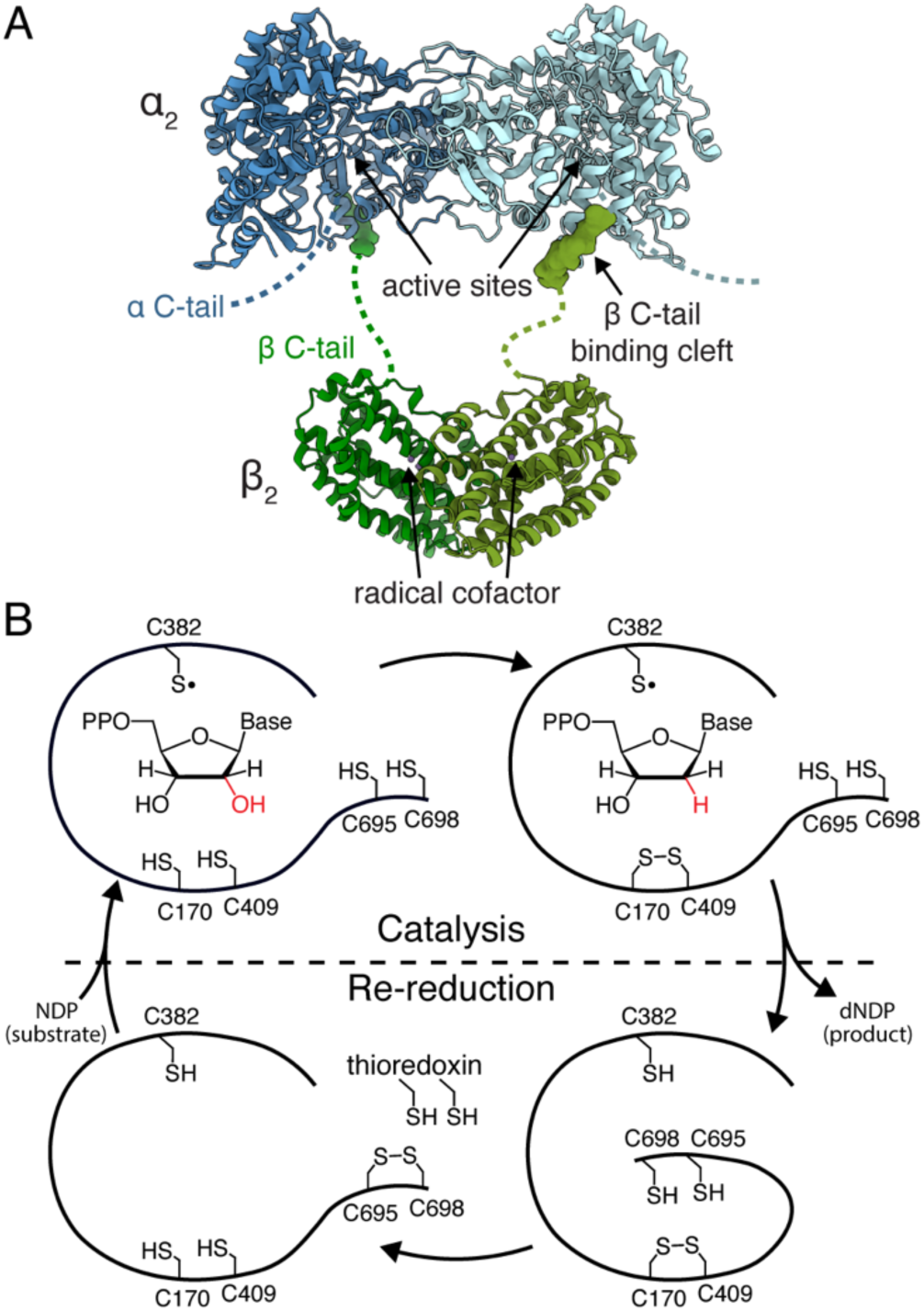
Class I ribonucleotide reductases (RNRs) require dynamic components for turnover. (A) The α_2_ subunit in *Bacillus subtili*s (*Bs*) class Ib RNR (pdb: 6mt9, blue) houses the active site and allosteric sites^43^. The β_2_ subunit (pdb: 4dr0, green) houses a dimanganese tyrosyl radical cofactor^36^. Both subunits are homodimeric and have functionally essential C-terminal tails, which are flexible (dashed). The two subunits are stably tethered through a strong interaction between the β C-terminus and its binding cleft on the α subunit (pdb: 6mw3, surface), but the interaction between the core parts of α_2_ and β_2_ is very dynamic. (B) The turnover cycle of class I RNRs with *Bs* numbering. PCET between the two subunits generates a thiyl radical (C382•) in the active site, which initiates reduction of ribonucleoside diphosphates to 2ʹ-deoxyribonucleoside diphosphates. Active-site cysteines (C170/C409), which are oxidized during substrate reduction, are reduced by cysteines on the α C-terminal tail (C695/C698), which are in turn reduced by external reductants such as thioredoxins.

Decades of research, especially on the *Escherichia coli* (*Ec*) class Ia RNR, have delineated the turnover cycle of class I RNRs^1,7^, which consists of two halves: catalysis and re-reduction (Figure 1B). The catalysis half-cycle begins with proton-coupled electron transfer (PCET) along a series of conserved amino acids that transfers the radical from the cofactor in the β subunit to the catalytic cysteine in the α subunit (C382 in Figure 1B). Radical transfer initiates reduction of the substrate and concomitant oxidation of an additional pair of active-site cysteines (C170/C409 in Figure 1B). The radical then transfers back to the β subunit. In the re-reduction half-cycle, the oxidized active-site cysteines are reduced by a cysteine pair located on the flexible α C-terminal tail (C695/C698 in Figure 1B), which is in turn reduced by external reductants, such as thioredoxins, resetting the enzyme for a new round of catalysis. For PCET to occur, the class I subunits interact as dimers (α_2_ and β_2_) to form an active α_2_β_2_ complex (Figure 1A), but this complex has proven to be highly dynamic, posing a major challenge for high-resolution structural characterization. Recently, the structure of an active class I RNR complex was reported by cryo-electron microscopy (cryo-EM) using unnatural amino acid incorporation to trap the enzyme in a PCET-competent conformation^8^. This landmark study revealed striking asymmetry within the *Ec* class Ia RNR complex, with α and β closely engaged on one side and the flexible β C-terminus folded into the active site to contribute an interfacial PCET residue, thereby providing a structural basis for multiple lines of evidence that suggest half-site reactivity^9–17^. Despite these remarkable advances, the full set of conformations needed for enzymatic turnover have remained enigmatic. Moreover, direct links cannot be made between discrete structural states and enzyme kinetics.

A major promise of single-particle cryo-EM is its ability to resolve the conformational heterogeneity of proteins, as it is a single-molecule imaging technique by essence, which directly measures the signals of individual molecules. Traditionally, this heterogeneity has been addressed through the 3D classification process that involves sorting particle images into subsets and reconstructing them separately based on the assumption of a discrete number of conformations. Over the past decade, significant algorithmic advances have enabled the modeling of continuous heterogeneity in cryo-EM datasets with methods such as manifold embedding^18^, cryoDRGN^19^, 3D variability analysis^20^, e2gmm^21^, 3DFlex^22^, Opus-DSD^23^, RECOVAR^24^, DRGN-AI^25^, among others. Of these, cryoDRGN stands out as one of the most widely used tools that has been applied to a diverse range of datasets^19,26–31^. CryoDRGN utilizes a variational auto-encoder (VAE) architecture to directly learn a mapping from single particle images with known poses to their corresponding 3D volumes using multi-layer perceptrons (MLPs), which have been shown to effectively model both compositional and conformational heterogeneity. Once a cryoDRGN model is trained on a dataset, conformational landscape analysis^27,32^ can be performed to estimate relative populations of different conformational species. This is typically achieved by generating 3D volumes at *k*-means clusters in the latent space and performing principal component analysis (PCA) on those volumes^32^. The principal components offer insight into the most significant modes within a dataset as well as a way to reduce the dimensionality of the landscape for visualization in 2D, where cluster volumes are mapped along two principal components. However, comparing landscapes across multiple datasets by this “volume-PCA” approach can be challenging as different microscope settings can introduce variations in pixel size and noise level and there may be inherent sample-to-sample differences.

Here, we use cryo-EM to investigate the conformational landscapes of the class I RNR from *Bacillus subtilis* (*Bs*) during turnover under native-like solution conditions. *Bs* RNR achieves activities comparable to *Ec* class Ia RNR using a class Ib cofactor (a dimanganese tyrosyl radical)^33^. Moreover, we had previously shown by small-angle X-ray scattering (SAXS) that *Bs* RNR forms a nearly homogenous solution of α_2_β_2_ tetramers due to the substantially stronger α/β affinity^33^ compared to the *Ec* enzyme^34^. This property allows us to obtain high-quality cryo-EM micrographs of the active *Bs* RNR complex. Through 3D classification, we reconstruct a series of α_2_β_2_ classes depicting a continuous range of conformations, from open to compact, including one that closely resembles the PCET-competent conformation observed in *Ec* class Ia RNR, pointing to a shared PCET conformation across all class I RNRs. Using cryoDRGN^19^ and landscape analysis, we then further characterize the continuum of conformations sampled by the enzyme. Rather than taking the volume-PCA based approach, we visualize the landscape by rigid-body fitting atomic models of α_2_ and β_2_ subunits into the *k*-means cluster volumes and then directly parameterizing their relative orientations as high-dimensional vectors, which are embedded in 2D via a density-preserving dimensionality reduction method, DensMAP^35^. This process allows us to convert our cryoDRGN results into physically interpretable 2D conformational landscapes and directly compare landscapes between datasets collected during different microscope sessions. The resulting landscapes demonstrate that RNR continuously samples a multitude of conformations while maintaining structural asymmetry throughout each step of turnover, revealing an intricate mechanism for half-site reactivity. Furthermore, the landscapes enable direct visualization of how the presence of substrates and products controls the accessibility of certain conformations, shedding light onto the dynamic regulation of enzyme turnover.

## Results

### Asymmetry and dynamics in RNR turnover

To characterize the motions involved in RNR turnover, we examined a cryo-EM dataset collected on a sample flash-frozen within 20 s of initiating turnover in *Bs* RNR in the presence of its endogenous reducing system, thioredoxin (TrxA) and NADPH-dependent thioredoxin reductase (TrxB) (Methods Section 2). The reaction mixture also contained nucleotides that were shown to yield high activity (Methods Section 1): substrate guanosine diphosphate (GDP), specificity effector thymidine triphosphate (TTP), and activator adenosine triphosphate (ATP). Following multiple rounds of particle filtering, a 3D consensus reconstruction was performed on the full set of high-quality α_2_β_2_ particles (Methods Section 3). As expected from a dataset with a high degree of heterogeneity, the resulting consensus reconstruction displays varied resolution across different parts of the map (Figure 2A-B, Table S1), where particles are aligned on the larger α_2_ subunit (160 kDa) to ∼2.5-Å resolution, while the smaller β_2_ subunit (80 kDa) is blurred. Despite the blurring, this smaller density is consistent with the crystal structure of the ferritin fold of the *Bs* β_2_ subunit (residues 16-290)^36^. In class I RNRs, the β subunit also contains a long disordered C-terminal tail (residues 291-329 in *Bs* RNR), where the residues near the very C-terminus are essential for binding the α subunit^37,38^ (Figure 1A) and the intervening residues (residues 291-308 in *Bs* RNR) are responsible for entering the active site of the α subunit during PCET^8^. Interestingly, we observe high-resolution density for 14 residues (K309 – F322) of the β C-terminus bound on both sides of α_2_ (Figure 2B-C), indicating that the blurring of β_2_ is not caused by low occupancy, but rather by a dynamic β_2_ subunit that is tethered to α_2_ via its C-terminal tails (Figure 1A). The presence of β tail density on both sides of *Bs* RNR contrasts with the cryo-EM map of the trapped *Ec* RNR, where this density was observed on only one side^8^, and may account for the significantly stronger inter-subunit affinity observed in the *Bs* enzyme.

**Figure 2.**
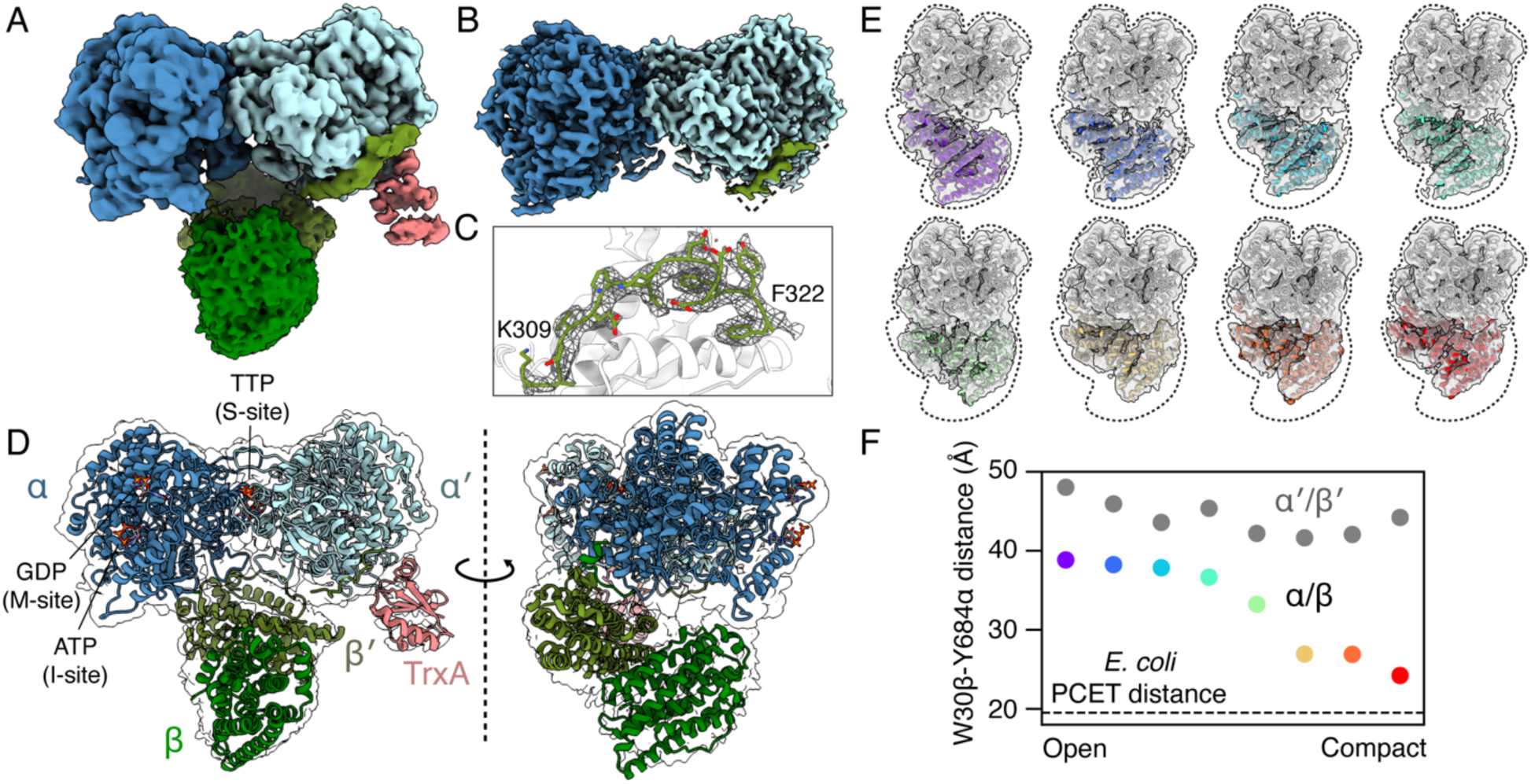
Cryo-EM reconstructions of *Bs* RNR during active turnover reveal continuous motions that maintain functional asymmetry. A multiple-turnover reaction mixture of 5 µM α, 5 µM β, 30 µM TrxA, 1 µM TrxB, 0.25 mM TTP, 1 mM ATP was initiated by addition of substrate GDP at a concentration of 0.5 mM. (A) Unsharpened consensus map contains density for α_2_ (blue) and β_2_ (green), as well as additional density on one side of α_2_ corresponding to TrxA (coral). (B) Sharpened consensus map contains high-resolution information for α_2_ and residues 309-322 of the β C-termini (green surface in dashed box). (C) The bound β C-terminus (mesh) is well-resolved on both sides of α_2_. (D) Consensus model of the *Bs* RNR complex produced by docking a refined model of α_2_ with the β C-terminus and allosteric effectors (ATP, TTP, GDP) bound to each monomer and the atomic models of β_2_ and TrxA into the unsharpened map (Methods, Supplementary Information Section S3). Surprising asymmetry is observed in β_2_ and TrxA binding. (E) 3D classification reveals continuous motion of β_2_ from open to compact (left to right, top to bottom), shown in same side-view orientation as in panel D. (F) Asymmetry is observed across all 3D classes, as shown by distance between interfacial residues involved in PCET: W30β and Y684α. Only one side of the RNR complex (α/β pair) can approach the proximity needed for PCET, while the other side (α′/β′ pair) remains separated by a long distance. Colors of individual points are consistent with panel E.

A consensus model of the full complex was built by refining an atomic model of α_2_ bound with nucleotides and β C-termini against the sharpened map and docking the β_2_ crystal structure in the unsharpened map (Figure 2D). The final consensus model is reminiscent of the *Ec* RNR structure^8^, where β_2_ is asymmetrically bound to α_2_, albeit in a more open and less engaged conformation. Consequently, the monomers within α_2_ and β_2_ become distinct, and we can associate each β subunit with the α subunit where its tail binds to define the two halves of the complex as α/β and α′/β′ (Figure 2D). Focused 3D classification of β_2_ was then performed to sort the heterogeneous α_2_β_2_ particles in the turnover reaction mixture by conformation. This process resulted in 8 well-resolved classes that reveal a continuous range of α_2_β_2_ conformations, from open to compact (Figure 2E, Table S2). In the 3D classes, clear secondary structural details are observed in the β_2_ density that align with the crystal structure, supporting the interpretation that the blurring in the consensus map was due to the averaging of a highly mobile β_2_ subunit. Notably, the PCET conformation seen in *Ec* RNR is not observed in any of these 3D classes.

In addition to asymmetrically bound β_2_, we observed a small feature in the consensus map on only one side of α_2_ (Figure 2A). Upon examining the remaining protein components in the reaction mixture, we found that an atomic model of TrxA fits unambiguously into the density (Figures 2D, S1A), revealing a thioredoxin-RNR interaction that had previously remained elusive. This direct interaction between α_2_ and TrxA may explain previous biochemical findings that using TrxA as the reducing agent increases the catalytic activity of *Bs* RNR by 10-fold compared to using the small molecule reductant 1,4-dithiothreitol (DTT)^33^. Like the β_2_ density, TrxA density disappears with sharpening, in this case owing to a low binding affinity that depends on the enzyme oxidation state, as will be detailed later. Nonetheless, the asymmetric binding of β_2_ and TrxA has profound implications for the half-site reactivity of the class I RNR mechanism. PCET requires interfacial residues between α_2_ and β_2_ (W30β → Y307β → Y684α) to come in proximity^1^. While Y307β is on the unresolved region (residues 291-308) of the β C-terminal tail, it is two residues away from the segment bound to α_2_ near the active-site opening, and therefore is likely to be close to Y684α (Figure S2A). However, the asymmetric posing of β_2_ relative to α_2_ causes large differences in W30β-Y684α separation on the two sides of the complex. On one side (α/β), W30β-Y684α distances appear to approach the distance needed for the disordered region of the β C-terminus to fold into the active site for PCET (Figure 2F). In contrast, distances on the other side (α′/β′) are consistently too long for PCET, as evidenced by the β tail density in certain 3D classes appearing stretched across this gap (Figure S2B). Curiously, TrxA, which is required for re-reduction, preferentially binds to the α′/β′ side where PCET cannot occur. This asymmetry suggests that the enzyme has evolved a mechanism for segregating catalysis and re-reduction processes across its two halves.

### Shared dynamics during different stages of turnover

To understand how the conformational ensemble is coupled to the turnover process, we collected three additional cryo-EM datasets with slight alterations to the reaction mixture (Methods Section 2, Table S1): a pre-turnover condition (lacking substrate), a product condition (replacing substrate with product), and a pre-reduction condition (stalled re-reduction due to depletion of NADPH after multiple turnovers). For all three datasets, we obtained 3D consensus reconstructions showing similarly blurry β_2_ but with varying TrxA occupancy (Figure S3, Tables S1-S2). Consistent with the expectation that the enzyme remains reduced in the pre-turnover and product conditions (due to lack of turnover), density for the active-site C170/C409 pair appears largely reduced, while they are largely oxidized in the pre-reduction condition and a mixture of reduced and oxidized in the turnover condition (Figure S4). Interestingly, TrxA occupancy in the consensus maps appears to be correlated with the level of active-site oxidation, with the pre-reduction condition exhibiting the strongest density, followed by the turnover condition, and the two non-turnover conditions (pre-turnover, product) having similarly low occupancies (Figure S3). For all conditions, blurry density is observed in the active site (Figure S5), which can be attributed to an average of nucleotides and the disordered C-terminal tail of the α subunit (residues 688-700, Figure 1A) competing for the same pocket, except in the pre-turnover condition where the blurring corresponds solely to the transient α-tail. In contrast with the dynamic nature of the active site, well-defined nucleotide density is observed in the allosteric sites (Figure S6).

Focused 3D classification of β_2_ yielded 3-8 well-resolved classes per condition that resemble the continuum of open to compact conformations seen in the turnover condition, suggesting that there are intrinsic conformational dynamics that are shared even in the absence of turnover (Figure S7A-D, Table S2). Remarkably, 3D classification of particles combined from all conditions yielded a more finely sampled range of this continuum (Figure S7E, Table S2), further supporting the presence of particles that adopt similar conformations across different datasets. Inspection of representative classes (Figure 3) provided insight into the source of asymmetry within the α_2_β_2_ complex. Intriguingly, the β′ subunit docks closer to the α subunit near the bound C-terminus of the β subunit (Figure 3, black stars), with its own C-terminus reaching across to bind the α′ subunit (Figure 3, red stars). Conversely, the β subunit is tethered to the α subunit primarily via its flexible C-terminus, allowing it to freely approach the active site (Figure 3, top row). Thus, the preference for the α_2_ and β_2_ subunits to associate in an asymmetric manner produces a hinge-like region in the RNR complex. Importantly, 3D classification of the combined particle set resolved a new conformation that closely resembles the PCET conformation seen in *Ec* RNR (Figures 3C, S8A-B), both in compactness and subunit orientation. However, density for the β C-terminus is absent from the active site (Figure S8C), indicating that this class does not correspond exactly to the PCET process. Nonetheless, its appearance in the combined classification further supports the presence of shared conformations across multiple datasets and highlights the ability of cryo-EM to capture highly transient conformations. Furthermore, the resemblance in the relative α_2_/β_2_ positioning between this PCET-like class in *Bs* RNR and the *Ec* RNR structure suggests that the PCET process likely involves a conserved α_2_β_2_ conformation across class I RNRs.

**Figure 3.**
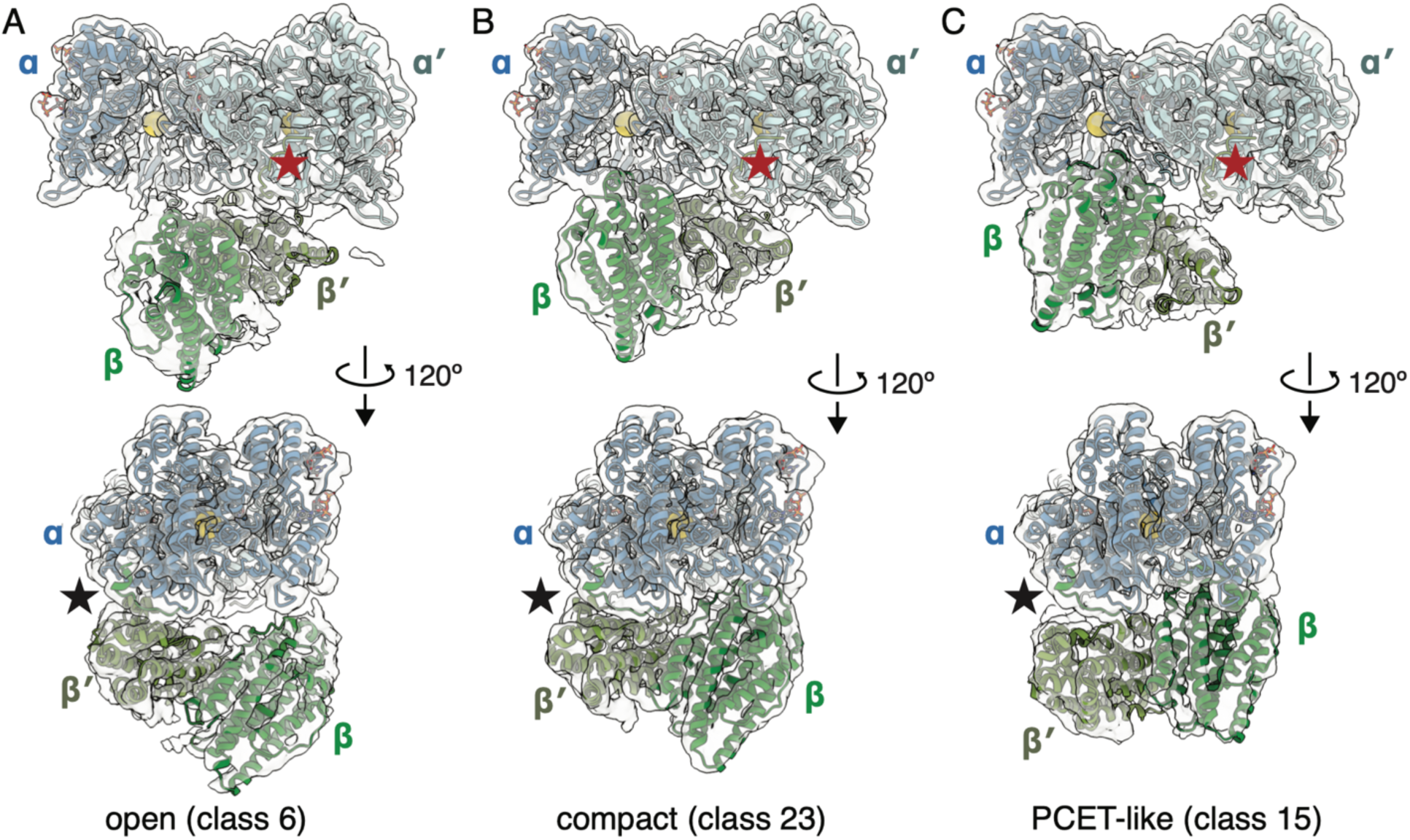
Representative 3D classes of the ɑ_2_β_2_ complex reveal asymmetric interactions between the dimeric subunits. Shown are (A) class 6 (open), (B) class 23 (compact), and (C) class 15 (PCET-like) from 3D classification of the combined particle set (Figure S7E). The C-termini of β and β′ are bound to ɑ and ɑ′, respectively. Although its own C-terminus is bound to ɑ′ (red star), the β′ subunit is anchored near the β C-terminus binding site on ɑ (black star). As a result, the β′ C-terminal tail is held stretched, and the ɑ′/β′ side is unable to close its inter-subunit distance. On the other hand, the β subunit is able to freely move with respect to the ɑ active site. *Top row*: a view of the β subunit approaching the active site of the ɑ subunit; the catalytic cysteine, C382, is shown as a large yellow sphere. *Bottom row*: a side view that shows the β′ subunit anchored near the β C-terminus binding site on ɑ (black star).

### Conformational landscapes of RNR

Because 3D classes appeared to represent a continuum of motions, we employed a deep-learning method, cryoDRGN^19,27^, to obtain conformational landscapes. Briefly, a cryoDRGN model was trained for each condition, after which 1000 volumes were generated from the latent space based on particle clustering (Methods Sections 6-7). Using custom code, physically interpretable conformational landscapes were constructed from the resulting volumes in the following manner (Methods Section 7, Code Availability). Atomic models of α_2_ and β_2_ were rigid-body fit into each volume, and the relative orientation of β_2_ from α_2_ was parametrized as a 7D coordinate representing the transformation (translation/rotation) between two coordinate systems defined on α_2_ and β_2_ individually. The 7D coordinates from all conditions were combined and embedded in 2D using DensMAP (density-preserving manifold approximation and projection^35^), from which a probability distribution was estimated to represent the combined conformational landscape of β_2_ orientations (Figure 4A). The joint embeddings of β_2_ orientations were then separated by condition to produce individual landscapes representing the distributions of α_2_β_2_ conformers within each ensemble (Figure 4B-E). We note that due to the particle filtering involved in the cryoDRGN training process and finite sampling of the latent space, low-population conformations may not be observed. However, processing was kept consistent between conditions such that comparisons can be made confidently. Importantly, discrete 3D classes fit well within these landscape representations, demonstrating consistency between conventional classification and deep-learning based analyses (Figure S9, Table S3). By mapping the resulting landscapes with metrics that describe the openness of β_2_ with respect to α_2_ (Figure S10A-B) and by mapping the coordinates of representative 3D classes (Figure 4A), we find that a trajectory from left to right correlates with a continuous transition from open to compact conformers, while the bottom-right quadrant contains PCET-like conformers. Notably, symmetric β_2_ orientations are essentially not observed (Figure S10C), and thus, the landscapes represent a continuum of *asymmetric* open and compact α_2_β_2_ conformers.

**Figure 4.**
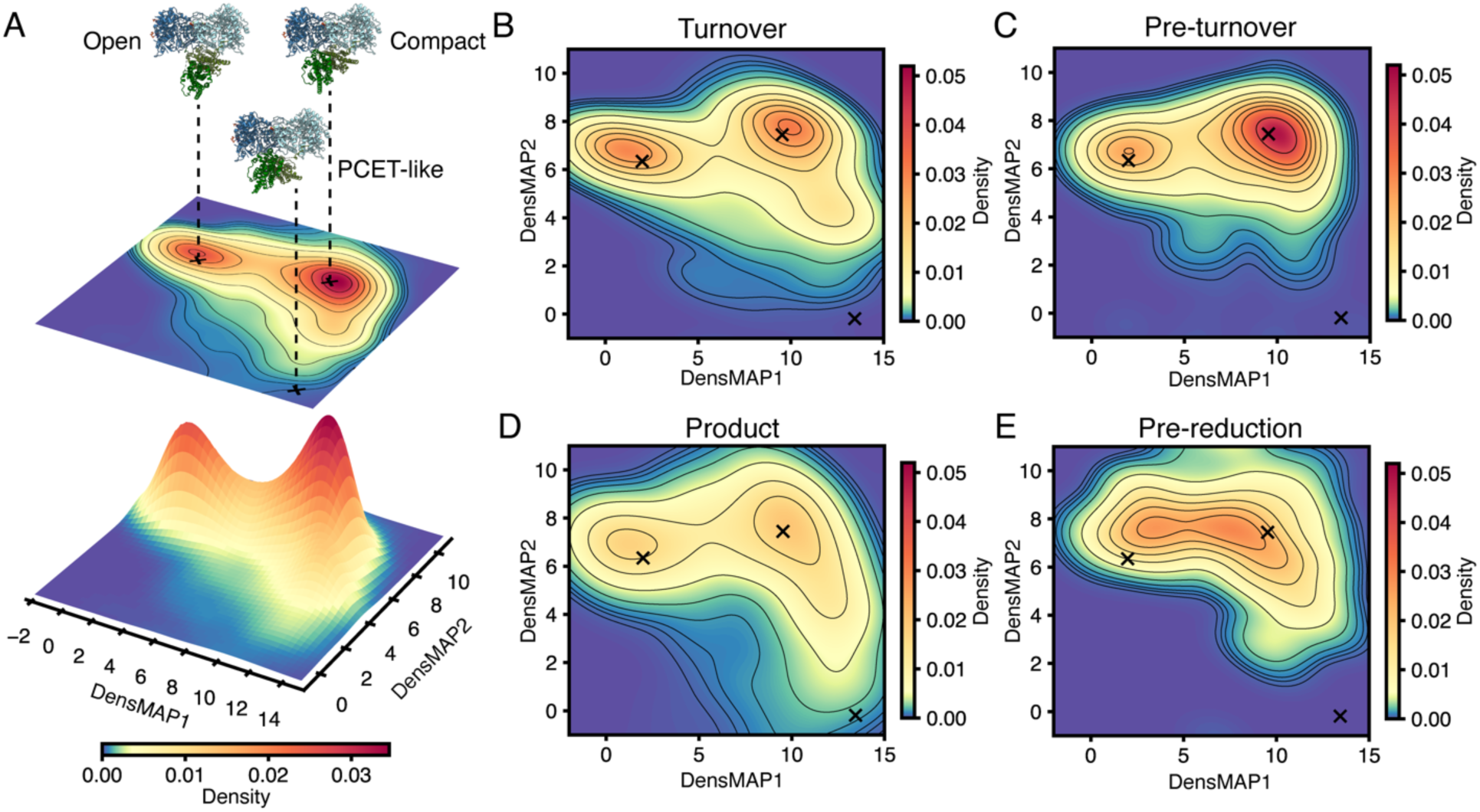
Conformational landscapes of *Bs* class Ib RNR highlight intrinsic dynamics and functionally relevant motions. Here, 7D coordinates representing the transformation (translation and rotation) between the α_2_ and β_2_ subunits are projected to 2D in a density-preserving representation (DensMAP), while the colorbar indicates population density (blue to red: low to high). (A) Joint conformational landscape obtained by combining coordinates from all conditions shown in panels B-E. Symmetric α_2_β_2_ conformations are essentially not observed (Figure S10C). Locations of representative open, compact, and PCET-like classes shown in Figure 3. (B-E) Conformational landscapes of individual conditions marked with the same three 3D classes as panel A. The enzyme exhibits intrinsic dynamics even before substrate is added (pre-turnover), while addition of substrate (turnover) or substrate analog (product) leads to an increased population of PCET-like conformations. A balance of open and compact populations is observed for the oxidized enzyme (pre-reduction), likely as a result of TrxA interactions.

The conformational landscapes from the individual conditions, while similar, are distinct from each other, indicating that RNR dynamics change throughout turnover (Figure 4B-E). Using the three marked coordinates corresponding to open, compact, and PCET-like 3D classes (Figure 3) as visual guides, we see that in the pre-turnover condition, compact conformers are enriched (Figure 4C). Nonetheless, the landscape is overall very broad, highlighting the existence of significant intrinsic motions in RNR even before the substrate is introduced. Compared with the pre-turnover landscape, the turnover and product landscapes share similar topologies and are spread more towards the lower right quadrant, where PCET-like conformers are found (Figure 4B,D). In the pre-reduction condition, the peaks corresponding to open and compact conformers are spaced closer together, suggesting that RNR undergoes more limited motion as the oxidized enzyme awaits the re-reduction process (Figure 4E). Thus, while the similarities of the landscapes are consistent with the observation of shared 3D classes, the differences highlight the roles of enzyme oxidation state and nucleotide occupancy in the active site in altering the conformational distributions to prime the enzyme for a specific stage of its turnover cycle.

Changes in the landscapes, particularly with respect to substrate and product, can be interpreted in the context of prior kinetic studies that have elucidated the rate constants for conformational and chemical steps in class I RNRs. These studies have shown that the rate of product formation during the catalysis half-cycle of class I RNRs is low (∼5-10 s^-1^) due to a slow conformational step that precedes PCET and catalysis, which are orders of magnitude faster (>150 s^-1^)^7,33,39^. Additionally, spectroscopic studies have shown that radical propagation is tightly regulated by the substrate/effector pair, with substrate being most critical in triggering this process^17,40–42^. The broadness of our landscapes provides an explanation for the rate-limiting conformational step in class I RNRs: the enzyme samples numerous α_2_β_2_ conformers and only rarely adopts the orientation required for PCET. The prevalence of shared α_2_β_2_ conformers across different conditions suggests that the intrinsic hinge-like motion of the enzyme complex (Figure 3) predominates over condition-specific effects. Despite the overall similarities, our landscapes reveal, for the first time, that the addition of substrate or product directly alters the conformational populations, such that a highly transient PCET conformation becomes accessible. In the trapped structure of *Ec* RNR^8^, the β C-terminus is observed folded into the active site, forming van der Waals contacts with the substrate and positioning an interfacial residue (Y307β in *Bs* numbering) for PCET (Figure S8D). The non-specific nature of this van der Waals interaction suggests that the β C-terminus may interact with the product in a similar manner. Such active-site stabilization by the substrate or product would explain why PCET-like conformers are enriched in the turnover and product landscapes. Interestingly, the largest population of PCET-like conformers is observed in the product condition (Figure 4D), suggesting that the product acts like a substrate analog in the reduced enzyme. The PCET conformation may have functional significance for the product-bound enzyme as reverse radical transfer must occur from the active site to the cofactor in the β subunit.

### Molecular mechanisms of TrxA binding

To investigate the molecular mechanisms of TrxA binding, we first estimated TrxA occupancy based on its map intensity relative to that of α_2_ within individual volumes from cryoDRGN landscape analysis (Methods Section 7). Histograms of TrxA occupancy on each side of the α_2_β_2_ complex are shown as violin plots in Figure 5A. Consistent with consensus reconstructions, we find that the median TrxA occupancy (middle, red dashed line in each distribution) is higher on the αʹ side in all conditions and that this value increases with the overall level of α_2_ oxidation (Figures 5A). Namely, the highest median TrxA occupancy is observed in the pre-reduction condition, while the largest variance in occupancy (spread of distribution) is observed in the turnover condition, which is the most heterogeneous in oxidation state. The preferential binding of TrxA to the αʹ side in more oxidized states is also evident when the TrxA occupancy is mapped onto the conformational landscapes (Figure S11). In addition, we observe greater TrxA occupancy on the left side of the conformational landscapes, indicating that TrxA binds more strongly to open α_2_β_2_ conformations than to compact conformations (Figure S11B).

**Figure 5.**
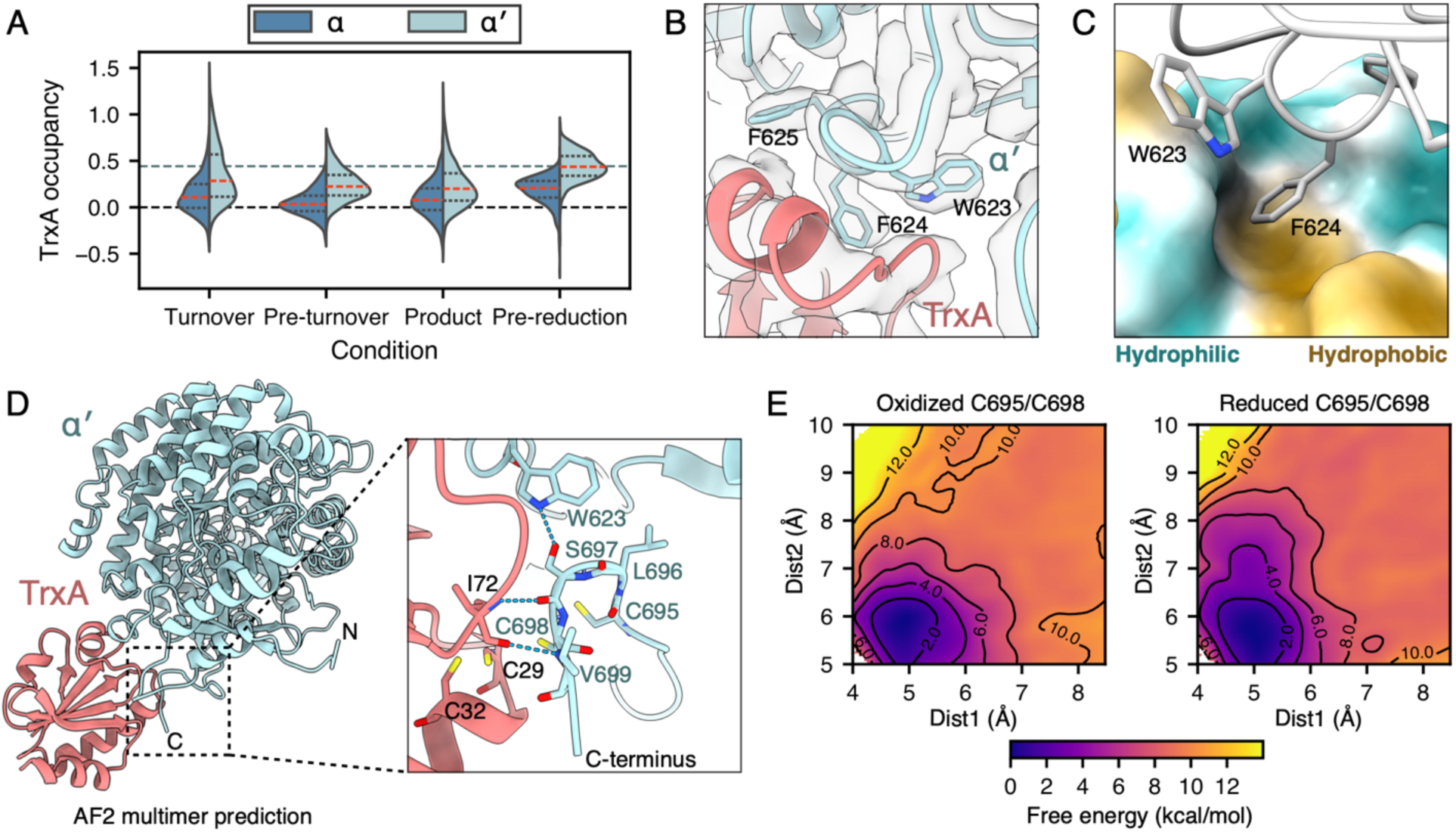
RNR-thioredoxin interactions favor the α′ side and depend on the oxidation state of α C-terminus. (A) Violin plot of TrxA occupancy obtained from landscape analysis, shown by condition and α_2_ protomer with quartiles within each distribution marked by dashed lines (median is colored in red). The highest TrxA occupancy is observed in the pre-reduction condition, while the largest variance is observed in the turnover condition, suggesting that TrxA preferentially binds oxidized RNR on the α′ side. (B) The interface between TrxA and the α′ subunit contains consecutive aromatic residues. Map shown is refined from particles with high TrxA occupancy from focused classification of pre-reduction dataset (Figure S1B). (C) F624α′ embeds in a hydrophobic pocket of TrxA, which is shown as surface colored by hydrophobicity. (D) AlphaFold2 multimer predictions of the TrxA-α′ complex depict the α′ C-terminal tail in a TrxA-bound conformation. (E) MD-generated 2D free energy landscapes of α′ C-terminal tail dissociation from TrxA-bound conformation show a steeper gradient when the C-terminal cysteine pair C695/C698 is oxidized, due to the rigidity provided by the disulfide bond. Two sampled distances are defined between the α′ C-terminal tail and its binding interface (Figure S13).

Focused classification of TrxA in the consensus refinement of the pre-reduction condition (Figure S1B), revealed a hydrophobic interface between TrxA and three aromatic residues on the surface of the α subunit: W623, F624, and F625 (Figures 5B-C). On the αʹ side of the α_2_β_2_ complex where TrxA is bound, we observe F624 embedded in a hydrophobic pocket of TrxA (Figure 5B-C), whereas on the α side, F624 adopts two rotamer conformations (Figure S12A), one of which conflicts with TrxA binding. MD simulations were conducted to determine whether the rotamer angle of F624 is correlated with the oxidation state of the active-site cysteine pair (C170/C409) or with the asymmetry of the α_2_β_2_ complex (Supplementary Methods Section S4, Figure S12B). The results suggest that F624 does not play a direct role in regulating the binding of TrxA. Instead, it likely contributes to the induced fit of the hydrophobic interface during TrxA binding.

We next examined whether TrxA binding is affected by the oxidation state of the cysteine pair (C695/C698) on the flexible α C-terminal tail. Previously, we were able to crystallographically trap this α-tail in the *Bs* RNR active site to show that substrate and tail binding are mutually exclusive^43^. Consistent with its function to dynamically move in and out of the substrate-binding pocket to reduce C170/C409, we do not observe density for the α-tail (residues 688-700) by cryo-EM (Figure S5). To obtain plausible tail conformations, we therefore used AlphaFold2 Multimer^44,45^ to predict the αʹ-TrxA complex, with all resulting predictions consistent with EM (root-mean-square deviation of 1.2 Å for the top prediction). Notably, in many of the predictions, the αʹ-tail adopts a conformation in which its two cysteines appear poised to react with the cysteine pair on TrxA (C29/C32), and other tail residues form several polar interactions with TrxA and one of the interfacial aromatic residues on the αʹ subunit, W623 (Figure 5D). To test whether the stability of this interaction is directly related to the oxidation state of C695/C698, we performed MD simulations on the reduced and oxidized states and obtained free energy landscapes along two distances defined between the αʹ-tail and its binding interface (Supplementary Methods Section S4, Figure S13) via 2D umbrella sampling (Figure 5E). We find that the disulfide bond in the oxidized αʹ-tail rigidifies the bound conformation, while the reduced αʹ-tail exhibits increased flexibility resulting in a less-steep pathway to dissociation. This kinetic trap results in a stronger interaction of the tail with TrxA when oxidized and thus leads to a higher observed occupancy. Overall, these results suggest that the binding affinity of TrxA is dependent on the oxidation state of the αʹ C-terminal cysteines.

### Conservation and diversity within class I RNRs

A question of interest is whether the dynamic interactions between α_2_, β_2_, and TrxA in *Bs* RNR are conserved across class I RNRs. Among the three biochemical classes of RNRs, the O_2_-dependent class I RNRs are unique in using a separate ferritin-like subunit (β_2_) to store the radical cofactor outside of the active site. Phylogenetically, class I RNRs comprise at least 11 clades (named by the gene prefix *nrd*) based on ɑ sequence,^3^ while biochemically, they can be classified by metal composition of the radical cofactor in β (of which 5 are currently known: Ia, Ib, Ic, Id, Ie)^5^. The class Ia cofactor (a Fe^III^-Fe^III^ tyrosyl radical) is predominant across class I clades, including the NrdAg clade, in which *Ec* RNR is a member. In contrast, *Bs* RNR uses a class Ib cofactor (a Mn^III^-Mn^III^ tyrosyl radical) and belongs to the largely bacterial NrdE clade, which is closely related to the NrdAg and NrdAh clades^3^ (Figure 6A). Based on the presence of a characteristic truncation of the N-terminal sequence, NrdE is thought to have emerged from this group via a gene duplication in bacteria^3,43^. Despite biochemical and phylogenetic differences, however, our cryo-EM results suggest that *Bs* RNR and *Ec* RNR likely share a similar ɑ_2_β_2_ conformation during PCET, which is further supported by the conservation of residues along the PCET pathway and structural features involved in stabilizing the PCET conformation^1–3,8,46^. On the other hand, whether the non-PCET conformational ensemble is conserved across class I RNRs is an important open question for future studies.

**Figure 6.**
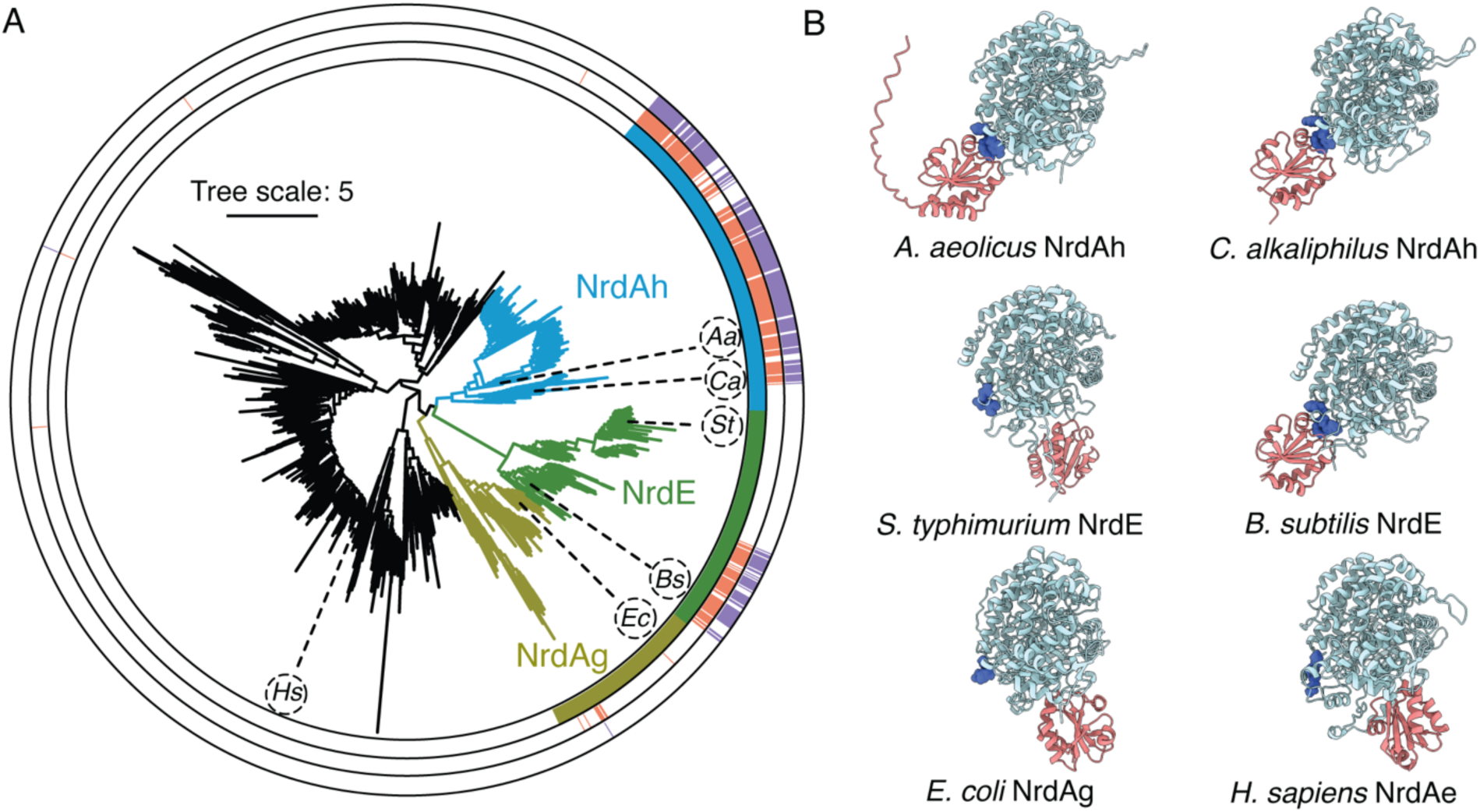
TrxA-binding interactions in class I RNRs. (A) Class I RNR phylogeny^2^ with NrdAg, NrdAh, and NrdE clades colored in gold, blue, and green, respectively, on both the tree and inner color strips. Many sequences in NrdAh and a subclade of NrdE contain an F624-equivalent aromatic residue (middle coral color strips) and at least one adjacent aromatic residue (outer purple color strips). Representative α-TrxA pairs subjected to AlphaFold2 multimer prediction are labeled on the tree by host organism: *Aa*, *Aquifex aeolicus*; *Bs*, *Bacillus subtilis*; *Ca*, *Chitinivibrio alkaliphilus*; *Ec*, *Escherichia coli*; *Hs*, *Homo sapiens*; *St*, *Salmonella typhimurium*. (B) The top AlphaFold2 multimer prediction (out of 25) for each of the α-TrxA pairs in panel A, with the α subunit shown in the same orientation. Residues corresponding to the WFF motif in *Bs* RNR (Figure 5B) are colored in dark blue and shown as spheres. Consistent with the presence of the interfacial aromatic residues, *Aa* and *Ca* RNRs are predicted to share the TrxA interaction seen in *Bs* RNR, while others are not predicted to share this trait.

Our effort to characterize *Bs* RNR under native-like conditions also led to the serendipitous capture of the RNR-TrxA complex for the first time. To investigate the prevalence of the observed TrxA-binding mode, we performed bioinformatic analyses on class I α sequences. A defining feature of this binding mode is the presence of three consecutive aromatic residues in the α subunit (WFF in *Bs* RNR) (Figure 5B). Sequence alignment of the 2022 class I α sequences from our previously published phylogeny^2^ revealed that 305 contain an aromatic residue at the F624-equivalent position, with 274 of these featuring at least one adjacent aromatic residue. Mapping these sequences onto the class I tree (Figure 6A) show that they cluster within the sister clades, NrdAh and NrdE. Notably, these aromatic residues are only present in one subclade of NrdE (which contains *Bs*) but not the other. This finding was corroborated by AlphaFold multimer predictions of TrxA-α pairs from organisms representing different class I clades (Figure 6B): *Aquifex aeolicus* (NrdAh), *Chitinivibrio alkaliphilus* (NrdAh), *E. coli* (NrdAg), human (NrdAe), and *Salmonella typhimurium* (NrdE subclade without interfacial aromatic residues). The resulting predictions suggest that only class I RNRs with the aromatic motif share the same TrxA interaction observed by cryo-EM in *Bs* RNR, while those from other organisms, including all eukaryotes (NrdAe), likely use alternative mechanisms for re-reduction. This finding aligns with prior studies suggesting that while the catalytic mechanism is universally conserved, the re-reduction mechanism is much more diverse, often involving different redox proteins across organisms^3,33,47–49^. This diversity in re-reduction mechanism could serve as the basis for future designs of antibiotics.

## Discussion

Dynamics have long been recognized to be critical for enzyme function, yet structural characterization of such motions has been challenging to achieve. Here, we performed advanced cryo-EM analyses to directly visualize the conformational landscapes of a class I RNR. We find that the active class I RNR undergoes significant continuous motions while maintaining asymmetry, explaining the abundance of biochemical evidence for half-site reactivity in both half-cycles of turnover^9–17^. Additionally, our landscapes provide direct structural evidence that conformations needed for PCET only become populated when the enzyme is provided with substrate or product, thus ensuring that radical transfer is regulated and reversible. While structural asymmetry does not necessarily mean that the two halves of the complex must alternate between the two halves of the turnover cycle, it provides an interlock that ensures that PCET only fires when the right constituents are in the active site and protects the α-tail from radical chemistry. As proxies for free energies, these landscapes also allow the visualization of structural dynamics associated with distinct steps detected by enzyme kinetics. Notably, it has long been known that PCET is preceded by a slow conformational step in class I RNRs^7,33,39^. The broadness of our landscapes provides direct visualization of the substantial conformational sampling that the enzyme undergoes in order to access transient conformations, such as that needed for PCET. Additionally, our results demonstrate that the asymmetric dynamics that physically separate the PCET and re-reduction processes in class I RNRs are further regulated by the presence of nucleotides in the active site as well as the oxidation state of the enzyme.

Based on our analyses, we propose a model for how the intrinsic dynamics of RNR leads to functional asymmetry (Figure 7). Both 3D classification and landscape analysis indicate that the class I RNR complex continuously samples asymmetric α_2_β_2_ conformations, maintaining an α/β (PCET-capable) side and an αʹ/βʹ (PCET-incapable) side. Although the overall dynamics of the α_2_β_2_ complex can be described as rigid-body motions of the α_2_ and β_2_ subunits, both subunits have flexible C-terminal tails that serve multiple functional roles. Both tails have been observed to interact with the other subunit. β-tail binding to the α subunit is essential for complex formation in class I RNRs^37,38^ (Figures 1A, 2B-C). The structure of the *Ec* RNR complex^8^ further suggests that the α-tail can bind the β subunit on the side engaged in PCET, and similar α-tail density is observed in some of our EM maps (Figure S14). Moreover, the flexible α and β tails play key roles in the two halves of turnover: the β-tail enables PCET by folding into the active site and contributing a critical interfacial residue^8^, while the α-tail shuttles between the active site and TrxA to fulfill the re-reduction process^43,50^. We thus propose that the asymmetry of the α_2_β_2_ complex regulates the two halves of the turnover cycle by creating different environments for the flexible α and β tails on each side, shifting their conformational ensembles and, in turn, modulating their functions.

**Figure 7.**
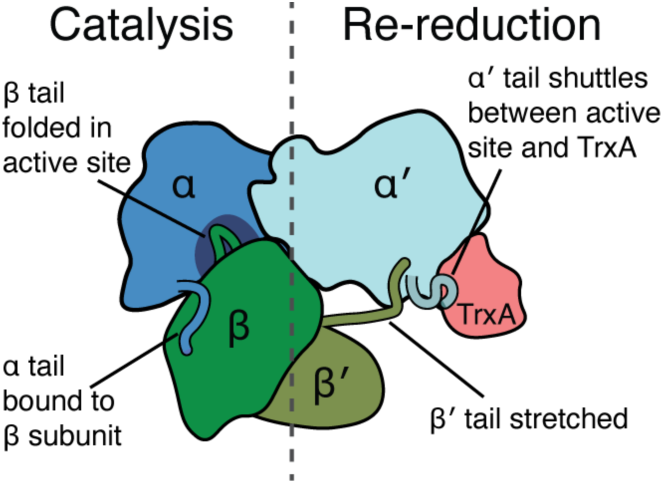
Proposed mechanism behind functional asymmetry in RNR. The α_2_β_2_ complex is highly dynamic but always asymmetric about the C2 axis of the α_2_ subunit (dashed line), leading to different interactions with the functional α and β C-terminal tails on each side. The β C-terminus binds the tail-binding cleft on both sides of the α_2_ subunit, whereas the α C-terminus can interact with the β_2_ subunit, the active site, or TrxA, depending on proximity. The α/β side of the complex can close into a PCET-capable conformation where the disordered region of the β tail can reach into the active site along with the substrate. In contrast, on the α′/β′ side of the complex, the β′ subunit is angled away from the α′ subunit, and the α tail is available for shuttling between the active site and TrxA. In order for the α/β and α′/β′ sides to switch, β_2_ will need to rotate to the opposite orientation, which likely involves temporary disengagement from α_2_.

In this framework, the global dynamics of the α_2_β_2_ complex are coupled to the local dynamics of the flexible C-terminal tails, which are additionally modulated by substrate binding to the active site and the oxidation state of the enzyme. Our cryo-EM results indicate that the α/β side of the α_2_β_2_ complex can dynamically open and close the inter-subunit distance to enable PCET. As the β subunit approaches the α subunit for PCET, the β-tail is able to fold and extend into the active site to make direct contact with the bound substrate, as captured in the *Ec* RNR structure^8^ (Figure S8D). The observed shift in our conformational landscapes towards the lower right quadrant under turnover and product conditions (Figures 3B,D) suggests that the closure of the α/β side into a PCET-competent conformation is facilitated by the presence of a nucleotide in the active site, which stabilizes the ordering of the β-tail. Entry of the β-tail into the active site also requires that it be unoccupied by the α-tail. Because the β subunit comes in close proximity to the α subunit on the α/β side of the complex, we propose that the affinity of the α-tail for the β subunit allows it to more readily vacate both the active site and the TrxA binding site. In contrast, on the αʹ/βʹ side of the complex, the βʹ subunit is tilted away from the αʹ subunit with its βʹ-tail stretched across the gap, preventing closure of the inter-subunit distance for PCET. However, this arrangement allows the αʹ-tail to freely move between the active site and TrxA. As our MD simulations suggest, oxidation of the αʹ-tail results in a more stable interaction with TrxA, leading to increased TrxA occupancy. In this model, it is notable that both the α and β tails achieve their most ordered structures when the correct conditions are met for re-reduction (oxidation of the tail cysteines) or catalysis (binding of nucleotides in the active site).

The dynamic protein-protein interactions observed in this study were also examined in an evolutionary context. Despite the diversity in sequence and cofactor usage within class I RNRs, our observation of a *Bs* RNR conformation closely resembling the PCET conformation of *Ec* RNR suggests that all class I RNRs share a common conformation for the radical transfer process. However, the highly variable α_2_-β_2_ affinity across different class I RNRs suggests that the conformational ensemble involved in the full turnover cycle may vary, potentially involving additional dissociation and association events for RNRs with low inter-subunit affinity. Additionally, our bioinformatic analyses revealed that the specific TrxA-α_2_ interaction observed in *Bs* RNR is only conserved in a small subset of bacterial class I RNRs, suggesting that the conformational ensembles in the re-reduction process may also vary across species. Investigating the conservation of these conformational ensembles across class I RNRs would thus be a compelling direction for future studies.

Although the conformational landscapes obtained here elucidate the overall distributions of β_2_ poses, many questions remain, such as the kinetics of β_2_ conformational switching as well as the dynamics of the α and β C-terminal tails. Future endeavors incorporating a temporal component, such as time-resolved cryo-EM and single-molecule experiments, may reveal the kinetic details of these motions, enabling direct comparison with biochemical data and validation of conformational landscapes obtained in this work. Furthermore, integrating computational approaches such as MD with cryo-EM data presents a promising approach for overcoming the limitations of conventional data processing. In particular, methods proposing to extract information from raw particle images, such as MD ensemble reweighting^51^, may provide an avenue for overcoming the loss of intrinsically disordered regions, such as the α and β C-terminal tails, during conventional cryo-EM averaging. Nevertheless, our findings underscore the critical role of conformational dynamics in regulating enzyme mechanisms, and our approach establishes a foundation for future comparative studies that examine the reproducibility of landscape analysis and the conservation of dynamics across species.

## Methods

### 1. Protein Expression, Purification, and Biochemical Characterization

The following proteins from *B. subtilis* were prepared as described previously^33,52,53^: tagless NrdE (class Ib RNR ɑ subunit), His_6_-tagged Mn-reconstituted NrdF (class Ib RNR β subunit), tagless NrdI (flavodoxin), His_6_-tagged TrxA (thioredoxin), and His_6_-tagged TrxB (NADPH-dependent thioredoxin reductase). The radical content of NrdF was determined to be 1.23 Y•/dimer, consistent with previously reported values^33,52^.

Enzyme activity was assayed spectroscopically as previously described^52^ in RNR storage buffer (50 mM HEPES, 150 mM NaCl, 15 mM MgCl_2_, 5% (w/v) glycerol, pH 7.6). A reaction mixture containing 1 µM NrdE, 1 µM NrdF, 0.25 mM TTP, 1 mM ATP, 0.5 mM GDP, 40 µM TrxA, 0.4 µM TrxB, 0.2 mM NADPH yielded an activity of 225 ± 28 U (mg α_2_β_2_)^-1^, comparable to the previously reported *V_max_* of 220 ± 3 U (mg α_2_β_2_)^-1^ with the GDP-TTP substrate-effector pair^52^. All protein concentrations in this work are given as monomer concentrations. Detailed methods are provided in Supplementary Methods (Section S1).

### 2. Cryo-EM Sample Preparation and Data Collection

Samples for cryo-EM were designed to mimic assay conditions, using TTP as the specificity effector and ATP as the activity effector. All samples contained 5 µM NrdE, 5 µM NrdF, 30 µM TrxA, 0.25 mM TTP, and 1 mM ATP in RNR storage buffer. Substrate (GDP), product (dGDP), TrxB, and NADPH concentrations were varied to capture four different stages of enzymatic turnover.

#### Turnover condition

All ingredients but the substrate were pre-mixed in one tube (40 μL), while a 2.5 mM GDP stock was kept separately. Both solutions were equilibrated at room temperature for 5 min before the reaction was initiated by adding 10 μL of GDP stock through repetitive pipetting to obtain a reaction mixture with excess GDP (0.5 mM) as well as high levels of TrxB (1 µM) and NADPH (1 mM) to ensure sufficient reduction speed. The sample was frozen within 20 s, with <10% of GDP turnover based on the enzyme activity. ***Pre-turnover condition:*** A solution was prepared with the same composition as the turnover condition, except with 0.4 µM TrxB and no GDP. ***Product condition:*** A solution was prepared with the same composition as the turnover condition, except with 0.4 µM TrxB and 0.5 mM dGDP instead of GDP. ***Pre-reduction condition:*** A solution was prepared with the same composition as the turnover condition, except with low TrxB (0.4 µM) and limiting NADPH (0.1 mM). The sample was frozen after a 3-min incubation at room temperature, with NADPH depleted but <20% of GDP turnover. This condition represents a state where the RNR ɑ subunit (active site and C-terminus) and TrxA/TrxB are oxidized and awaiting re-reduction by NADPH before another round of turnover can begin.

#### Grid preparation and data collection

Grids were frozen using an FEI Vitrobot Mark IV and imaged on a Talos Arctica (Thermo Fisher) at the Cornell Center for Materials Research (CCMR) with a Gatan K3 direct electron detector and BioQuantum energy filter. Additional details are in Supplementary Methods (Section S2). Cryo-EM datasets in this work are interpreted under the assumption that the conformational ensemble is preserved through the rapid cryo-cooling of grids, based on recent computational work indicating that energy barriers >10 kJ mol^-1^ (2.4 kcal mol^-1^), such as those between sidechain rotamer conformations, are unlikely to be crossed during this process^54^.

### 3. Cryo-EM Data Processing

Datasets for all four conditions were processed individually through similar workflows in cryoSPARC 4^55^ and RELION 4^56^. Motion correction was performed using RELION’s MotionCorr2^57^ with 7 x 5 patches (pre-turnover and pre-reduction datasets) or cryoSPARC patch motion correction (turnover and product datasets). Dose-weighted micrographs motion-corrected in RELION were imported into cryoSPARC. Patch CTF estimation was performed in cryoSPARC, and micrographs were filtered by a 5-Å CTF fit resolution cutoff and visual inspection.

For each dataset, particles were picked with Topaz^58^ using a pretrained ResNet16 model, extracted with a box size of 256 x 256 pixels, Fourier cropped to 128 x 128 pixels, and subjected to 2D classification. Classes containing secondary structural details were kept and further filtered with a class probability cutoff of 0.99. The top 400 - 600 micrographs with the most remaining particles were used to train a Topaz ResNet8 model initialized from pretrained weights, which was used for final particle picking. The resulting particles were again extracted with a size of 256 x 256 pixels, Fourier cropped to 128 x 128 pixels, and subjected to 2D classification to further exclude low-quality classes. *Ab initio* reconstruction and homogeneous refinement were performed on kept particles, and duplicate particles with separation distance <40 Å were removed. This process yielded 1.011M particles from 3258 micrographs (turnover), 893K particles from 2532 micrographs (pre-turnover), 991K particles from 2777 micrographs (product), and 552K particles from 3785 micrographs (pre-reduction) (Table S1).

Micrographs initially motion-corrected in cryoSPARC were motion-corrected again in RELION 4. Particle coordinates from cryoSPARC were converted to RELION star format with pyem^59^. Particles were re-extracted from the motion-corrected micrographs in RELION with a box size of 256 x 256 pixels and subjected to 3D auto refine and CTF parameter refinement (anisotropic magnification, beam tilt, trefoil, spherical aberration, per-micrograph astigmatism, and per-particle defocus), with particles split into 9 optics groups based on the 3 x 3 multishot position of their originating micrographs during collection. Bayesian polishing parameters were then trained with 8000 particles, and two iterations of Bayesian polishing and CTF refinement were performed to obtain polished shiny particles. The shiny particles were imported into cryoSPARC for one to two rounds of multi-class *ab initio* reconstruction and heterogeneous refinements into five classes. Particles from classes with poor β_2_ density were excluded. This process led to particle sets consisting of 597K (turnover), 667K (pre-turnover), 455K (product), and 368K (pre-reduction) particles, which were used for 3D classification in RELION 4 (Figures S15, S17, S19, S21: red path).

For each dataset, an additional round of heterogeneous refinement was performed to produce a slightly more optimized particle set for cryoDRGN analyses. True pixel sizes were estimated through comparison of the spherical aberration obtained from CTF parameter refinement (with a single optics group) to the true value of the instrument (2.7 mm), resulting in the following values: 1.017 Å (turnover), 1.022 Å (pre-turnover), 1.017 Å (product), and 1.014 Å (pre-reduction). CTF parameters were re-estimated after pixel-size correction, and homogeneous refinement and non-uniform refinement^60^ were finally performed in cryoSPARC to generate the consensus map for each condition. The sharpened map from this consensus refinement (Figures S15, S17, S19, S21: purple triangle) was used to build and refine atomic models of the α_2_ bound with the β C-terminal tail for each condition (Table S1). Model building and refinement were performed in Phenix^61^ and Coot^62^ (Supplementary Methods Section S3). This process resulted in particle sets consisting of 596K (turnover), 666K (pre-turnover), 505K (product), and 361K (pre-reduction) particles (Figures S15, S17, S19, S21: blue path).

Focused classification of TrxA was performed in cryoSPARC on the 552K pre-reduction particle set (Figure S21) with a mask created in ChimeraX^63^ by using molmap to generate a 20-Å resolution map of a TrxA AlphaFold2^44^ model docked in the EM map and thresholding at 0.1. Two rounds of 3D classification into 10 classes were performed, and particles from classes with high TrxA occupancy were combined (171K particles). Homogeneous refinement and nonuniform refinement were performed, and the resulting half-maps were sharpened with deepEMhancer^64^ using the tightTarget model (Figure S22B, Table S2).

### 4. Analysis of Conformational Changes via 3D Classification

Particle sets were imported into RELION 4 (Figures S15, S17, S19, S21: red path), and discrete 3D classification was performed individually on each condition. 3D auto refine and CTF parameter refinement were first performed to produce a consensus reconstruction. Particle subtraction was then performed with a spherical mask (40 Å radius) encompassing the β_2_ region, removing α_2_ signal from particles. 3D classification without alignment was finally performed with 20 classes, focused on the β_2_ region within the spherical mask. To address pseudosymmetry of the class I RNR complex (the major signal for alignment comes from the larger, symmetric α_2_ while the asymmetrically posed β_2_ contributes less to alignment), an additional consensus reconstruction was generated using a reconstructed 3D class with a well-resolved β_2_ subunit in a compact conformation as a reference map. The consensus map resulting from this step was also used in rigid-body docking that yielded the consensus full-complex models of RNR for each condition (Figures S15, S17, S19, S21: green square, Table S2, Supplementary Methods Section S3). Particle subtraction and 3D classification were performed on the new consensus reconstruction in the same fashion as before. Subtracted particles from classes with β_2_ density consistent with the β_2_ model, featuring clear secondary structures (largely helical), were reverted to original particles and reconstructed individually for each class. This process yielded 8 classes for the turnover condition, 5 classes for the pre-turnover condition, 8 classes for the product condition, and 3 classes for the pre-reduction condition (Figures S15, S17, S19, S21, Table S2). Full-complex models were built for every 3D class (detailed in Supplementary Methods Section S3).

### 5. Combined Refinement of All Conditions Prior to CryoDRGN

Particles from the consensus refinements were collapsed into a single optics group per condition, and CTF parameters were re-estimated with only per-particle defocus optimization in cryoSPARC for cryoDRGN analysis (which does not support multiple optics groups or higher order aberration corrections). To align particle poses across conditions with differing pixel sizes, particles were imported to RELION 4 with each condition consisting of one optics group. Particles from all four conditions were combined and jointly refined, and a consensus reconstruction was obtained using a compact class from the pre-reduction condition as the reference map (Figure S23). Focused 3D classification without alignment was performed on the all-condition combined refinement with particle subtraction of the α_2_ signal (as described above) to generate 50 classes. Particles from classes with β_2_ aligning to the opposite direction or with density unresembling β_2_ were discarded (16 classes out of 50 classes, 442K particles out of 2.127M particles). Further 3D classification of discarded particles did not yield better resolved classes. Subtracted particles from kept classes were then reverted to original ones. 13 classes with well-resolved β_2_ density were reconstructed individually (Figure S23 black boxes, Table S2). Full-complex models were built for every 3D class (detailed in Supplementary Methods Section S3). Particles from all kept classes were then split back into the four conditions, containing 476K (turnover), 536K (pre-turnover), 406K (product), and 287K (pre-reduction) particles for cryoDRGN analysis.

### 6. Training CryoDRGN Models and Particle Filtering

CryoDRGN models were trained on each condition individually in cryoDGRN version 2.3. Particles were first downsampled to a box size of 128 x 128 (∼2.04 Å pixel size) and converted to Fourier space representations. An 8D latent variable model with 1024 x 3 encoder and decoder dimensions was trained on each dataset for 50 epochs. The latent dimension (8) and the multilayer perceptron (MLP) dimension (1024 x 3) were chosen based on previous applications of cryoDRGN^19,65^. Smaller MLP dimensions were unable to model the full range of motion of β_2_ (not shown).

Particle filtering was performed during cryoDRGN training in several stages. In the first round of filtering, particles were filtered by UMAP representation, as recommended by cryoDRGN developers. UMAP embeddings of the latent space representation were inspected, and particles in junk clusters were removed (Figures S25-S28, blue clusters in top right plot). A second latent variable model was then trained on the remaining particles with the same parameters as the first, and *k*-means clustering was applied in latent space to generate 50 volumes representing each cluster. Particles from clusters with visually poor β_2_ density were excluded. Additionally, *z*-score outliers were filtered following guidance from cryoDRGN developers. A third latent variable model was then trained on the kept particles, from which a *k*-means clustering with *k*=1000 was performed to generate 1000 cluster volumes.

Of the 1000 cluster volumes, those with poor β_2_ density were identified via an automated procedure. The refined pre-turnover α_2_ model and a β_2_ crystal structure (PDB: 4dr0^36^, residues before Thr18 removed) were rigid-body fit into each volume using a custom ChimeraX script. Starting configurations for α_2_ and β_2_ were taken from the all-condition combined refinement. α_2_ was docked first, and its expected map was generated at Nyquist resolution with molmap and subtracted from the cluster volume. The subtracted volume was thresholded by a level computed as the mean plus 0.25 times the difference between maximum and mean intensity of all voxels to minimize the influence of “hot voxels” observed in certain cluster volumes. The β_2_ model was then docked in the subtracted and thresholded map. Binary masks for α_2_ and β_2_ were generated with molmap (20 Å resolution, threshold 0.1). The β_2_ “occupancy” was calculated in a similar fashion to previous work^27^, where the ratio of average voxel intensities within the β_2_ and α_2_ masks was compared to the expected ratio for full occupancy from a volume generated with molmap at Nyquist-limit resolution (∼4.08 Å). For each volume, the number of connected voxels in all overlapping regions between α_2_ and β_2_ masks was computed, and the largest value among those was used as an “overlap score”. After visually inspecting various cutoff values, a β_2_ “occupancy” cutoff of 0.65 and an “overlap score” cutoff of 150 was found to reasonably filter clusters with poor β_2_ density from the 1000 cluster volumes. Principal component analysis (PCA) of the latent space representation was used to further filter particles with poor β_2_ density for two datasets, which displayed principal component (PC) endpoint volumes with poor β_2_ density. For the turnover condition, particles with PC2 > 0.25 were excluded, while for the pre-reduction condition, particles with PC1 < −0.25 were excluded. Following this step, particle sets contained 255K (turnover), 286K (pre-turnover), 246K (product), and 128K (pre-reduction) particles. A final latent variable model was then trained on each dataset, yielding the cryoDRGN models used for landscape analysis. Volumes along the first two PCs of the final cryoDRGN model for each condition are animated as movies (Movies S1-S4).

### 7. Landscape Analysis of Multiple Datasets

For each dataset, *k*-means clustering was applied to the latent space of the final cryoDRGN model to generate 1000 cluster volumes. Cluster volumes were filtered as described above, using a β_2_ “occupancy” cutoff of 0.7 and a “overlap score” cutoff of 150, to produce the final datasets for landscape analyses: 863 clusters consisting of 225K particles for the turnover condition, 727 clusters consisting of 212K particles for the pre-turnover condition, 801 clusters consisting of 199K particles for the product condition, and 858 clusters consisting of 113K particles for the pre-reduction condition.

#### Landscape construction and visualization

To map cluster volumes from different datasets in a common landscape representation, α_2_ and β_2_ models were fit to each volume, and individual coordinate systems were defined for each subunit (Figure S29 and detailed in a following section). The orientation of β_2_ relative to α_2_ was then parametrized as the rigid-body transformation between the two coordinate systems with a 7D coordinate, comprising the translation vector (3D) between the α_2_ and β_2_ centers of mass (COMs) and the quaternion (4D) representing the rotation.

DensMAP^35^ was used to embed the 7D coordinates into a 2D representation. To quantify the similarity of any two β_2_ conformations in the embedding process, we defined the distance metric as the sum of distance between the two β_2_ COMs (in Angstroms) and the angle of rotation between the two quaternions (in degrees). An n_neighbors value of 50 was used. The 7D coordinates from all four datasets, as well as those calculated from 3D classes obtained in RELION, were combined to produce a single joint embedding (Figure S9A, Table S3). To test the stability of the resulting embeddings, the DensMAP calculation was performed with 100 different random seeds (Figure S30). While the majority of the embedding was reproducible, the embedding around the PCET-like region showed slight variations due to the smaller population. We thus chose a representative embedding (random state 64) for further analysis, based on the known similarity between individual classes in the PCET-like region.

The embedded coordinates from the joint embedding (random state 64) were then separated by condition to produce the 2D representation for each condition (Figure S9B-C). Gaussian kernel density estimation was used to estimate the 2D probability distribution of each conformational landscape, with the number of particles in each cluster used as weights. The smoothing bandwidth used during estimation was calculated with Scott’s Rule^66^ for the combined embedding, and a fixed value of 0.328 (equivalent to 800 data points with Scott’s Rule) for the individual conditions.

#### Validation of clustering resolution

To verify that a *k*-means clustering with *k*=1000 sufficiently samples the latent space, we repeated the landscape analysis from trained cryoDRGN models with a smaller number of clusters (*k*=500). The resulting representation (Figure S31) is consistent in terms of topology, 3D class locations, and the estimated landscape, confirming the sampling sufficiency at *k*=1000.

#### Definitions of subunit coordinate systems

Rigid-body models of the ɑ_2_β_2_ complex were produced for every cluster volume by fitting the refined pre-turnover α_2_ model (chains A-B) and a β_2_ crystal structure (PDB: 4dr0^36^, chains A-B renamed to C-D), as shown in Figure S29A.

The first coordinate system was centered on the COM of α_2_, with the *y*-axis defined as the unit vector along the C2 axis of α_2_ (computed as the intersecting line between the plane formed by the residue-180 Cα atoms in each monomer and the dimer COM, and the plane formed by the residue-190 Cα atoms in each monomer and the dimer COM). The *z*-axis was defined as the cross product between the *y*-axis and the vector between the dimer COM and the residue-308 Cα atom in chain A. The *x*-axis was then defined as the cross product of the *y*- and *z*-axes.

The second coordinate system was centered on the COM of β_2_, with the *yʹ*-axis defined as the unit vector along the C2 axis of the β_2_ (computed as the intersecting line between the plane formed by the residue-111 Cα atoms in each monomer and the dimer COM, and the plane formed by the residue 215-Cα atoms in each monomer and the dimer COM). The *zʹ*-axis was the cross product between the *yʹ*-axis and the vector between dimer COM and the residue-179 Cα atom in chain D. The *xʹ*-axis was the cross product of the *yʹ*- and *zʹ*-axes.

#### Analysis of ɑ_2_β_2_ asymmetry and openness

For each modeled ɑ_2_β_2_ complex, the distance between the Cα atoms of two residues involved in inter-subunit radical transfer, Y684 in chain A in α_2_ and W30 in chain C in β_2_, was calculated. Euler angles *ψ, θ, φ* corresponding to extrinsic rotation with about the *x*-*y*-*z* axes (Figure S29B) were also calculated. Notably, Euler angle *φ* is correlated with the “openness” of β_2_ relative to α_2_, while Euler angle *ψ* is correlated with the asymmetric posing of β_2_.

#### Analysis of TrxA occupancy

TrxA occupancy in each cluster volume was calculated in a manner similar to the β_2_ occupancy calculation described above. Binary masks for TrxA on each side were generated from a docked TrxA model using molmap (20 Å resolution, threshold 0.1). Occupancy values were calculated as the ratio of average voxel intensities within the TrxA and α_2_ masks divided by the expected ratio for full occupancy from a volume generated at Nyquist-limit resolution (∼4.08 Å) with molmap.

### 8. All-Atom MD Simulations and Umbrella Sampling

MD simulations were performed using AmberTools23^67^ and Amber22^68^. We used the ff19SB force field^69^ with OPC water^70^ in our simulations as it the most accurate Amber protein force field that was shown to improve amino-acid dependent properties. We used Li-Merz 12-6 ions model^71,72^ to parametrize unbound ions and used MCPB.py^73^ and GAMESS-US^74^ to parametrize the dimanganese center in β_2_ as bound ions following previous MD work on class I RNRs^75^. We parameterize the TTP ligand following previous works^75^. Details of parameterization, equilibration, and umbrella sampling can be found in Supplementary Methods (Section S4).

## Supporting information

Movie_S1_turnover

Movie_S2_preturnover

Movie_S3_product

Movie_S4_prereduction

## Data Availability

Cryo-EM maps have been deposited to the Electron Microscopy Data Bank with accession codes EMD-44947, EMD-44985, EMD-44991, EMD-44992, EMD-44995, EMD-44999, EMD-45000, EMD-45004, EMD-45010, EMD-45011, EMD-45014, EMD-45015, EMD-45016, EMD-45017, EMD-45018, EMD-45019, EMD-45020, EMD-45021, EMD-45023, EMD-45024, EMD-45026, EMD-45029, EMD-45030, EMD-45031, EMD-45037, EMD-45044, EMD-45045, EMD-45046, EMD-45047, EMD-45048, EMD-45049, EMD-45051, EMD-45052, EMD-45053, EMD-45054, EMD-45057, EMD-45061, EMD-45064, EMD-45065, EMD-45066, EMD-45067, EMD-45068, EMD-45069, EMD-45070, EMD-45071, EMD-45072. The associated atomic models have been deposited to the Protein Data Bank with accession codes 9bw3, 9bwx, 9bx2, 9bx3, 9bx6, 9bx8, 9bx9, 9bxc, 9bxs, 9bxt, 9bxx, 9bxz, 9by0, 9by1, 9by2, 9by3, 9by7, 9by8, 9by9, 9bya, 9byc, 9byd, 9byg, 9byh, 9byl, 9byt, 9byv, 9byw, 9byx, 9byy, 9byz, 9bz2, 9bz3, 9bz5, 9bz6, 9bz9, 9bza, 9bzd, 9bze, 9bzf, 9bzh, 9bzi, 9bzj, 9bzk, 9bzm, 9bzo. Final cryoDRGN model weights for the four conditions used in landscape analysis as well as configuration and result files from MD simulations have been deposited to Zenodo with accession code 14837414 (https://doi.org/10.5281/zenodo.14837414). Raw EM data for the four conditions have been deposited to EMPIAR with accession codes EMPIAR-12420, EMPIAR-12421, EMPIAR-12422, EMPIAR-12423.

## Code Availability

Custom code used to generate the densMAP embeddings and conformational landscapes are available as Chimera-X scripts and Jupyter notebooks on GitHub (https://github.com/ando-lab/2024-xu-scripts)^76^.

## Acknowledgments

The authors are grateful to Prof. JoAnne Stubbe (MIT) for sharing plasmids of *Bs* RNR, NrdI, TrxA, and TrxB and many insightful conversations over the years. We are also grateful to Drs. Maxwell Watkins and Steve Meisburger for discussions during an earlier phase of this project and to Prof. Brian Crane (Cornell) for critical reading of this manuscript. Cryo-EM data in this work were collected at the Cornell Center for Materials Research (CCMR), which is supported by the National Science Foundation (NSF) under award DMR-1829070. This work also made use of the National Center for CryoEM Access and Training (NCCAT) and the Simons Electron Microscopy Center located at the New York Structural Biology Center, supported by the NIH Common Fund Transformative High Resolution Cryo-Electron Microscopy program (U24 GM129539) and by grants from the Simons Foundation (SF349247) and NY State Assembly. This work was supported by National Science Foundation (NSF) grant MCB-1942668 and startup funds from Cornell University to N. A.

## Author Contributions

N. A. conceived of the project, oversaw and supervised the research, analyzed data, and acquired funding. D. X. performed sample preparation, biochemical characterization, cryo-EM characterization, computation, bioinformatics, and analyses described in this work. W. C. T. prepared *Bs* NrdE and performed analysis of a preliminary EM dataset collected on Bs RNR with chemical reductant, and A. A. B. prepared Bs TrxB. D. X. and N.A. wrote the manuscript.

