## Supplementary_Information for "Conformational Landscapes of a Class I Ribonucleotide Reductase Complex during Turnover Reveal Intrinsic Dynamics and Asymmetry"

**This PDF file includes:**

Supplementary Methods

Figures S1 to S34

Tables S1 to S4

References

**Other Supplementary Materials for this manuscript include the following:**

Movies S1 to S4

### Supplementary Methods

#### S1. Protein Expression, Purification and Characterization

NrdE (RNR α subunit), Mn-reconstituted NrdF (RNR β subunit), NrdI (flavodoxin), TrxA (thioredoxin), and TrxB (NADPH-dependent thioredoxin reductase) from *B. subtilis* (*Bs*) were prepared according to published protocols[1–3] with key steps and modifications described below. Plasmids containing genes for N-$His_6$-Smt3-tagged NrdE, N-$His_6$-tagged NrdF, N-$His_6$-Smt3-tagged NrdI, N-$His_6$-tagged TrxA, and N-$His_6$-tagged TrxB were generously provided by JoAnne Stubbe (MIT). All proteins were expressed in an *E. coli* BL21 derivative (T7 Express, NEB) using Luria Broth (Miller) medium, unless described otherwise. Nickel and cobalt affinity chromatography steps described below were performed with HisPur Ni-NTA resin (ThermoFisher) and TALON cobalt affinity resin (Takara), respectively. All protein concentrations below are given as monomer concentrations.

***Bs* RNR α subunit.** N-terminally $His_6$-Smt3-tagged NrdE was purified with a nickel affinity column, followed by anion exchange chromatography (HiPrep Q FF 16/10, Cytiva). The $His_6$-Smt3 tag was cleaved using SUMO protease and separated from tagless NrdE (having only its native sequence) using a second nickel affinity column. We note that *Bs* NrdE, purified at this stage, is partially inhibited by endogenously bound deoxyadenosine 5′-monophosphate (dAMP)[3], but its effect is reversed by addition of purine nucleotides, such as adenosine 5′-triphosphate (ATP) and guanosine 5′-diphosphate (GDP)[4], which are present in all of the experiments described in this study. Finally, tagless NrdE was exchanged into RNR storage buffer (50 mM HEPES, 150 mM NaCl, 15 mM $MgCl_2$, 5% (w/v) glycerol, pH 7.6) supplemented with 1 mM tris(2-carboxyethyl)phosphine) (TCEP).

***Bs* NrdI.** N-terminally $His_6$-Smt3-tagged NrdI, used in the reconstitution of NrdF, was purified with a cobalt affinity column, and the tag was cleaved and separated in the same manner as NrdE to produce tagless NrdI (having only its native sequence). Cofactor reconstitution was then performed by incubating ~15 μM tagless

NrdI with ~800 μM flavin mononucleotide (FMN) for 4-6 hours before exchanging into reconstitution buffer (50 mM HEPES, 5% glycerol, pH 7.6) to remove free FMN.

***Bs* RNR β subunit.** N-terminally $His_6$-tagged apo-NrdF was expressed with 1,10-phenanthroline added to the cell culture at a final concentration of 0.1 mM 20 min before induction with isopropyl β-d-1-thiogalactopyranoside (IPTG). $His_6$-tagged apo-NrdF was then purified with a cobalt affinity column and exchanged into reconstitution buffer. Metal reconstitution of NrdF was performed in a Coy anaerobic chamber under a $N_2/H_2$ atmosphere (95%/5%) at room temperature. Deoxygenated reconstitution buffer was used to prepare anoxic $MnCl_2$ and $Na_2S_2O_4$ solutions inside the chamber, and apo-NrdF solution was allowed to equilibrate with the chamber atmosphere for an hour. Tagless NrdI was fully reduced to its hydroquinone form with $Na_2S_2O_4$ solution, and a 2-fold molar excess of $MnCl_2$ was mixed with the apo-NrdF solution. Equimolar apo-NrdF and NrdI (quantified by FMN concentration) were used. Both solutions were incubated for 20 min, before the two solutions were mixed and taken out of the Coy chamber. Oxygen-saturated reconstitution buffer was then added at ~2-3 times the volume to the mixture to form the dimanganese tyrosyl radical ($Mn(III)_2$–Y•) cofactor. Mn-reconstituted NrdF (Mn-NrdF) was purified from mismetallated NrdF and NrdI by high-resolution strong anion exchange (Mono Q 10/100, Cytiva) before exchanging into RNR storage buffer. The radical content of Mn-NrdF was determined to be 1.23 Y•/dimer (comparable to previously reported values[1,2]) with electron paramagnetic resonance (EPR) spectroscopy by comparison to a Fe-reconstituted NrdF sample with known radical content and a 2,2,6,6-tetramethyl-1-piperidinyloxy (TEMPO) standard.

***Bs* RNR reducing system.** N-terminally $His_6$-tagged TrxA was purified with a cobalt column, followed by size exclusion chromatography (HiLoad 16/600 Superdex 200 pg, Cytiva) in buffer containing 50 mM sodium phosphate, 150 mM NaCl, pH 7.6. Pure fractions were combined and exchanged into RNR storage buffer. N-terminally $His_6$-tagged TrxB was expressed, purified, and assayed following published methods[1,2] with the following modifications. Protein was expressed in BL21-derived C43 *E. coli* cells (OverExpress

C43(DE3) cells, Sigma) and induced with 0.4 mM IPTG for 6 hours. Harvested cells were lysed in 50 mM sodium phosphate, 300 mM NaCl, 10 mM imidazole, and 5% (v/v) glycerol, pH 7.6, with 2 mM flavin adenine dinucleotide (FAD), 2 U $mL^{-1}$ DNase, 1 U $mL^{-1}$ RNase A and 0.2 mg $mL^{-1}$ lysozyme. $His_6$-tagged TrxB was then purified using a cobalt affinity column before exchanging into RNR storage buffer. Flavin adenine dinucleotide (FAD) content was measured to be 1.13 FAD per TrxB monomer following previously described procedures[1]. Redoxin activity of $His_6$-tagged TrxB was assayed using 5,5'-dithiobis-(2-nitrobenzoic acid) (DTNB) as a substrate at 25 ˚C to be 4.75 μmol DTNB (min mg $TrxB)^{-1}$ as previously described[1].

**S2. Cryo-EM Grid Preparation and Data Collection**

All cryo-EM grid freezing was performed on an FEI Vitrobot Mark IV with the chamber humidity set to 100% and the temperature set to 25°C. QuantiFoil holey carbon R 1.2/1.3 300-mesh copper grids were used for all samples, which were glow discharged on a PELCO easiGlow system for 45 s with 20 mA current prior to sample application. Four μL of sample was applied onto the grid, and the grid was then blotted for 3.5 s and immediately plunge frozen in liquid ethane cooled by liquid nitrogen.

Datasets for all conditions shown in this work were collected on a Talos Arctica (Thermo Fisher Scientific) at the Cornell Center for Materials Research (CCMR). The microscope operates at 200 keV with a Gatan K3 direct electron detector and BioQuantum energy filter (20 eV slit width) at a nominal magnification of 79,000× (nominal pixel size of 1.04 Å $pixel^{-1}$). Movies were collected with a nominal defocus range from –0.8 to –2.0 μm and a total dose of 50 $e^-$ $Å^{-2}$ over 50 frames, with a 3 x 3 multishot scheme in SerialEM[5]. Total numbers of movies collected for each condition are: 4537 for the turnover condition, 3476 for the pre-turnover condition, 3503 for the product condition, 4410 for the pre-reduction condition. The exposure time, frame time, and dose rate for each condition were: 2.23 s, 0.0445 s, and 22.60 $e^-$ $Å^{-2}$ $s^{-1}$ for turnover condition; 2.031 s, 0.0405 s, and 24.77 $e^-$ $Å^{-2}$ $s^{-1}$ for pre-turnover condition; 2.164 s, 0.0435 s, and 22.98 $e^-$ $Å^{-2}$ $s^{-1}$ for product condition; 2.031 s, 0.0405 s, and 24.61 $e^-$ $Å^{-2}$ $s^{-1}$ for pre-reduction condition.

### S3. Cryo-EM Model Building and Refinement

**Consensus refinements of $\alpha_2$ bound with β C-termini and ligands.** For every condition, the sharpened consensus map (Figures S15, S17, S19, S21: purple triangle) showed strong density for residues 311 - 322 of the β C-terminal tail bound to the back of each monomer in $\alpha_2$, with the turnover and product maps showing reasonable density for two additional residues, 309 and 310. Thus, for each condition, consensus models of $\alpha_2$ with bound β C-termini was real-space refined in Phenix[6,7] using the AlphaFold2[8] model of an α monomer (obtained from Uniprot[9] entry P50620) as the starting structure. Two copies of the α monomer model were rigid-body fit into the cryoSPARC consensus map in ChimeraX, and the ligands and bound β C-terminus were then built manually in COOT[10]. *Bs* RNR is known to have three allosteric sites: the S-site, which determines substrate specificity, and the I- and M-sites, which together control the overall activity of the enzyme[3,4]. For all conditions, $Mg^{2+}$-TTP was modeled at the S-site, and an ATP was modeled at the I-site. For the pre-turnover condition, no nucleotide density was observed at the purine-preferring M-site, indicating that ATP does not bind strongly here. Therefore, for the turnover and pre-reduction conditions, a GDP was modeled at the M-site, which was consistent with the density. For the product condition, a dGDP was modeled at the M-site. The restraint file for dGDP was generated using the Grade2 server[11]. The nucleotide density in all refined structures can be visualized in Figure S6. The initial models were refined against the cryoSPARC unsharpened consensus map and then against the sharpened map, after which the atomic model was inspected in COOT to fix issues manually, as well as removing terminal residues with poor density (residues 1 – 5 and 689 – 700 in $\alpha_2$). The density around the active-site cysteine pair (C170/C409) was also inspected (Figure S4); the pre-turnover and product conditions show clearly reduced density, while the pre-reduction shows strong density for a disulfide bond, and the turnover condition shows a mixture between oxidized and reduced density. We thus modeled a disulfide bond between C170 and C409 in COOT for the pre-reduction condition, while the pair was kept reduced in the other conditions. Finally, one or two rounds of real-space refinement against the sharpened map was performed to produce the 4 final consensus atomic models of $\alpha_2$ bound with nucleotides and β C-termini (Table S1).

**Full-complex models of 3D classes.** Because of the lack of atomic detail in the $\beta_2$ and TrxA density, full-complex models were built for every 3D class (37 total, shown in Figure S7) and unsharpened consensus maps (Figures S15, S17, S19, S21: green square) by rigid-body docking atomic models of $\alpha_2$, $\beta_2$, and TrxA into the unsharpened maps obtained from RELION 3D auto refine (Table S2). For consensus maps and 3D classes obtained from individual conditions, the $\alpha_2$ model used for docking was the refined consensus model (bound with β C-termini and ligands) from the corresponding condition (described above). For 3D classes from the all-condition combined refinement, the refined consensus $\alpha_2$ model from the turnover condition was used. For $\beta_2$, residues 16 – 290 and the dimanganese center in both chains from PDB entry 4dr0 were used in docking (with chains A and B renamed to C and D). Residues 19 – 104 of TrxA in the top-ranking model of the AlphaFold2 multimer prediction of the α/TrxA complex were used for docking in maps containing strong TrxA density (turnover consensus full-complex model and all models from the pre-reduction condition). For each full-complex model, clashes at the interfaces between different chains were fixed by either using ISOLDE[12] or by alternating side-chain rotamers.

### S4. All-Atom Molecular Dynamics (MD) Simulations and Umbrella Sampling

Umbrella sampling, an enhanced sampling technique for MD, was performed to investigate the molecular mechanisms for preferential binding of TrxA to one side of $\alpha_2$ in the *Bs* RNR active complex. Two hypotheses were tested. First, we tested whether the flexibility of the interfacial F624α may be coupled to the asymmetry of the $\alpha_2\beta_2$ complex or the oxidation state of the active-site cysteines (C170/C409). The free energy profiles along the rotamer $\chi_1$ angle of F624 obtained by umbrella sampling reveal no significant dependence on the oxidation state of the active-site cysteines or the side of the $\alpha_2$ dimer (Figure S12B), suggesting that F624 does not play a major role in regulating TrxA's binding preference for oxidized α′. Second, we tested whether the oxidation state of the α C-terminal tail affects TrxA binding. For both tests, methods are detailed below. All preparation and MD simulation for in this work were performed in AmberTools23[13] and Amber22[14].

**Starting models for F624 $\chi_1$ umbrella sampling.** To examine the dynamics of residue F624 in α at the TrxA binding interface, MD simulations were performed using starting structures produced by docking a consensus atomic model of $\alpha_2$ bound with β C-termini and the $\beta_2$ crystal structure (PDB: 4dr0) into the map of pre-reduction class 11 (Figure S7D), which displays an open conformation with strong TrxA density. For the starting structure representing the oxidized state, the $\alpha_2$ consensus model from the pre-reduction condition was used (where a disulfide bond is formed between C170 and C409 in both active sites), while for the reduced state, the consensus $\alpha_2$ model from the pre-turnover condition was used (where the C170/409 pair is modeled as reduced in both active sites). Nucleotide ligands at the M- and I-sites were removed as their physiological roles are to prevent filament formation (which is irrelevant in this simulation), while $Mg^{2+}$-TTP was kept at the S-site because of its essential role in dimer formation. Modeller[15] was then used to build unmodeled regions. Both α C-terminal tails were manually moved away from the TrxA binding interface in COOT as transient interactions between the C-termini and the TrxA binding interface were observed in many of our MD runs (results not shown). For the oxidized state, a disulfide bond was also manually introduced between C695 and C698 in both α C-termini using COOT.

**Starting model for 2D umbrella sampling of α-TrxA interactions in oxidized-tail condition.** To examine how the oxidation of the α C-terminus may modulate the interactions between α and TrxA, a starting structure was produced by docking the AlphaFold predicted α/TrxA complex, an additional copy of the α subunit from the predicted α/TrxA complex, and the $\beta_2$ crystal structure (PDB: 4dr0) into the map of pre-reduction class 11 (Figure S7D). Two copies of $Mg^{2+}$-TTP and bound β C-terminal tails were added to the model by aligning the $\alpha_2$ consensus structure from the pre-turnover condition. Modeller was then used to build unmodeled regions. A disulfide bond was manually introduced between C695 and C698 in both α C-termini using COOT. Cysteines in TrxA and $\alpha_2$ active sites were kept reduced.

**Parametrization of TTP.** The TTP pdb file was obtained from the starting structure, and hydrogen atoms were added with the reduce command. The antechamber[16] command then was used to generate the mol2 file for TTP with GAFF2[17], after which the parmchk2 command was used to generate the frcmod file containing all parameters. Following procedures in the literature[18], we modified the partial charges of TTP in the mol2 file in the following fashion. Partial charges for the thymine base and deoxyribose were obtained from the Amber library, while those for the phosphate tail were obtained from a previous work[19]. We then subtracted an equal and small amount of charge from every atom to achieve the correct total charge (-4). We also modified the parameters of the phosphate tail in the frcmod file with those reported in the literature[19].

**Parametrization of the Metal Cofactor in $\beta_2$.** Because of the absence of crystal structures of *B. subtills* $\beta_2$ with the active Mn(III)$_2$–Y• cofactor, we parametrized the cofactor as Mn(II)$_2$ with a protonated tyrosine (Y105β) which is consistent with the experimental condition for PDB entry 4dr0[20]. We assumed that because $\beta_2$ is not closely engaged with $\alpha_2$ in the configurations that we sample in MD, an inactive cofactor form in $\beta_2$ should have minimal impact on the dynamics of interest within $\alpha_2$. Modeling was performed based on the metal center in chain B of 4dr0. Water molecules were not included during parametrization of the metal center. The MCPB.py program[21] was used to perform the parametrization. For input, a total spin of 11 (11/2 for each $Mn^{2+}$ ion) was used, a cutoff distance of 2.8 Å was used, and large_opt was set to 1 to optimize the hydrogen positions of the large model. Quantum chemical calculations were performed with GAMESS-US[22] using the input files generated by MCPB.py. We parametrized both metal sites in $\beta_2$ with the obtained parameters including atomic partial charges and force constants.

**Preparation and parametrization of starting structures.** All starting structures were prepared in the following fashion. Residue names of cysteines involved in disulfide bonds were manually changed to CYX in the pdb file. Protonation states of histidine were then checked with the H++ server[23], and residue names were edited accordingly. LEaP was then used to parametrize the structures and generate the topology and initial coordinate files. The ff19SB force field[24] was used to model the proteins. Disulfide bonds were formed between corresponding sulfur atoms. TTP and the manganese cofactor were modeled with parameters stated above. The $Mg^{2+}$ ion coordinating the triphosphate of TTP was modeled as an unbound ion. The OPC water model[25] along with the matching Li-Merz 12-6 ions model[26,27] were used. The system was solvated in a periodic truncated octahedron box, with at least 10 Å of water from the protein surface to the box edge. The net charge of the protein (-48 for $\alpha_2/\beta_2$, -56 for $\alpha_2/\beta_2$/TrxA) was neutralized by addition of $Na^+$ counterions. Additional $Na^+$ and $Cl^-$ ions were added to reach a concentration of 150 mM NaCl, and additional $Mg^{2+}$ and $Cl^-$ ions were added to reach a concentration of 15 mM $MgCl_2$. The systems contained ~280K atoms for $\alpha_2/\beta_2$ and ~380K atoms for $\alpha_2/\beta_2$/TrxA. Initial system setup information for different conditions can be found in Table S4.

**Relaxation and equilibration of systems.** For all simulations, a Langevin thermostat with a collision frequency of 1 $ps^{-1}$ was used for temperature control, while the Monte Carlo barostat[28] was used for pressure control with a target pressure of 1 bar. A cut-off distance of 10 Å was used for nonbonded interactions. The SHAKE algorithm[29] was used to constrain bond lengths involving hydrogen atoms. Minimizations were performed with pmemd.MPI, while MD simulations were performed with pmemd.cuda[30,31]. All systems generated from starting models were relaxed and equilibrated in the same fashion. First, the systems are minimized with 5000 cycles of steepest descent with all atoms restrained with a 100 kcal $mol^{-1}$ $Å^{-2}$ force constant. The systems are then gradually heated up from 100 K to 310 K over 1 ns with a 1 fs time step under the NVT ensemble while restraining protein and ligand atoms with a 100 kcal $mol^{-1}$ $Å^{-2}$ force constant. This was followed by two rounds of relaxation at 310 K over 1 ns with a 1 fs time step under the NPT ensemble, with a restraining force constant of 100 kcal $mol^{-1}$ $Å^{-2}$ for the first round and 10 kcal $mol^{-1}$ $Å^{-2}$ for the second round on protein and ligand atoms. The protein side chains were then minimized along with solvent with 2000 cycles of steepest descent followed by 3000 cycles of conjugate gradient, while a 10 kcal $mol^{-1}$ $Å^{-2}$ force constant was used to restrain the backbone atoms and ligand TTP. The protein atoms were then relaxed through three rounds of simulation at 310 K over 1 ns with a 1 fs time step under the NPT ensemble, with decreasing restraining force constraints of 10 kcal $mol^{-1}$ $Å^{-2}$, 1 kcal $mol^{-1}$ $Å^{-2}$, and 0.1 kcal $mol^{-1}$ $Å^{-2}$, applied to the backbone atoms and ligand TTP. Finally, the systems were relaxed with a simulation at 310 K over 1 ns with a 1 fs time step under the NPT ensemble with no restraint to produce the relaxed configuration. The relaxed configuration was further equilibrated at 310 K for 20 ns with a 2 fs time step under the NPT ensemble to produce the initial configurations used for production runs. Root-mean-square deviations (RMSDs) of the 20 ns unconstrained equilibration runs are plotted in Figure S32, showing that individual protein components were well-equilibrated after 10 ns in all runs, although the full complex exhibited larger deviations due to the flexible tails and drift in orientation between $\alpha_2$ and $\beta_2$.

**Umbrella sampling for F624 $\chi_1$ angle.** We performed umbrella sampling[32] on the $\chi_1$ angle of F624 on both sides of $\alpha_2$, and with both oxidized and reduced active-site cysteines in $\alpha_2$. The four sets of umbrella sampling were performed in an identical fashion. Simulations were run at 310 K under the NVT ensemble with a time step of 2 fs. During all simulations, a weak restraining force constant of 0.01 kcal mol$^{-1}$ Å$^{-2}$ was applied to the Cα atom of residues in rigid parts of $\alpha_2$ (residues 8-685) and $\beta_2$ (residues 18-271) as well as the α C-terminal tails (residue 686-700), in order to prevent the complex from drifting away from the initial $\beta_2$ configuration and to prevent the α C-terminal tails from transiently interacting with the TrxA binding interface (α C-terminal tails are far away from the interface in the initial configurations used). A total of 72 simulation windows were performed, spanning 360° with an increment of 5°. The $\chi_1$ angle observed in the initial configuration (-60° or -180°) was used as the angle restraint for the first window. The starting structure for equilibration in each window was updated every 6-window batch (30°). The initial configuration was used as the starting structure for the first 6 windows, while an equilibrated configuration from the last window of the preceding 6-window batch was used as the starting structure for the subsequent 6-window batch. An angle restraint force constant of 200 kcal mol$^{-1}$ rad$^{-2}$ was used. For each window, the system was first equilibrated under restraint for 5 ns, after which the $\chi_1$ angle of F624 was recorded every 10 fs for 15 ns. A total sampling of 1.44 μs was performed for each set of umbrella sampling. Histograms of sampled angle in each window are plotted in Figure S33, which shows sufficient overlap between windows. The free energy profiles were reconstructed with the weighted histogram analysis method (WHAM)[33]. Reconstruction was performed from -65° to 300° with 360 bins and a convergence tolerance of 0.0001.

**Initial configuration for 2D umbrella sampling of α tail in reduced-tail condition.** In order to keep initial configurations as close as possible between oxidized-tail and reduced-tail conditions, we generated the initial configuration for the reduced-tail condition from the equilibrated oxidized-tail configuration. The equilibrated configuration of the oxidized-tail condition was converted to a pdb file using CPPTRAJ[34] including water and ions, in which oxidized CYX residues (C695 and C698) were renamed to CYS residues. LEaP was used to convert this pdb file back into the topology and coordinate files, and the PARMED program[35] was used to set the box parameters in the files to those of the equilibrated oxidized-tail configuration. This configuration was first minimized with 2000 cycles of steepest descent followed by 3000 cycles of conjugate gradient, while a 10 kcal $mol^{-1}$ $Å^{-2}$ force constant was applied on all protein and ligand atoms except for the side chain atoms of C695 and C698 on $\alpha_2$. The minimized configuration was then relaxed with 1 ns simulation with 1 fs time step at 310 K under the NPT ensemble, while a weak restraining force constant of 0.01 kcal $mol^{-1}$ $Å^{-2}$ was applied to the Cα atom of residues in rigid parts of $\alpha_2$ (residues 8-685), $\beta_2$ (residues 18-271), and TrxA (residues 19-103). The resulting configuration was used as the initial configuration for 2D umbrella sampling of the reduced-tail condition.

**2D umbrella sampling for α tail.** We performed 2D umbrella sampling on the α C-terminal tail in the TrxA bound conformation with both oxidized and reduced α C-terminal tails in an identical fashion. Simulations were run at 310 K under the NVT ensemble with a time step of 2 fs. During simulation, a weak restraining force constant of 0.01 kcal $mol^{-1}$ $Å^{-2}$ was applied to the Cα atom of residues in rigid parts of $\alpha_2$ (residues 8-685), $\beta_2$ (residues 18-271), and TrxA (residues 19-103), in order to prevent the complex from drifting away from the initial $\beta_2$ configuration. 2D umbrella sampling was performed on two distances defined between the α C-terminal tail and its binding interface with TrxA (Figure S13). The COM of the binding interface was defined as the COM of Cα atoms of W623 on α and S71, I72 and G89 on TrxA. Distance 1 was then defined as the distance between the Cα atom of S697 on α and the COM of the binding interface. Distance 2 was defined as the distance between the COM of Cα atoms of C698 and V699 on α, and the COM of the binding interface. We sampled distance 1 from 4.0 Å to 8.5 Å in 10 windows (0.5 Å increment) and distance 2 from 5.5 Å to 10 Å in 10 windows (0.5 Å increment), resulting in a total of 100 windows. The first window (with window centers of 4.0 Å and 5.5 Å) used the initial configuration as its starting model for equilibration, while subsequent windows used the equilibrated configuration from a window adjacent to themselves as the starting structure. A force constant of 7 kcal $mol^{-1}$ $Å^{-2}$ was used for both distance restraints. For each window, the system was first equilibrated under restraint for 5 ns, after which the two distances were recorded every 20 fs for 25 ns. A total sampling of 3 μs was performed for each set of 2D umbrella sampling. 2D histograms of sampled distances were visually inspected, and regions of low sampling were subjected to two additional sampling windows centered at (distance 1, distance 2) = (5.75 Å, 7.25 Å) and (7.75 Å, 5.75 Å) for the oxidized tail and (6.25 Å, 6.75 Å) and (6.25 Å, 7.0 Å) for the reduced tail with a higher force constant restraint (10 kcal $mol^{-1}$ $Å^{-2}$). The final 2D histograms, plotted in Figure S34, display sufficient distance sampling. The 2D free energy profiles were then reconstructed from 102 windows with WHAM, from 4.0 Å to 8.5 Å for distance 1 and 5.0 Å to 10.0 Å for distance 2, with 100 bins in both dimensions, and a convergence tolerance of 0.0001.

### S5. Phylogenetic and Bioinformatic Analyses of TrxA-α Interactions in Class I RNRs

The class I clade from the all-RNR phylogenetic tree published previously[36] was pruned and used for the analysis. Full sequences corresponding to those in the tree were retrieved from Uniprot[9] and aligned with MAFFT[37]. The sequence alignment around the TrxA binding interface was inspected and manually adjusted to improve alignment of those residues, which was then used for further analyses to identify sequences with hydrophobic interfacial residues. Figure 6A was plotted using ggtree[38]. Uniprot accession codes of sequences used for AlphaFold2 multimer predictions of TrxA-α complexes are: *Aquifex aeolicus*, O66503 α, O67747 TrxA1; *Bacillus subtilis*, P50620 α, P14949 TrxA; *Chitinivibrio alkaliphilus*, U7D2Y2 α, U7D5E5 TrxA; *Escherichia coli*, P00452 α, P0AA25 TrxA; *Homo sapiens*, P23921 α, P10599 Txn; *Salmonella typhimurium*, Q08698 α, P0AA28 TrxA.

### S6. AlphaFold Predictions

All AlphaFold predictions were performed in AlphaFold2 version 2.3[8,39] using the full sequence database to generate the multiple sequence alignments and a max template date of 2023-03-22.

**Table S1.** EM data processing and consensus refinement statistics.

| **Condition** | Turnover | Pre-turnover | Product | Pre-reduction |
|---|---|---|---|---|
| EMDB | EMD-45031 | EMD-44947 | EMD-45017 | EMD-45004 |
| PDB | 9byh | 9bw3 | 9by1 | 9bxc |
| **Data collection and processing** | | | | |
| Microscope | Talos Arctica | Talos Arctica | Talos Arctica | Talos Arctica |
| Camera | K3 | K3 | K3 | K3 |
| Nominal Magnification | 79,000 | 79,000 | 79,000 | 79,000 |
| Voltage (keV) | 200 | 200 | 200 | 200 |
| Electron exposure ($e^-/Å^2$) | 50 | 50 | 50 | 50 |
| Defocus range (μm) | -0.8 to -2.0 | -0.8 to -2.0 | -0.8 to -2.0 | -0.8 to -2.0 |
| Pixel size (Å) | 1.017 | 1.022 | 1.017 | 1.014 |
| Micrographs used (no.) | 3258 | 2532 | 2777 | 3785 |
| Initial particles (no.) | 1,010,952 | 892,650 | 990,504 | 551,703 |
| Final particles (no.) | 595,923 | 665,600 | 504,919 | 360,764 |
| Symmetry imposed | C1 | C1 | C1 | C1 |
| Map resolution (Å)<br>FSC threshold | 2.53<br>(0.143) | 2.42<br>(0.143) | 2.55<br>(0.143) | 2.76<br>(0.143) |
| Map resolution range (Å) | 2.244 - 25.796 | 2.289 – 36.529 | 2.264 – 32.719 | 2.397 – 38.611 |
| **Refinement** | | | | |
| Model resolution (Å)<br>FSC threshold | 2.7<br>(0.5) | 2.6<br>(0.5) | 2.8<br>(0.5) | 3.0<br>(0.5) |
| Map sharpening B factor ($Å^2$) | 103.2 | 93.5 | 101.9 | 109.1 |
| Model composition<br>Non-hydrogen atoms<br>Protein residues<br>Ligands | <br>11526<br>1394<br>TTP, MG, ATP, GDP | <br>11442<br>1390<br>TTP, MG, ATP | <br>11524<br>1394<br>TTP, MG, ATP, DGI | <br>11498<br>1390<br>TTP, MG, ATP, GDP |
| B factors ($Å^2$)<br>Protein<br>Ligand | <br>34.19<br>43.72 | <br>33.32<br>42.06 | <br>36.21<br>46.36 | <br>46.42<br>56.74 |
| R.m.s. deviations<br>Bond lengths (Å)<br>Bond angles (°) | <br>0.002 (0)<br>0.497 (0) | <br>0.002 (0)<br>0.423 (0) | <br>0.003 (0)<br>0.486 (0) | <br>0.003 (0)<br>0.536 (0) |
| Validation<br>MolProbity score<br>Clashscore<br>Rotamer outliers (%) | <br>1.52<br>9.86<br>0.89 | <br>1.85<br>22.73<br>0.81 | <br>1.59<br>8.90<br>0.49 | <br>1.70<br>11.97<br>0.81 |
| Ramachandran plot<br>Disallowed (%)<br>Allowed (%)<br>Favored (%) | <br>0.00<br>1.88<br>98.12 | <br>0.00<br>1.74<br>98.26 | <br>0.00<br>2.60<br>97.40 | <br>0.00<br>2.53<br>97.47 |

**Table S2.** Details for additional maps and docking models

| | | EMDB | PDB | Particle Number | Resolution (Å) | Pixel size (Å) |
|---|---|---|---|---|---|---|
| Turnover | Unsharpened Consensus | EMD-45037 | 9byl | 596609 | 2.99 | 1.017 |
| | Class 1 | EMD-45044 | 9byt | 65320 | 3.52 | 1.017 |
| | Class 4 | EMD-45045 | 9byv | 34075 | 3.83 | 1.017 |
| | Class 5 | EMD-45046 | 9byw | 13554 | 4.64 | 1.017 |
| | Class 8 | EMD-45047 | 9byx | 29382 | 3.94 | 1.017 |
| | Class 9 | EMD-45048 | 9byy | 22787 | 4.07 | 1.017 |
| | Class 12 | EMD-45049 | 9byz | 28388 | 3.94 | 1.017 |
| | Class 14 | EMD-45051 | 9bz2 | 30885 | 3.83 | 1.017 |
| | Class 17 | EMD-45052 | 9bz3 | 26267 | 4.01 | 1.017 |
| Pre-turnover | Unsharpened Consensus | EMD-44985 | 9bwx | 667306 | 2.91 | 1.022 |
| | Class 2 | EMD-44991 | 9bx2 | 30614 | 3.79 | 1.022 |
| | Class 5 | EMD-44992 | 9bx3 | 26273 | 3.90 | 1.022 |
| | Class 9 | EMD-44995 | 9bx6 | 68044 | 3.44 | 1.022 |
| | Class 12 | EMD-44999 | 9bx8 | 48084 | 3.59 | 1.022 |
| | Class 15 | EMD-45000 | 9bx9 | 32509 | 3.79 | 1.022 |
| Product | Unsharpened Consensus | EMD-45018 | 9by2 | 455171 | 3.10 | 1.017 |
| | Class 3 | EMD-45019 | 9by3 | 56752 | 3.57 | 1.017 |
| | Class 8 | EMD-45020 | 9by7 | 45014 | 3.67 | 1.017 |
| | Class 9 | EMD-45021 | 9by8 | 29205 | 3.88 | 1.017 |
| | Class 10 | EMD-45023 | 9by9 | 19330 | 4.14 | 1.017 |
| | Class 11 | EMD-45024 | 9bya | 24384 | 4.01 | 1.017 |
| | Class 12 | EMD-45026 | 9byc | 26170 | 3.94 | 1.017 |
| | Class 16 | EMD-45029 | 9byd | 19182 | 4.20 | 1.017 |
| | Class 19 | EMD-45030 | 9byg | 39606 | 3.77 | 1.017 |
| Pre-reduction | Unsharpened Consensus | EMD-45010 | 9bxs | 367506 | 3.37 | 1.014 |
| | TrxA Focus-classified | EMD-45011 | 9bxt | 170747 | 2.88 | 1.014 |
| | Class 11 | EMD-45014 | 9bxx | 26401 | 4.26 | 1.014 |
| | Class 15 | EMD-45015 | 9bxz | 8136 | 8.11 | 1.014 |
| | Class 16 | EMD-45016 | 9by0 | 34996 | 4.19 | 1.014 |
| Combined | Class 6 | EMD-45053 | 9bz5 | 81905 | 3.93 | 1.014 |
| | Class 7 | EMD-45054 | 9bz6 | 112947 | 3.87 | 1.014 |
| | Class 15 | EMD-45057 | 9bz9 | 22630 | 4.64 | 1.014 |
| | Class 18 | EMD-45061 | 9bza | 73289 | 3.93 | 1.014 |
| | Class 23 | EMD-45064 | 9bzd | 88786 | 3.82 | 1.014 |
| | Class 26 | EMD-45065 | 9bze | 55609 | 4.19 | 1.014 |
| | Class 28 | EMD-45066 | 9bzf | 27691 | 4.40 | 1.014 |
| | Class 29 | EMD-45067 | 9bzh | 16571 | 5.90 | 1.014 |
| | Class 31 | EMD-45068 | 9bzi | 66248 | 3.99 | 1.014 |
| | Class 40 | EMD-45069 | 9bzj | 53114 | 4.12 | 1.014 |
| | Class 43 | EMD-45070 | 9bzk | 46987 | 4.19 | 1.014 |
| | Class 45 | EMD-45071 | 9bzm | 49977 | 4.19 | 1.014 |
| | Class 50 | EMD-45072 | 9bzo | 26473 | 4.48 | 1.014 |

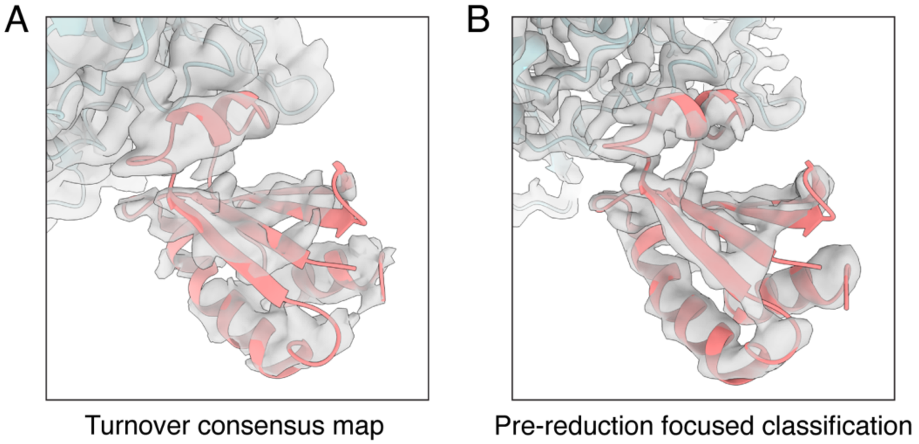


**Figure S1.** Close-up of TrxA model fit in EM maps. (A) AlphaFold2 prediction of TrxA (coral) fits unambiguously in the extra density found in our unsharpened consensus map of the turnover condition (Figure 2A, Table S2), providing the first visualization of a thioredoxin-RNR interaction. (B) Focused classification of TrxA with the pre-reduction dataset (Figure S21, S22B, Table S2) resulted in a map with improved TrxA density, although residue-level information remains unresolved.

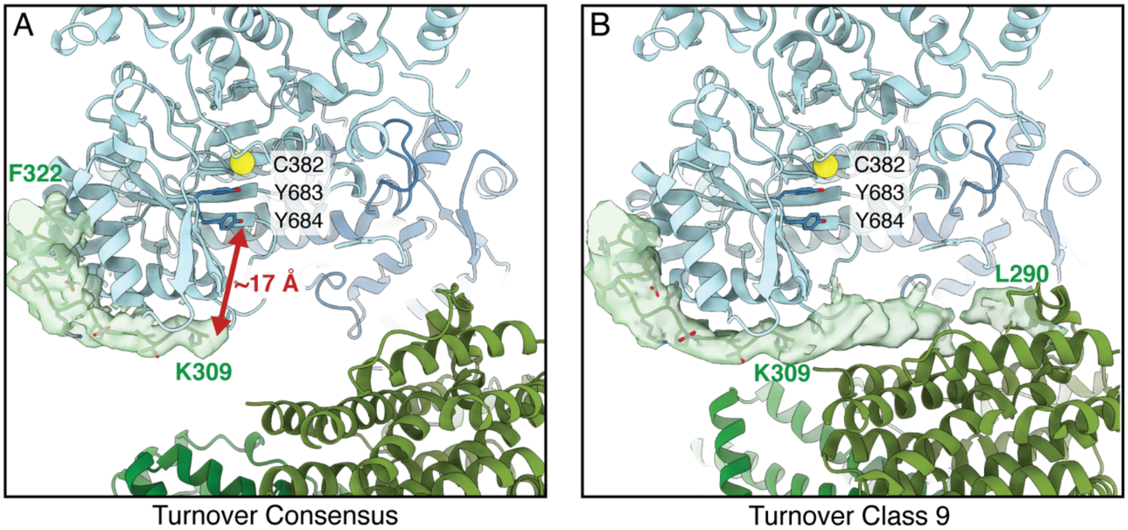


**Figure S2.** Density for the β C-terminal tail in the turnover condition. (A) The consensus reconstruction for the turnover condition shows clear density for K309-F322 of the β C-terminus on both sides of the $\alpha_2\beta_2$ complex (tail density in chain D shown as green surface), while residues 291-308 are unresolved. The interfacial PCET residue Y307β is not resolved, but given that it is two residues from K309β, we believe that it is within the ~10-Å range needed for orthogonal PCET with Y684α in the active site. The PCET pathway in α proceeds through the Y684/Y683 dyad to generate a thiyl radical on C382. (B) 3D classification of the turnover condition yields several classes (e.g., classes 1, 9, 12 in Figure S7A, Table S2) with density for the β′ C-terminus (green surface) on the α′/β′ side that stretches across the gap between K309β′ and L290β′. The tail residues 291-308 were not modeled into the density as the density does not contain atomic details, likely due to the flexibility of this tail region, which is suspended across a gap.

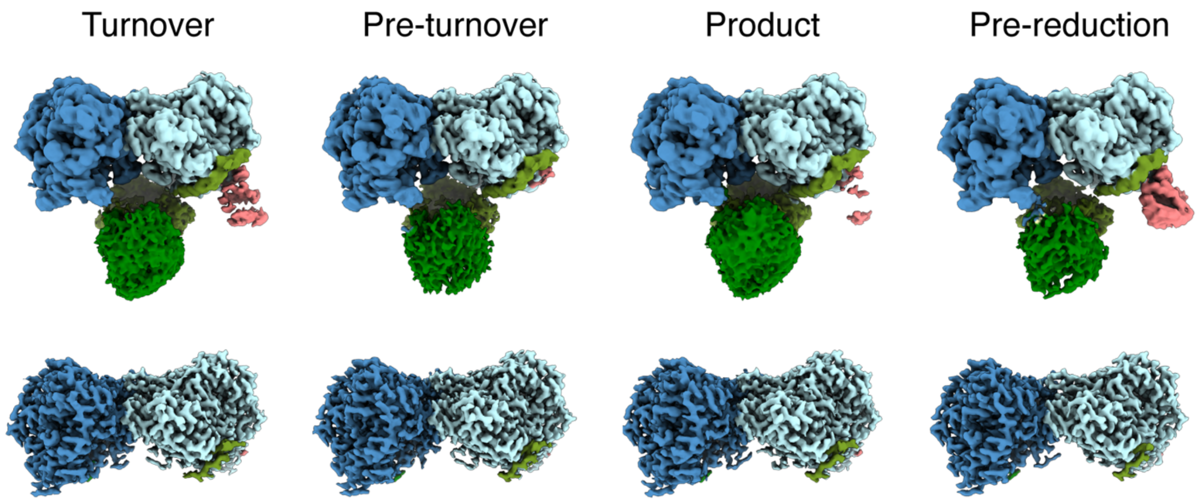


**Figure S3.** Consensus reconstructions for each condition. Volumes are colored by protein subunit in the same scheme as in Figure 2. Unsharpened maps (Table S2) are shown in the top row while sharpened maps (Table S1) are shown in the bottom row. All reconstructions show similarly blurry density for the core part of β2 (green) but with varying TrxA occupancy (coral). The β C-terminus (green) is resolved on both sides of $\alpha_2$ in all sharpened maps.

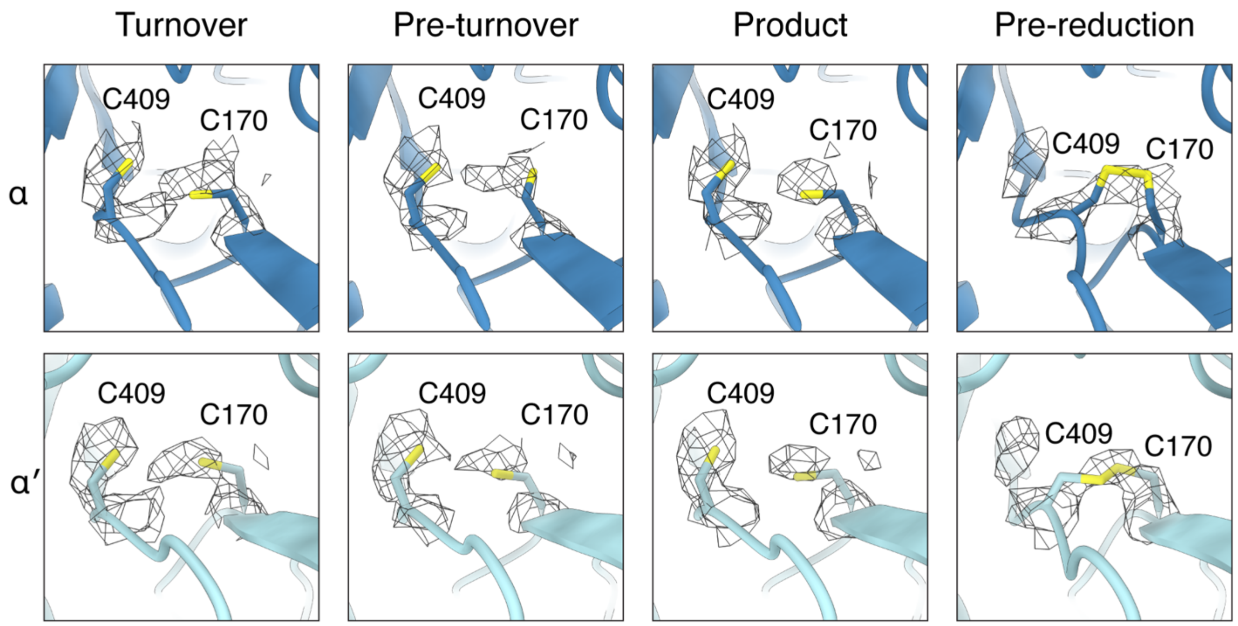


**Figure S4.** Density for active-site cysteines C170/C409 in the sharpened consensus maps, shown by condition and $\alpha_2$ protomer. Disulfide bond formation results in the movement of C409 towards C170. Only the pre-reduction map shows density that is largely consistent with disulfide formation. The turnover map shows mixed density for oxidized and reduced C170/C409 but was modeled as reduced because of clear density for C409 in the reduced conformation. The pre-turnover and product maps show density that is largely consistent with the reduced conformation of this cysteine pair. The pre-turnover map is shown here at a threshold of 1.2 while other maps are shown at a threshold of 0.9.

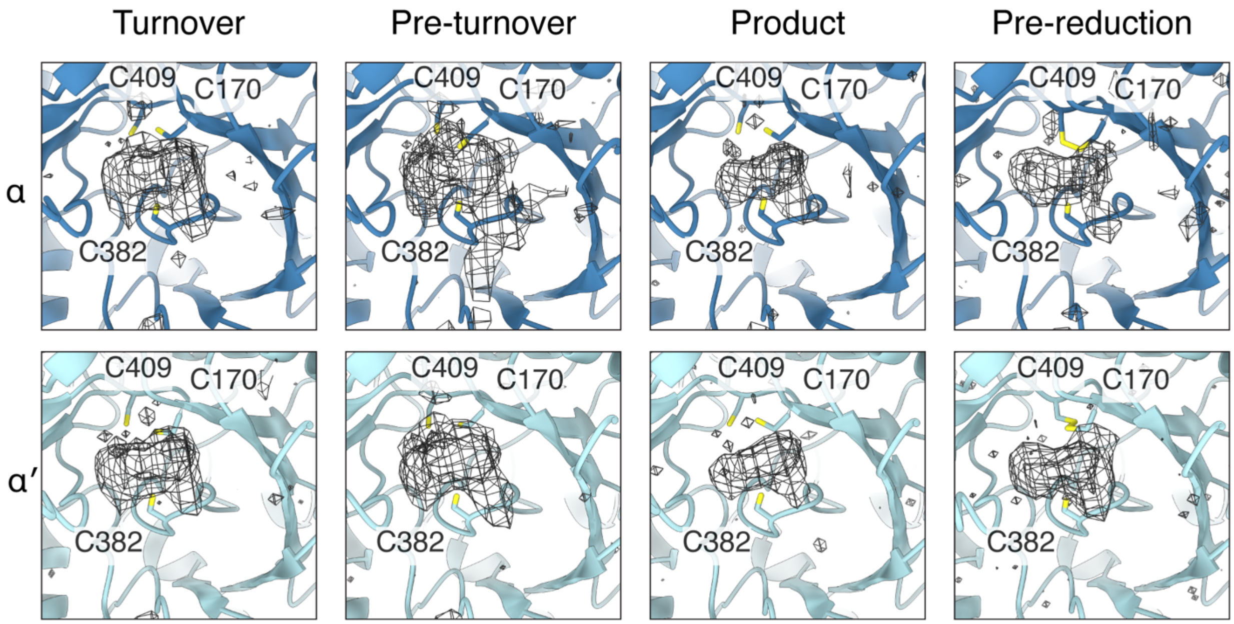


**Figure S5.** Unmodeled density at the active site in the unsharpened consensus maps, shown by condition and $\alpha_2$ protomer. Nucleotide (substrate or product) and the α C-terminus can bind the same pocket in *B. subtills* RNR[4], regardless of the C170/C409 oxidation state without clashes. The excess density in the pre-turnover map likely represents the α C-terminus dynamically sampling this pocket, while in other maps, it likely represents an average of nucleotide and α C-terminus sampling the pocket.

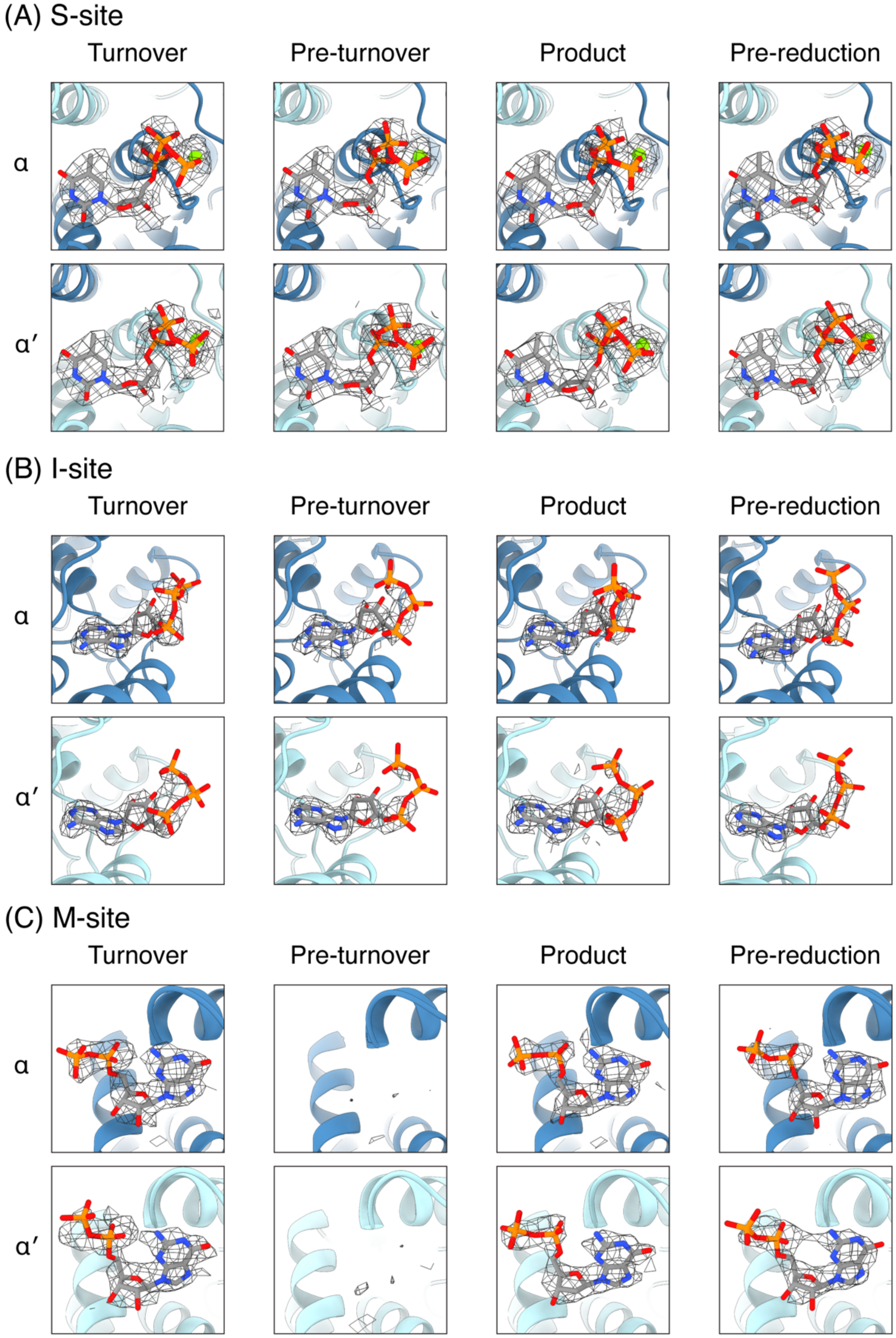


**Figure S6.** Nucleotide density in refined consensus models of $\alpha_2$. (A) Density at the S-site is consistent with the specificity effector, TTP, coordinating a $Mg^{2+}$ ion as commonly seen in RNR structures. (B) Density at the I-site is consistent with ATP with a disordered γ-phosphate as observed previously in a crystal structure of *B. subtilis* $\alpha_2$[4]. (C) Density is observed in the M-site for conditions that contain GDP (substrate) or dGDP (product). Guanine nucleotides act as activators when bound to the M-site in *B. subtilis* RNR[40].

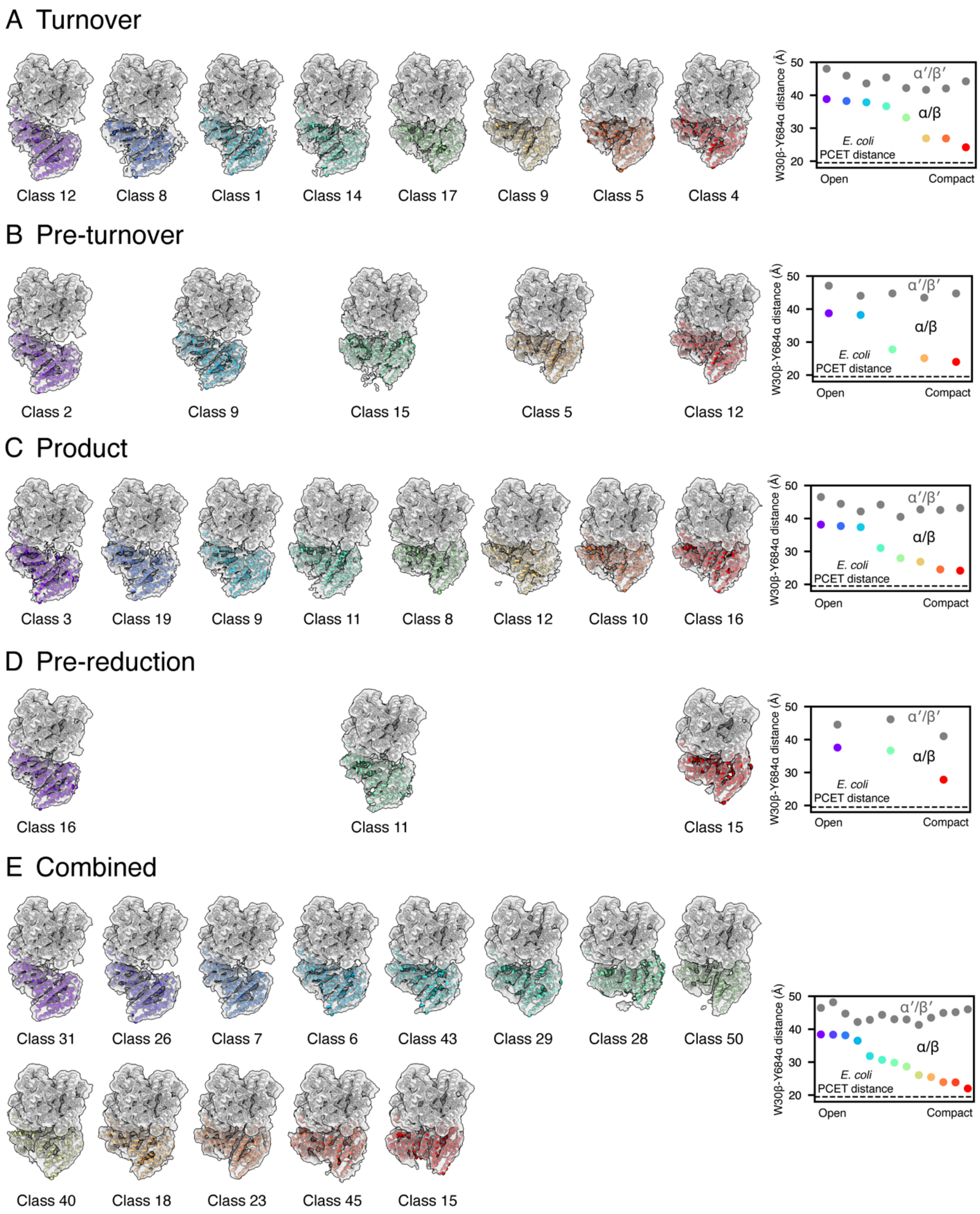


**Figure S7.** 3D classes from each condition and the combined refinement (Table S2), in decreasing order of W30β-Y684α distance between the α/β pair. W30β-Y684α distances between α/β and α′/β′ for each class are plotted on the right for each dataset (with colors for α/β distances matching those of the corresponding 3D classes). (A) Turnover condition. (B) Pre-turnover condition. (C) Product condition. (D) Pre-reduction condition. (E) Combined refinement from all conditions.

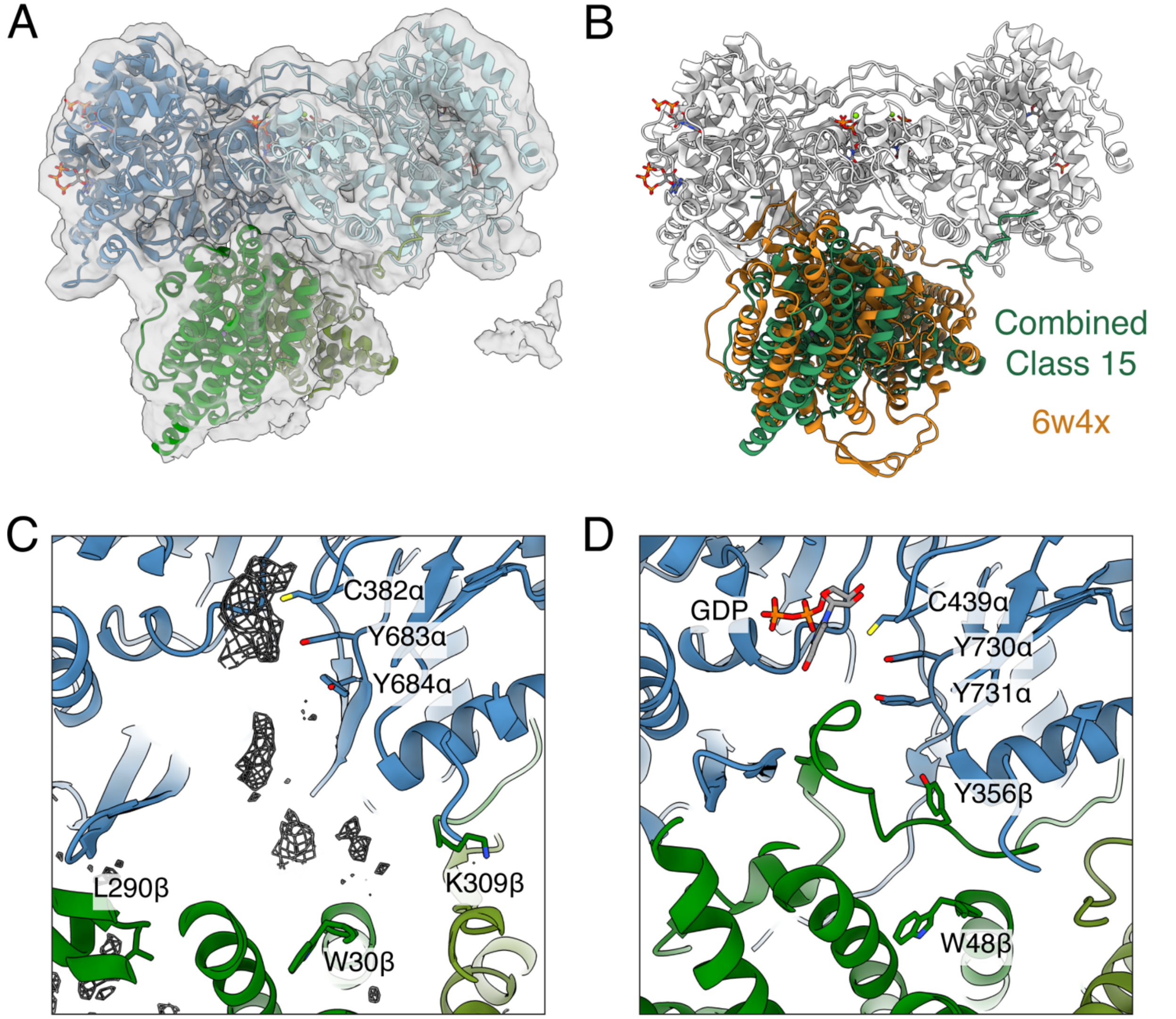


**Figure S8.** Comparison of PCET-like conformation observed for *B. subtilis* RNR with the cryo-EM structure of the *E. coli* RNR active complex[41]. PCET involves alignment of a series of residues Y105β→ W30β→Y307β→Y684α→Y683α→C382α in *Bs* numbering (in *Ec* numbering: Y122β→W48β→Y356β →Y731α→Y730α→C439α) (A) Atomic model of the most compact class from the combined refinement (class 15 in Figure S7E, Table S2). (B) Models of class 15 from combined refinement (green) and *E. coli* RNR active complex (pdb: 6w4x) (orange) aligned by the $\alpha_2$ subunit (white) show resemblance. $\alpha_2$ subunit in 6w4x is not shown. (C) However, the active site of class 15 does not display density for a folded β C-terminal tail as was observed in the *E. coli* RNR active complex, indicating that the conformation captured in class 15 does not exactly correspond to the PCET process. Instead, the residues between L290β and K309β of the β C-terminus remain disordered in class 15 (unmodeled density shown as mesh). (D) In the active site of *E. coli* RNR active complex, the folded β C-terminal tail positions Y356β for the PCET process.

**Table S3.** DensMAP 2D embeddings of 3D classes.

| 3D class | Dens-MAP1 | Dens-MAP2 | 3D class | Dens-MAP1 | Dens-MAP2 |
|---|---|---|---|---|---|
| Combined Class 15 | 13.44 | -0.19 | Product Class 10 | 11.33 | 6.32 |
| Combined Class 18 | 8.63 | 8.44 | Product Class 11 | 5.90 | 8.34 |
| Combined Class 23 | 9.53 | 7.44 | Product Class 12 | 8.70 | 8.47 |
| Combined Class 26 | 0.23 | 5.29 | Product Class 16 | 10.51 | 7.50 |
| Combined Class 28 | 10.25 | 0.46 | Product Class 19 | 0.37 | 7.17 |
| Combined Class 29 | 8.39 | 6.49 | Product Class 3 | 0.13 | 6.08 |
| Combined Class 31 | 0.12 | 6.17 | Product Class 8 | 11.74 | 5.18 |
| Combined Class 40 | 10.91 | 6.87 | Product Class 9 | 1.24 | 6.43 |
| Combined Class 43 | 4.42 | 7.44 | Pre-reduction Class 11 | 0.19 | 6.25 |
| Combined Class 45 | 12.27 | 2.09 | Pre-reduction Class 15 | 11.79 | 5.71 |
| Combined Class 50 | 9.86 | 5.96 | Pre-reduction Class 16 | 0.48 | 7.07 |
| Combined Class 6 | 1.98 | 6.34 | Turnover Class 12 | -0.28 | 5.66 |
| Combined Class 7 | 0.20 | 7.22 | Turnover Class 14 | 0.11 | 6.70 |
| Pre-turnover Class 12 | 8.64 | 7.67 | Turnover Class 17 | 7.90 | 5.75 |
| Pre-turnover Class 15 | 12.40 | 2.49 | Turnover Class 1 | 0.55 | 7.07 |
| Pre-turnover Class 2 | -0.22 | 5.87 | Turnover Class 4 | 8.56 | 7.92 |
| Pre-turnover Class 5 | 10.67 | 6.94 | Turnover Class 5 | 10.99 | 8.34 |
| Pre-turnover Class 9 | 0.42 | 6.96 | Turnover Class 8 | 0.28 | 6.29 |
| | | | Turnover Class 9 | 10.02 | 7.33 |

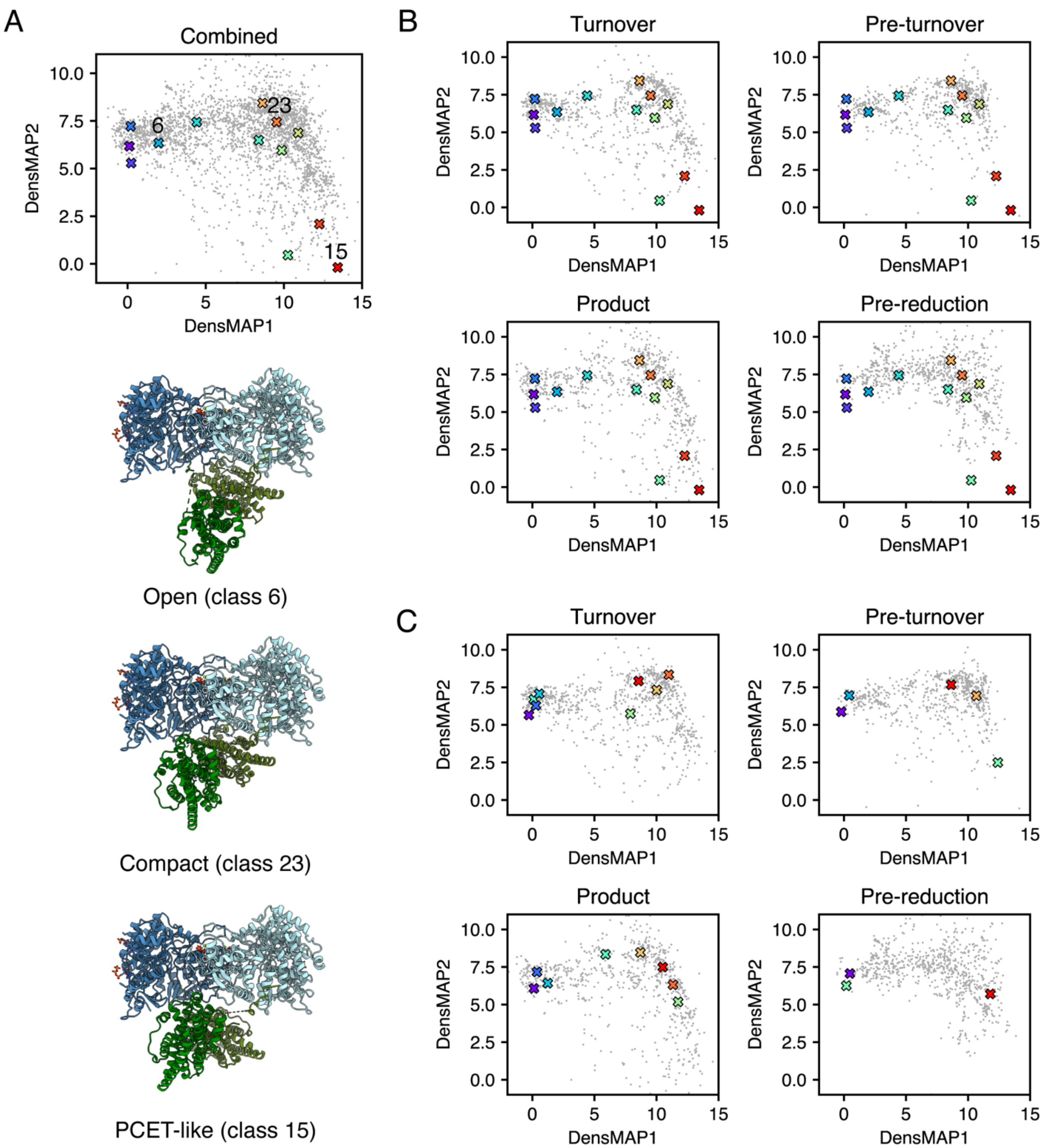


**Figure S9.** DensMAP 2D embeddings, where each gray dot represents a conformation observed in a cluster volume from cryoDRGN landscape analysis and 3D classes are marked as crosses and colored according to class colors in Figure S7. Coordinates of the 3D classes are also provided in Table S3. (A) 3D classes from combined refinement (Figure S7E, Table S2) plotted with combined embedding. Models for the three 3D classes marked in Figure 4 (combined classes 6, 15, and 23) are shown in the bottom left. (B) 3D classes from combined refinement (Figure S7E, Table S2) plotted with embeddings of individual conditions. (C) 3D classes from individual conditions (Figure S7A-D, Table S2) plotted with embeddings of individual conditions.

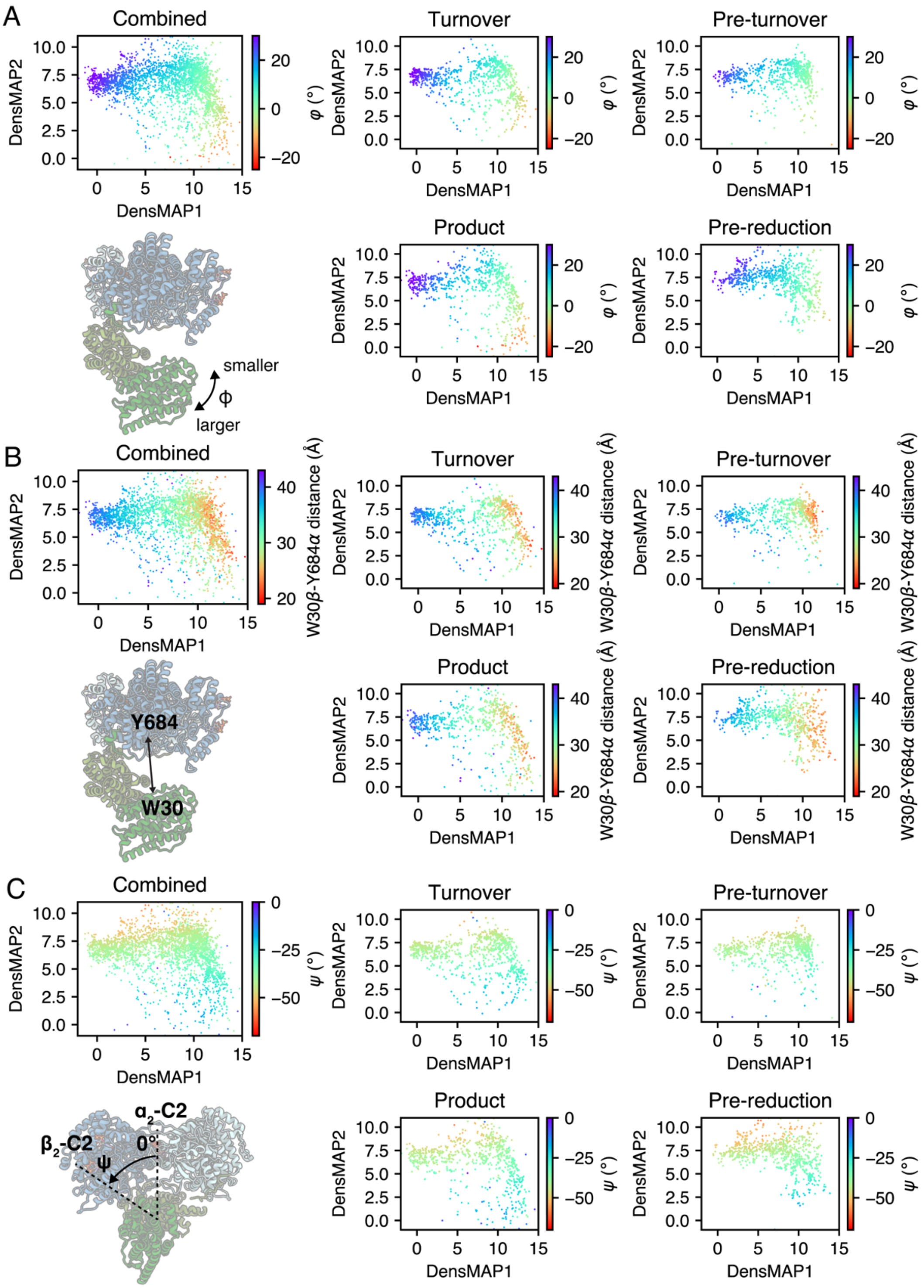


**Figure S10**. DensMAP 2D embeddings, where each dot represents a conformation of a cluster volume from cryoDRGN landscape analysis colored by (A) Euler angle $\varphi$ which describes the openness of $\beta_2$ relative to $\alpha_2$, and (B) W30β-Y684α distances between α/β, which represent the accessibility of the PCET process. Both coloring schemes indicate that the trajectory from the top left corner to the bottom right corner in the embeddings roughly corresponds the continuous transition from open to compact $\alpha_2\beta_2$ conformations. (C) 2D embeddings colored by Euler angle $\psi$ (the angle between the C2-axis of $\alpha_2$ and the C2-axis of $\beta_2$) show that perfectly symmetric conformations (where $\psi = 0°$) are not observed in our landscapes.

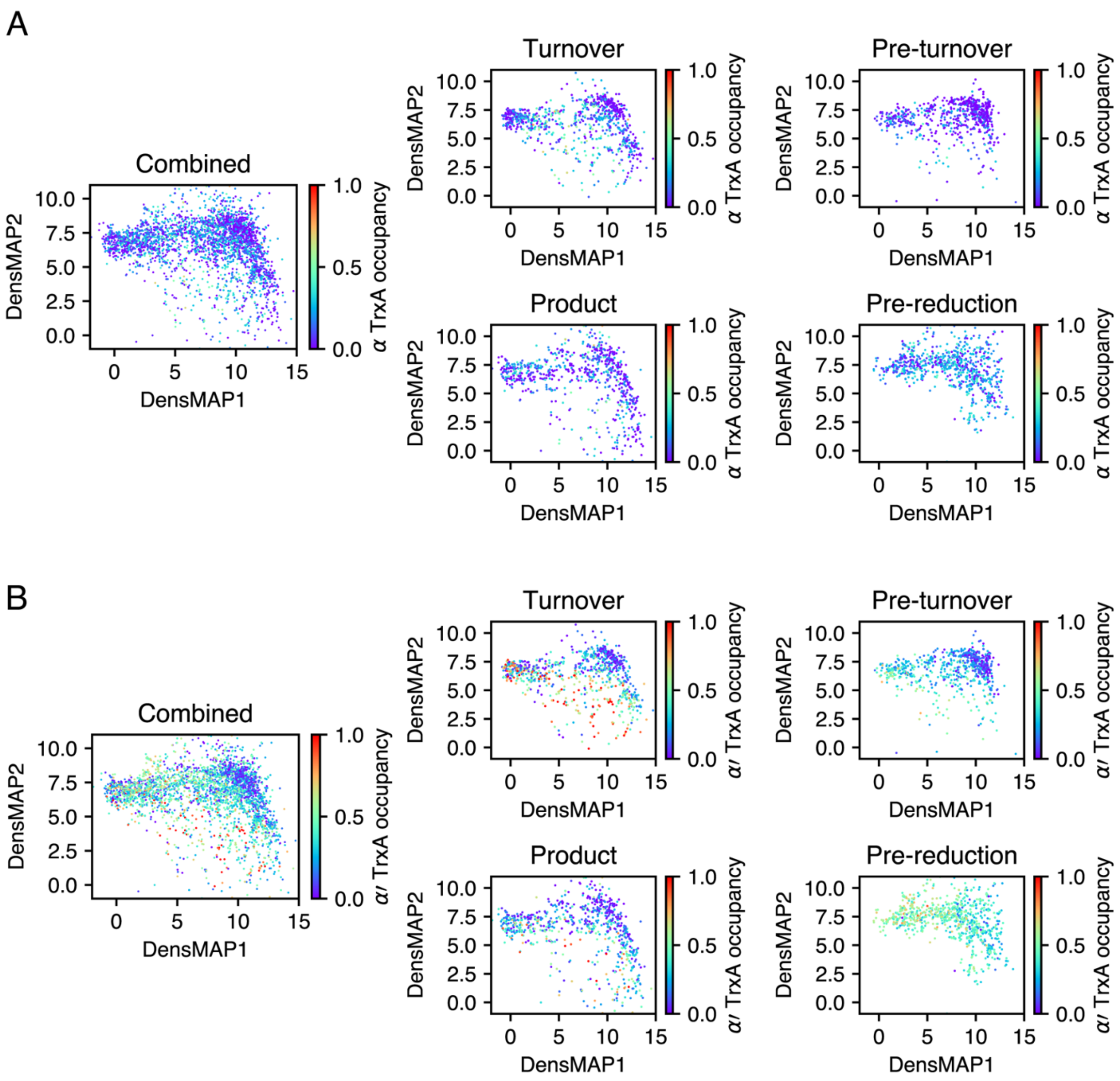


**Figure S11.** TrxA occupancy in the cluster volumes from cryoDRGN analysis. (A-B) DensMAP 2D embeddings, where each dot represents a conformation of a cluster volume from cryoDRGN landscape analysis colored by TrxA occupancy on (A) α or (B) α′. Overall, we see that TrxA prefers to bind oxidized $\alpha_2$ on the α′ side in open conformations.

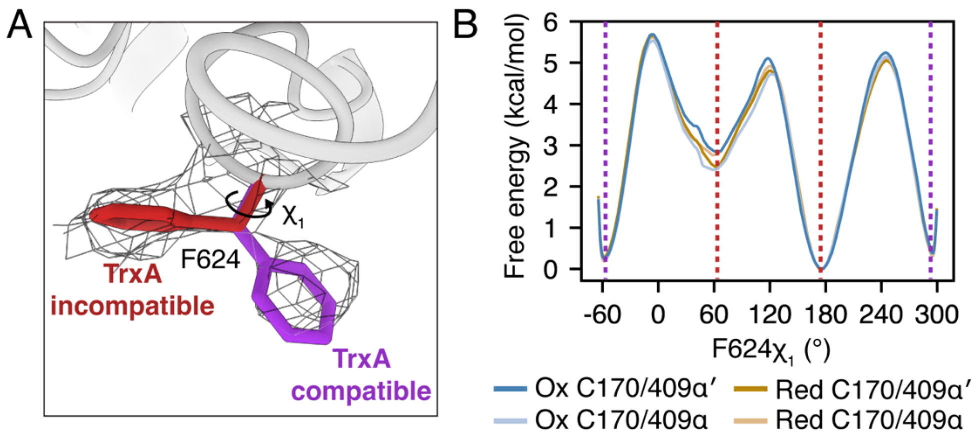


**Figure S12.** Investigation of the role of F624α in TrxA binding with MD simulations. (A) F624α displays two rotamer conformations (density shown as mesh) when TrxA is not bound: one (purple) is compatible with TrxA, while the other (red) conflicts. (B) Free energy profile along the rotamer $\chi_1$ angle of F624 is not dependent on the oxidation state of the active site or whether it is on the α or α′ side of the RNR complex. The dashed lines correspond to TrxA compatible (purple) and incompatible (red) conformations.

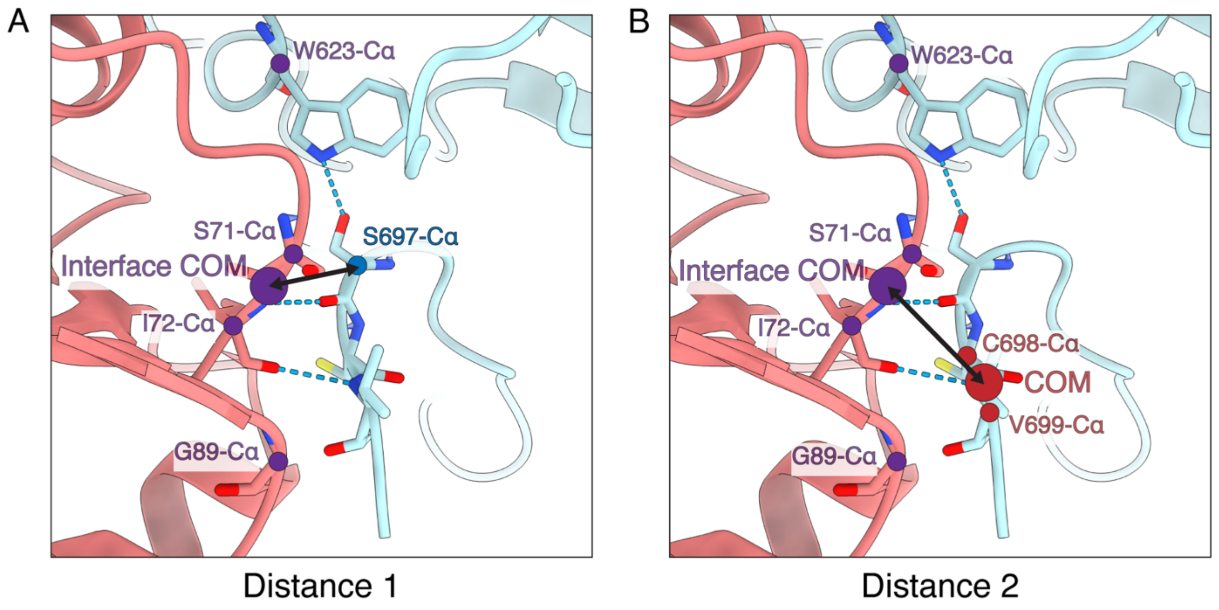


**Figure S13.** Definition of the two distances used in 2D umbrella sampling of α′ C-terminal tail dissociation from TrxA-bound conformation. The interface center-of-mass (COM) is defined as the COM of Cα atoms of W623 in the α′ subunit (light blue) and S71, I72, and G89 in TrxA (coral). (A) Distance 1 is defined as the distance between the interface COM and the Cα atom of S697 in α′. (B) Distance 2 is defined as the distance between the interface COM and the COM of the C698 and V699 Cα atoms in α′.

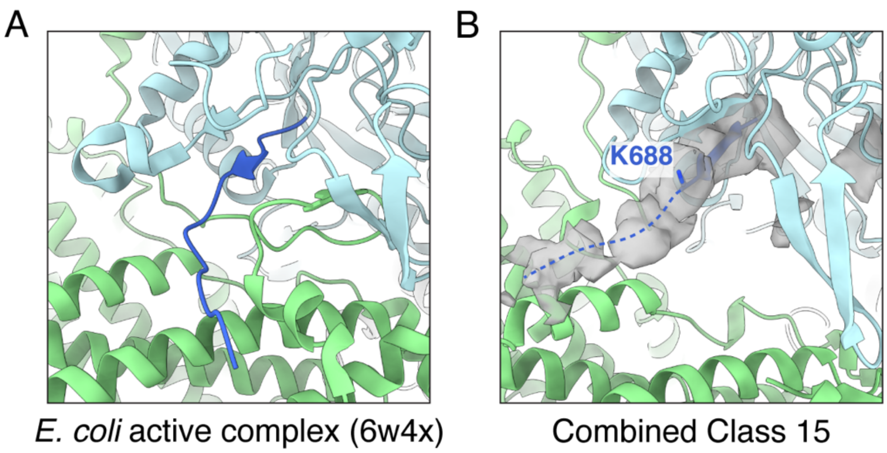


**Figure S14.** Interaction of α C-terminal tail with β subunit. The α subunit is colored in light blue while the β subunit is colored in light green. C-terminal residues after 731 in the *E. coli* RNR α subunit (pdb: 6w4x) and after 684 in the *B. subtilis* RNR α subunit (combined class 15 in Figure S7E) are colored in dark blue. (A) The *E. coli* RNR active complex shows the α C-terminal tail binding to the β subunit on the PCET side. (B) Combined class 15 exhibits extra density close to the β subunit after the last modeled α residue K688, indicating that the α C-terminal tail also interacts with the β subunit in *B. subtilis* RNR.

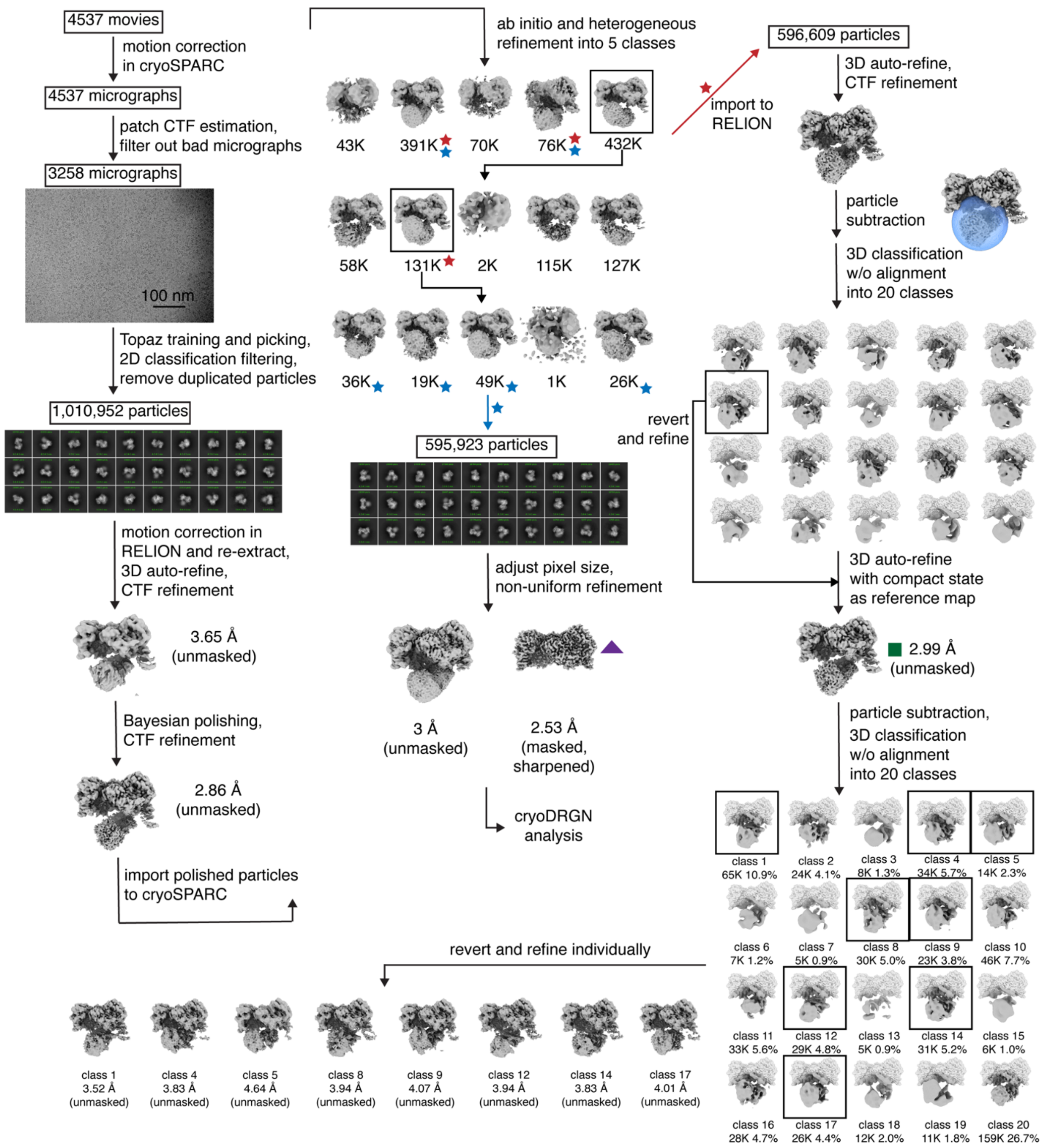


**Figure S15.** EM processing workflow for the turnover dataset. Initial processing led to a 1.011M particle set. Following CTF parameter refinement and Bayesian polishing, particles were further filtered through several rounds of multi-class *ab initio* reconstruction and heterogeneous refinement to produce two similar particle sets. Classes marked with blue stars were combined to produce a 596K particle set (blue arrow) that yielded a consensus reconstruction of the $\alpha_2$ subunit bound with nucleotides and β C-terminus (purple triangle, Figure S16A, Table S1) and routed for downstream cryoDRGN analysis (Figure S23, S25). Classes marked with red stars yielded a 597K particle set for focused classification of the $\beta_2$ subunit, where a consensus reconstruction (green square) was first produced to reduce misalignment caused by pseudosymmetry before ultimately producing 8 refined 3D classes (Figure S16B, Table S2).

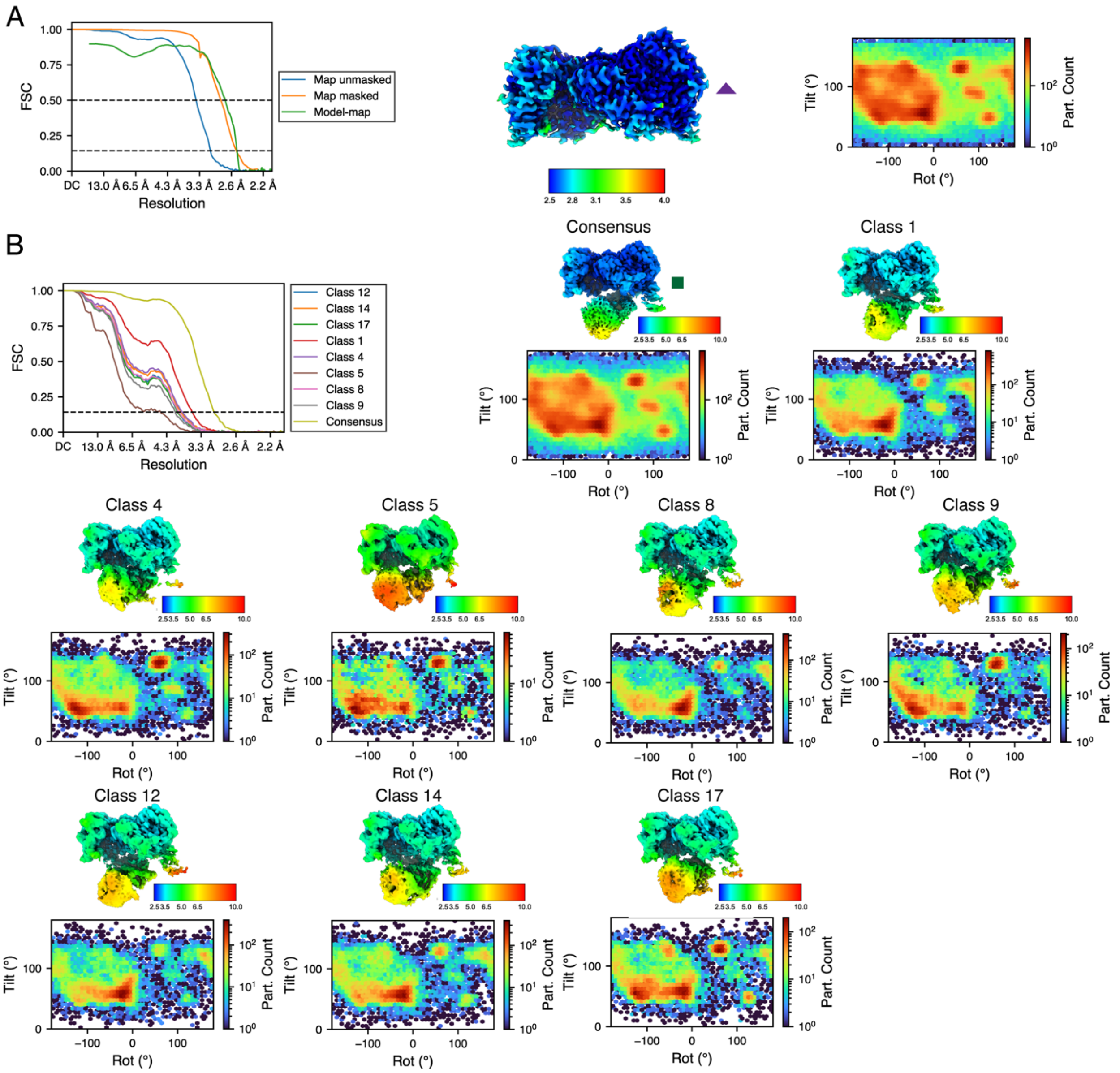


**Figure S16.** Fourier shell correlations (FSCs), local resolutions, and angular distributions for maps obtained for the turnover condition. (A) Consensus refinement in cryoSPARC, which was used for refining the consensus atomic model of the $\alpha_2$ subunit bound with nucleotides and β C-terminus. (B) Unsharpened maps from RELION for the consensus reconstruction and 3D classes, which were used for rigid-body docking of full-complex models.

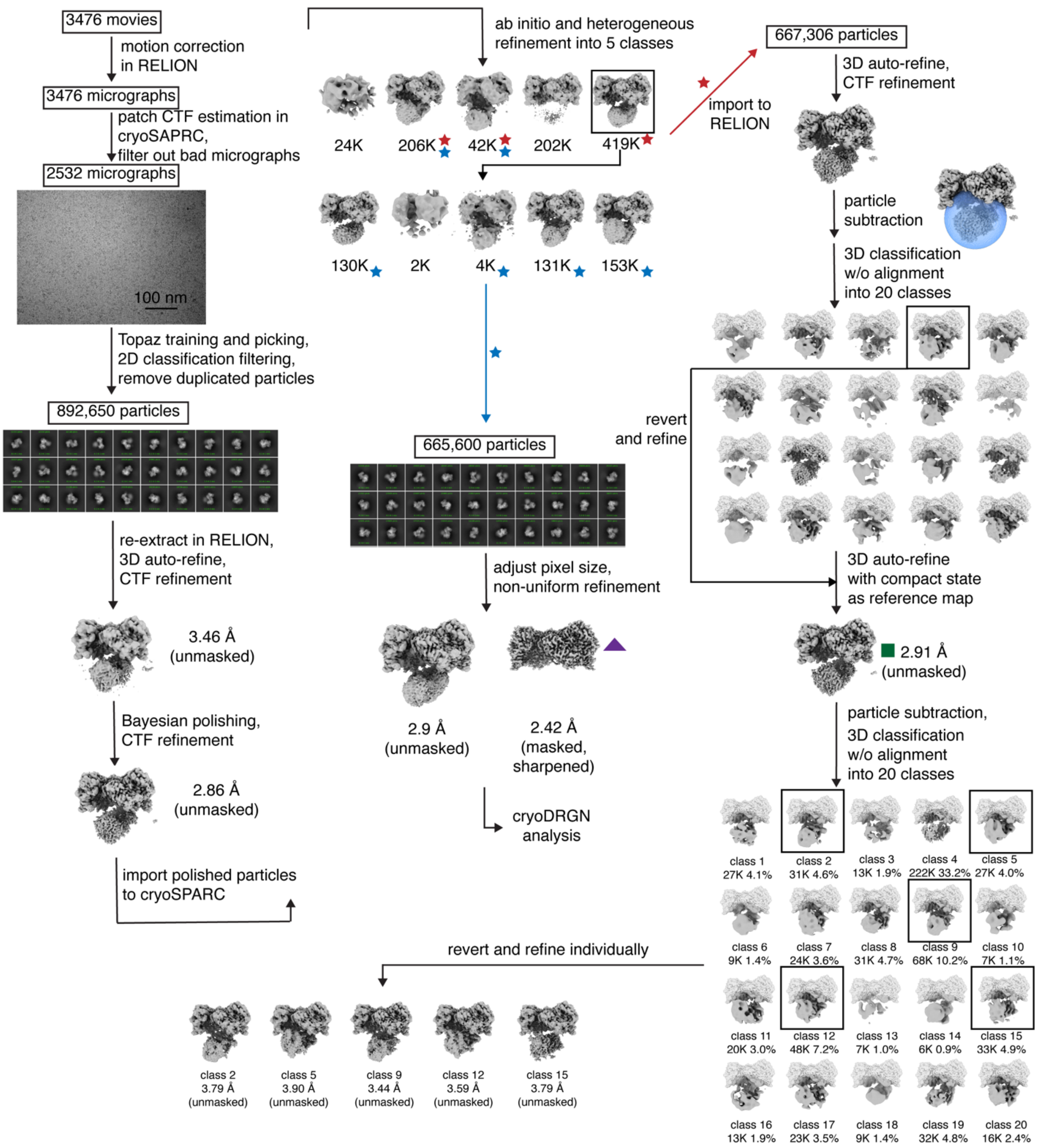


**Figure S17.** EM processing workflow for the pre-turnover dataset. Initial processing led to an 892K particle set. Following CTF parameter refinement and Bayesian polishing, particles were further filtered through two rounds of multi-class *ab initio* reconstruction and heterogeneous refinement to produce two similar particle sets. Classes marked with blue stars were combined to produce a 666K particle set (blue arrow) that yielded a consensus reconstruction of the $\alpha_2$ subunit bound with nucleotides and β C-terminus (purple triangle, Figure S18A, Table S1) and routed for downstream cryoDRGN analysis (Figure S23, S26). Classes marked with red stars yielded a 667K particle set for focused classification of the $\beta_2$ subunit, where a consensus reconstruction (green square) was first produced to reduce misalignment caused by pseudosymmetry before ultimately producing 5 refined 3D classes (Figure S18B, Table S2).

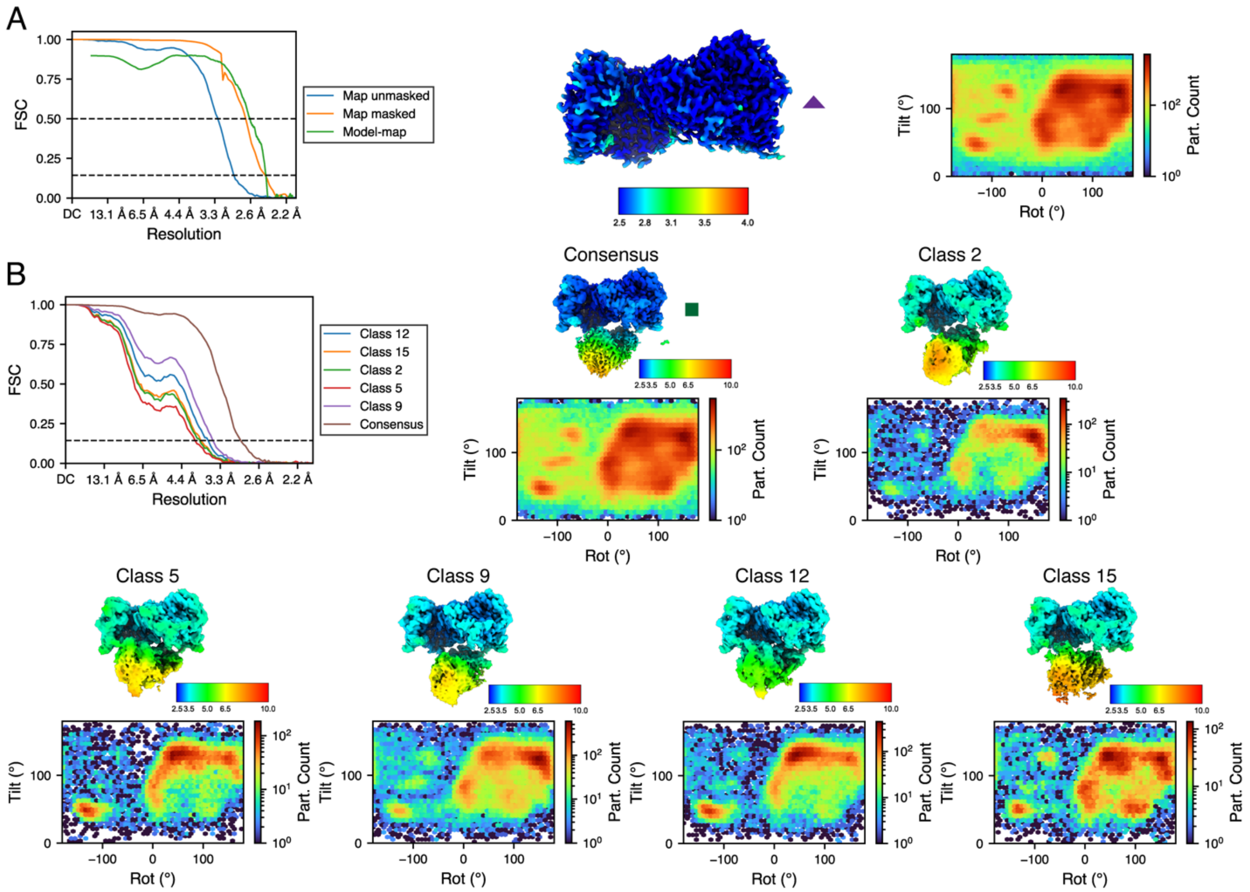


**Figure S18.** Fourier shell correlations (FSCs), local resolutions, and angular distributions for maps obtained for the pre-turnover condition. (A) Consensus refinement in cryoSPARC, which was used for refining the consensus atomic model of the $\alpha_2$ subunit bound with nucleotides and β C-terminus. (B) Unsharpened maps from RELION for the consensus reconstruction and 3D classes, which were used for rigid-body docking of full-complex models.

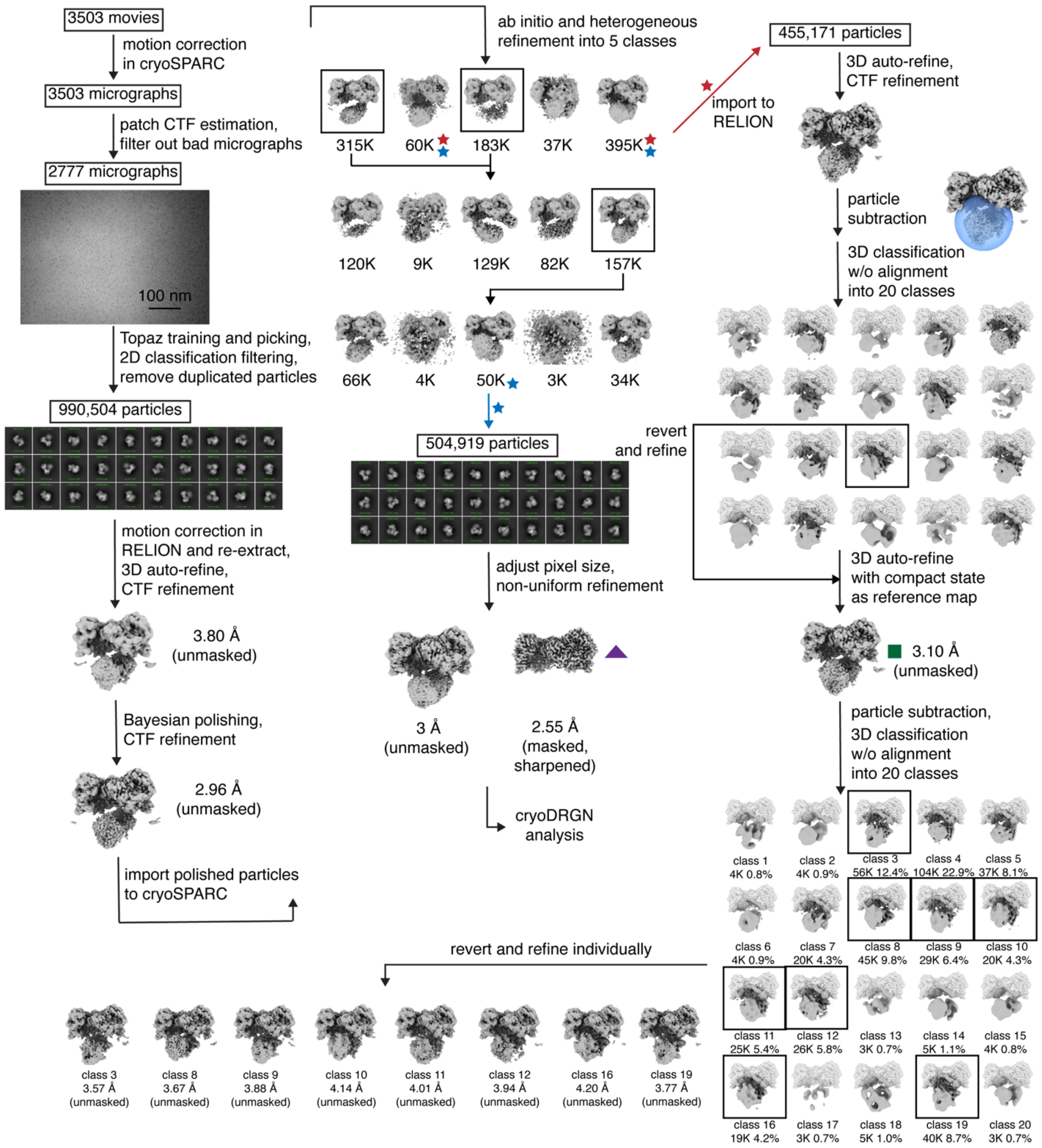


**Figure S19.** EM processing workflow for the product dataset. Initial processing led to a 991K particle set. Following CTF parameter refinement and Bayesian polishing, particles were further filtered through several rounds of multi-class *ab initio* reconstruction and heterogeneous refinement to produce two similar particle sets. Classes marked with blue stars were combined to produce a 505K particle set (blue arrow) that yielded a consensus reconstruction of the $\alpha_2$ subunit bound with nucleotides and β C-terminus (purple triangle, Figure S20A, Table S1) and routed for downstream cryoDRGN analysis (Figure S23, S27). Classes marked with red stars yielded a 455K particle set for focused classification of the $\beta_2$ subunit, where a consensus reconstruction (green square) was first produced to reduce misalignment caused by pseudosymmetry before ultimately producing 8 refined 3D classes (Figure S20B, Table S2).

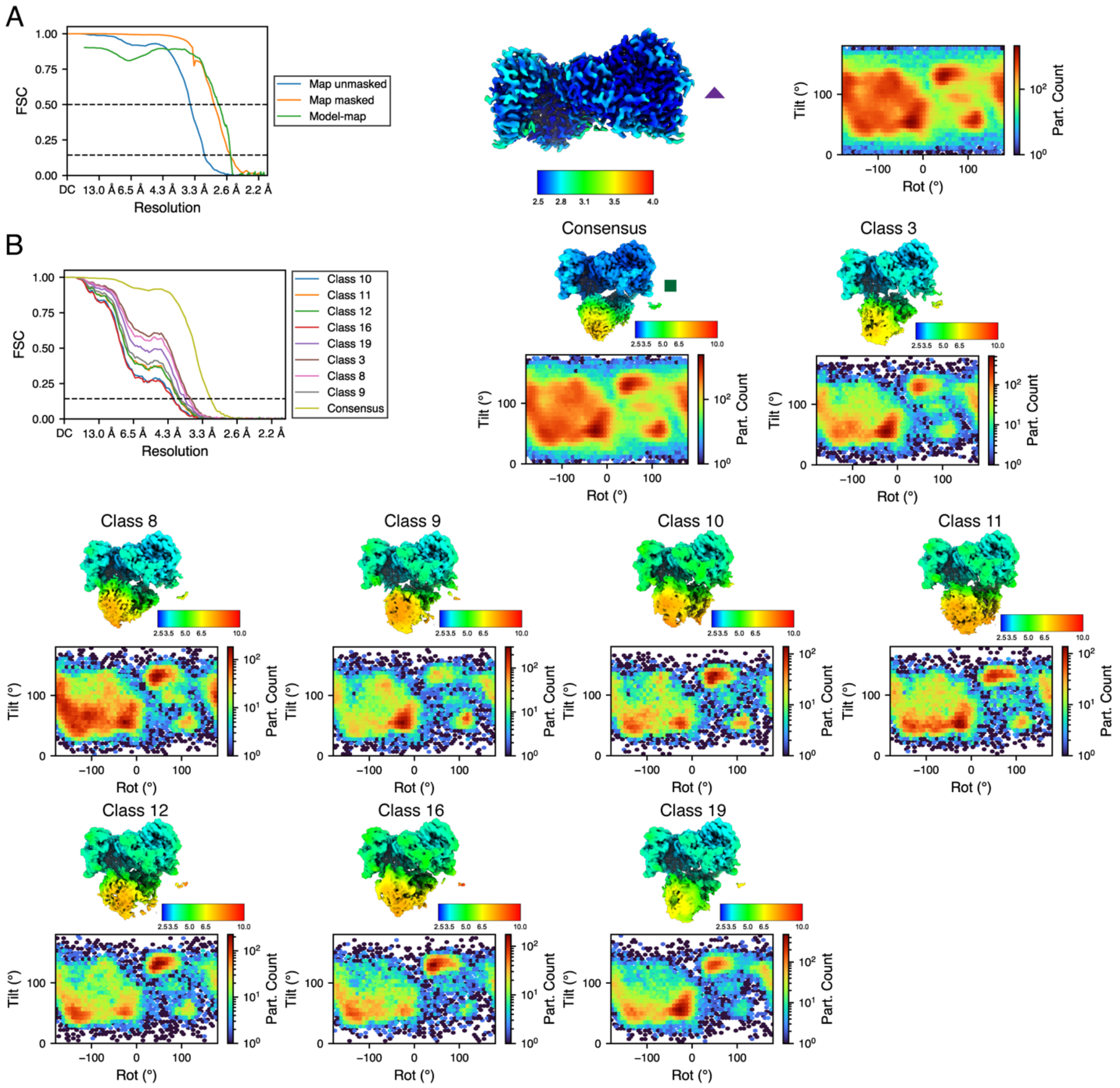


**Figure S20.** Fourier shell correlations (FSCs), local resolutions, and angular distributions for maps obtained for the product condition. (A) Consensus refinement in cryoSPARC, which was used for refining the consensus atomic model of the $\alpha_2$ subunit bound with nucleotides and β C-terminus. (B) Unsharpened maps from RELION for the consensus reconstruction and 3D classes, which were used for rigid-body docking of full-complex models.

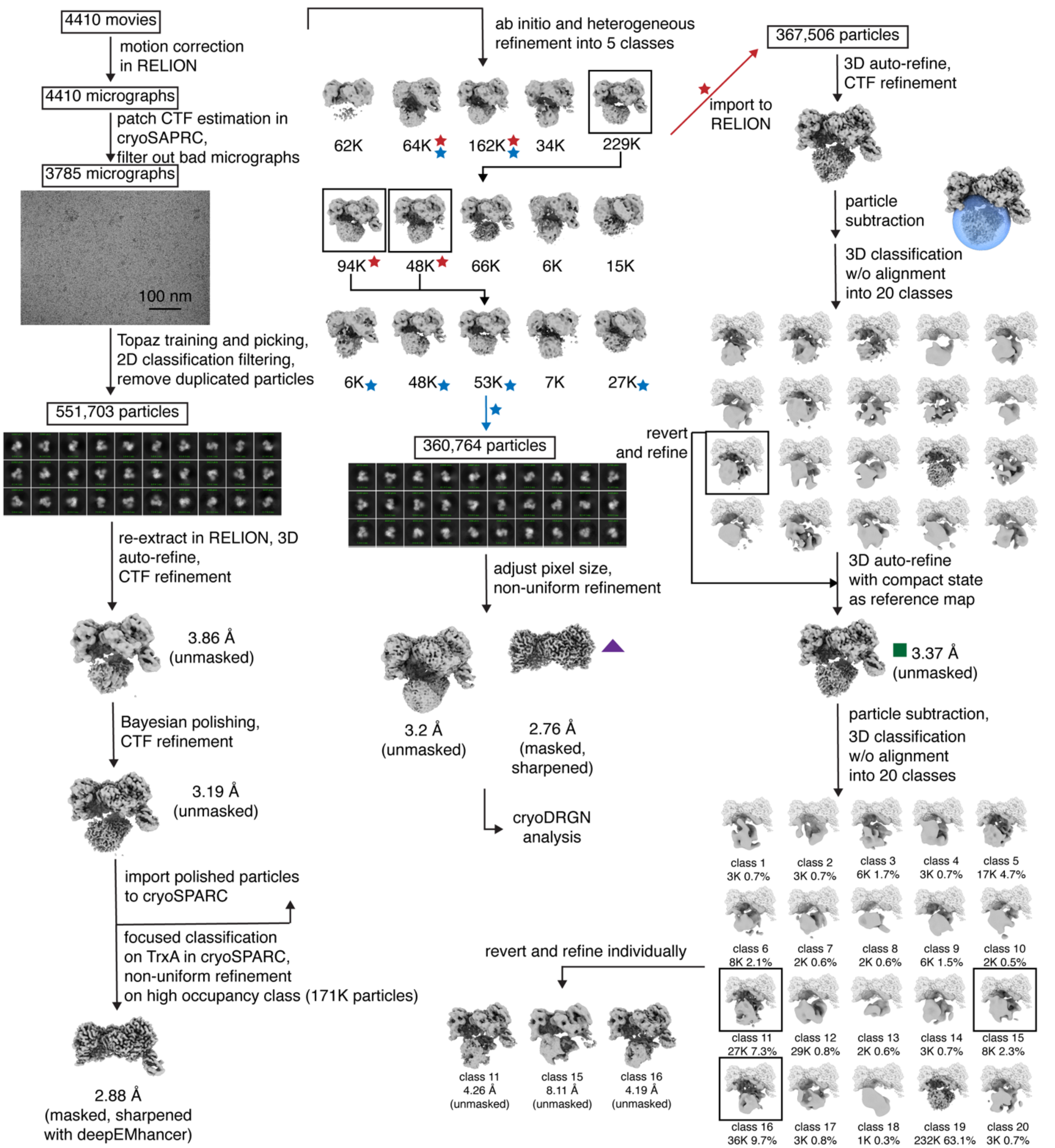


**Figure S21.** EM processing workflow for the pre-reduction dataset. Initial processing led to a 552K particle set. Following CTF parameter refinement and Bayesian polishing, focused classification on TrxA was performed to produce a map of the $\alpha_2$ subunit with TrxA bound on one side (Figure S22B, Table S2). The 552K particle set was further filtered through several rounds of multi-class *ab initio* reconstruction and heterogeneous refinement to produce two similar particle sets. Classes marked with blue stars were combined to produce a 361K particle set (blue arrow) that yielded a consensus reconstruction of the $\alpha_2$ subunit bound with nucleotides and β C-terminus (purple triangle, Figure S22A, Table S1) and routed for downstream cryoDRGN analysis (Figure S23, S28). Classes marked with red stars yielded a 368K particle set for focused classification of the $\beta_2$ subunit, where a consensus reconstruction (green square) was first produced to reduce misalignment caused by pseudosymmetry before ultimately producing 3 refined 3D classes (Figure S22C, Table S2).

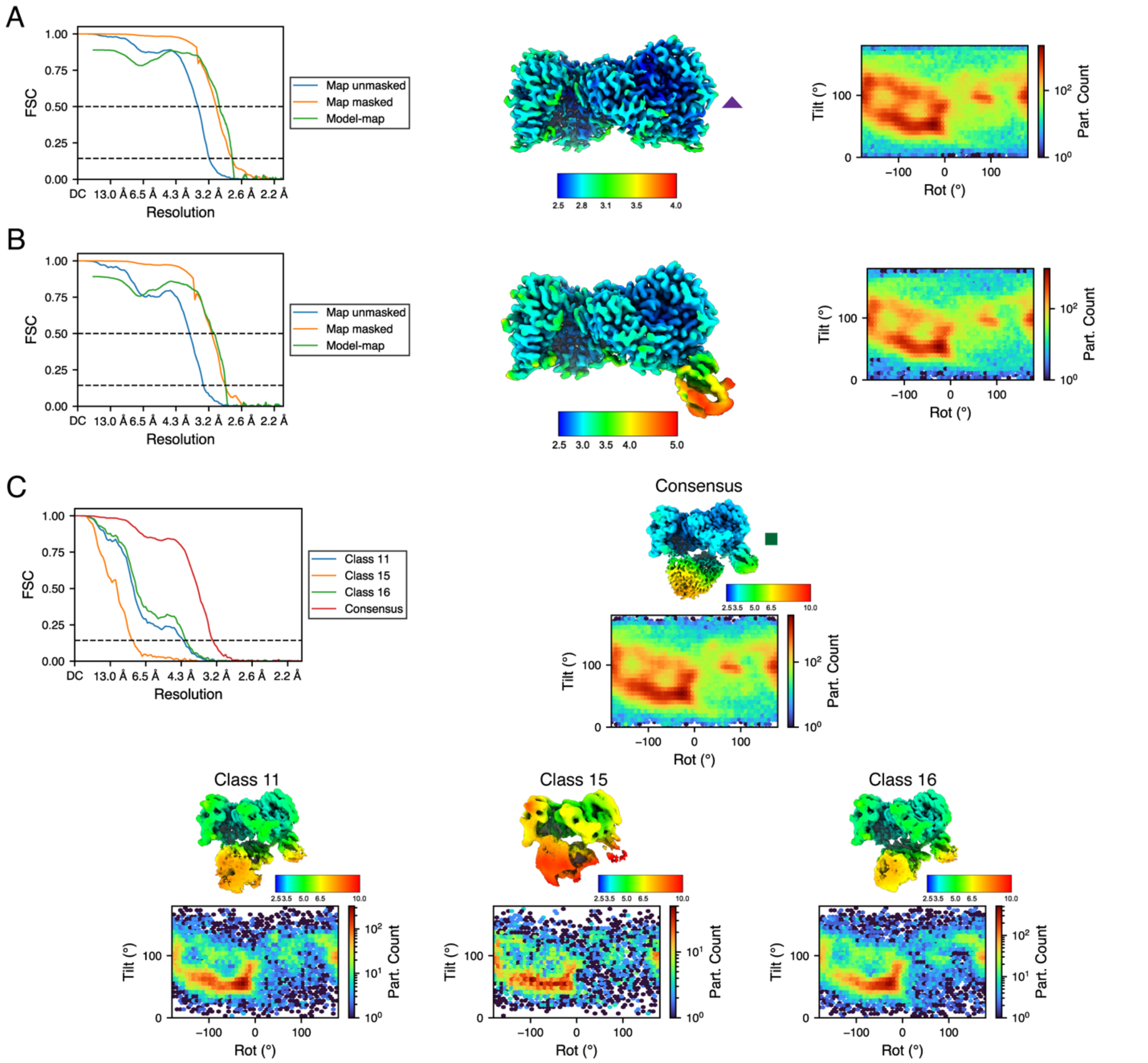


**Figure S22.** Fourier shell correlations (FSCs), local resolutions, and angle distributions for maps obtained for the pre-reduction condition. (A) Consensus refinement in cryoSPARC, which was used for refining the consensus atomic model of the $\alpha_2$ subunit bound with nucleotides and β C-terminus. (B) Refinement after focsed classification on TrxA in cryoSPARC. FSCs were computated with the cryoSPARC sharpened map, while the map shown in the middle was denoised and sharpened with deepEMhancer[42]. (C) Unsharpened maps from RELION for the consensus reconstruction and 3D classes, which were used for rigid-body docking of full-complex models.

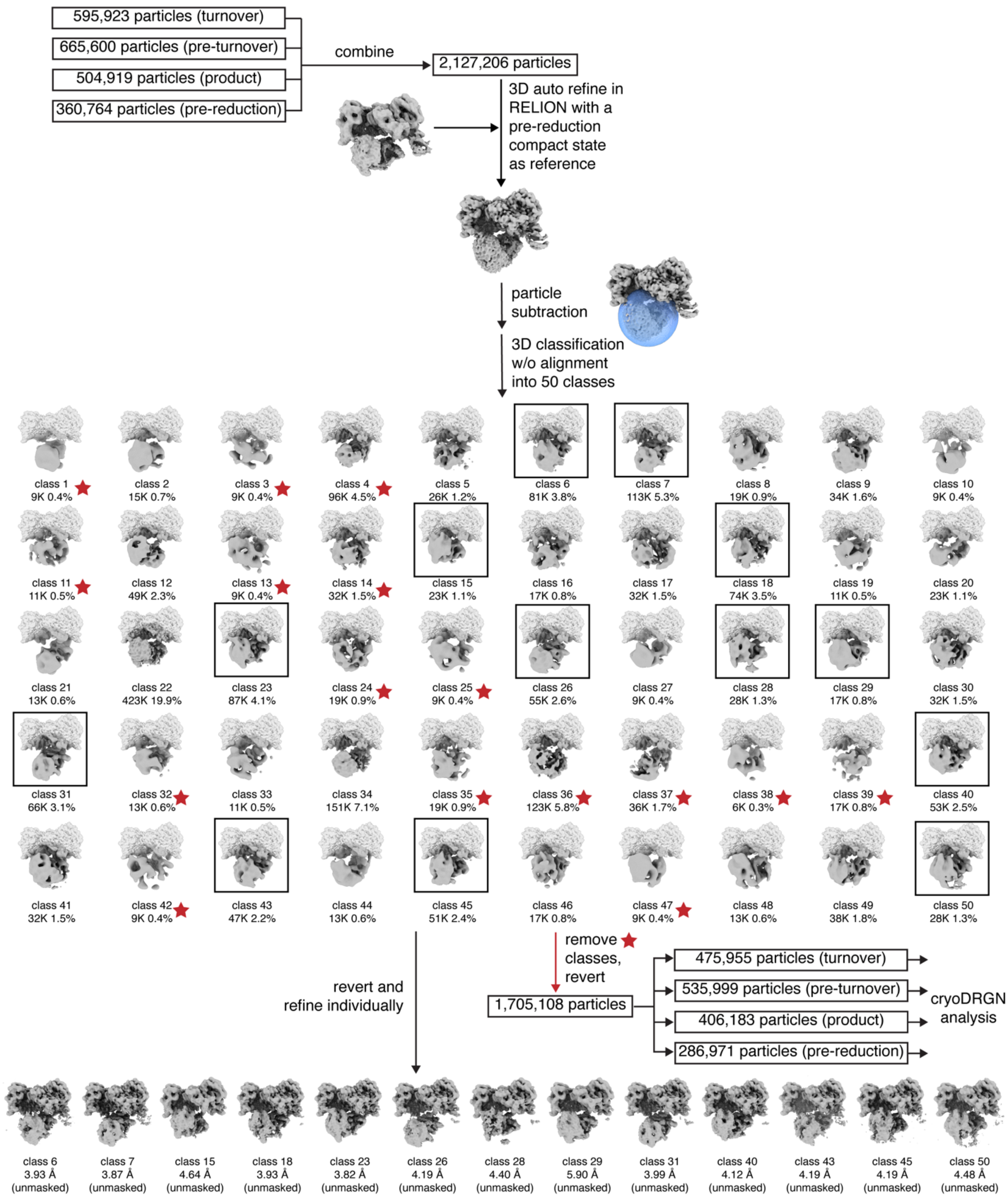


**Figure S23.** Additional particle filtering prior to cryoDRGN analysis through combined refinement and classification of particles from all conditions (Figures S15, S17, S19, S21) in RELION. Classes marked with red stars were filtered to produce 1.705M particles for subsequent cryoDRGN analysis (Figures S25-S28), while classes in black boxes were refined individually to produce 13 3D classes (Figure S24, Table S2).

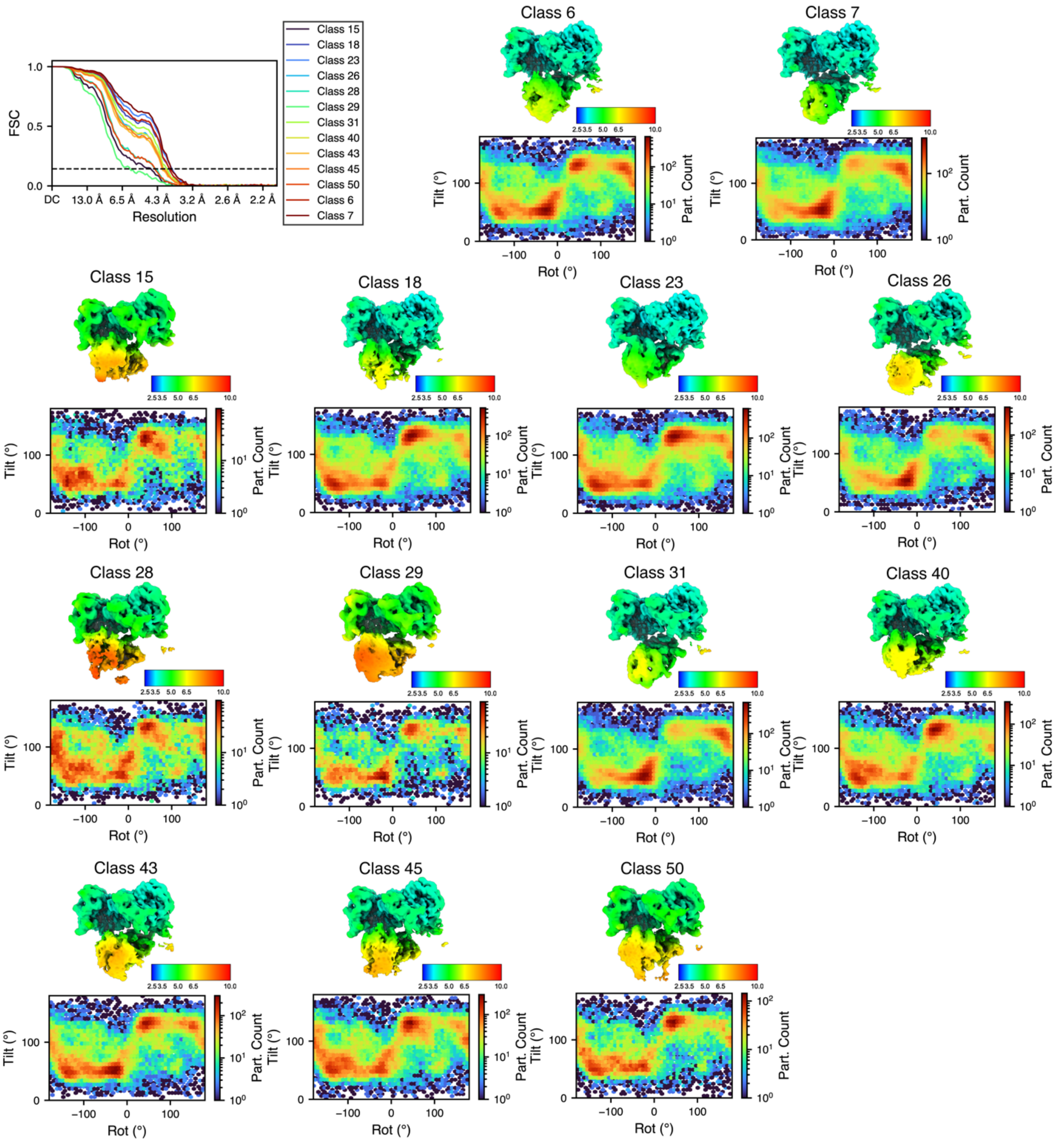


**Figure S24.** Fourier shell correlations (FSCs), local resolutions, and angular distributions for unsharpened maps from RELION for 3D classes obtained from the combined refinement of all conditions.

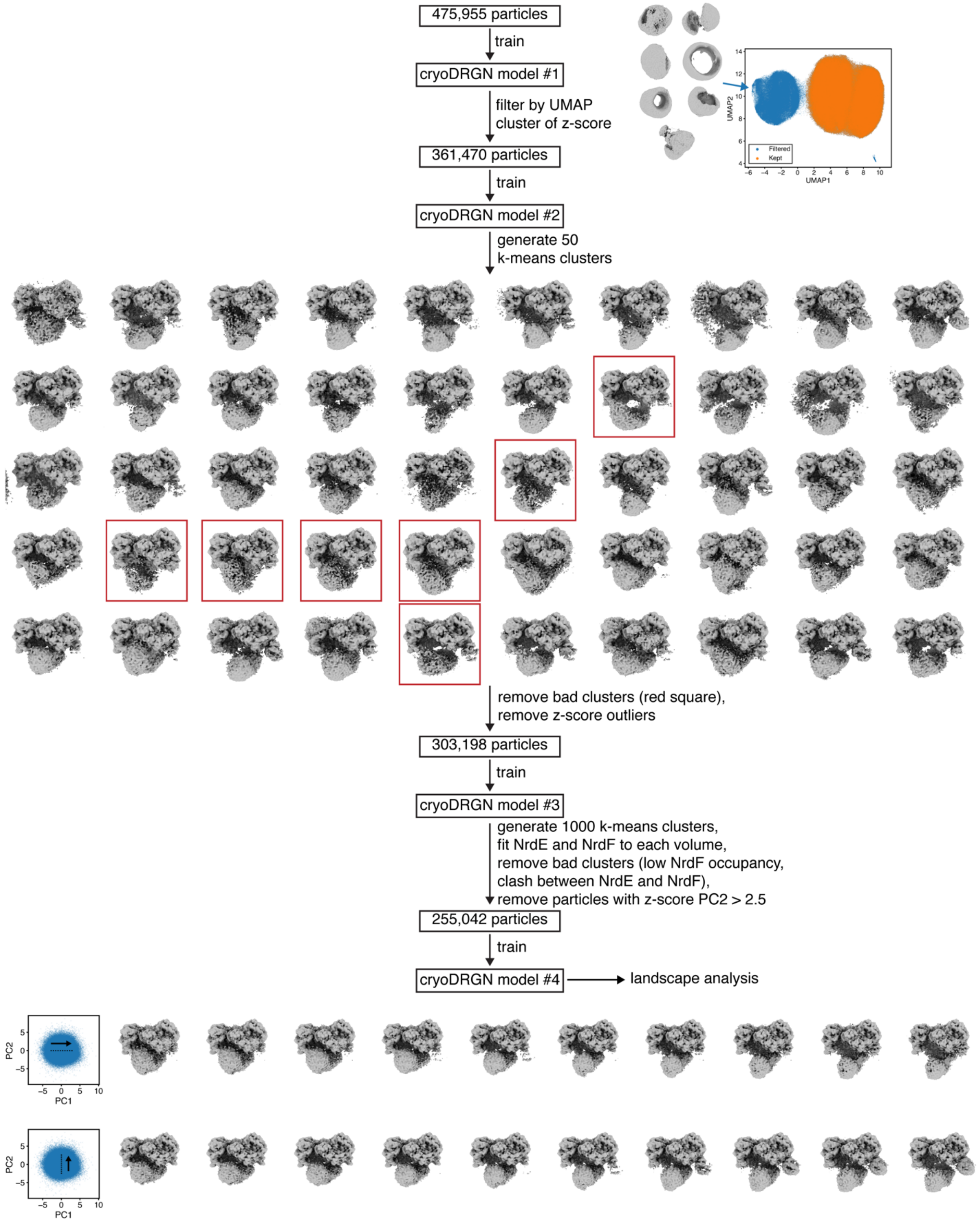


**Figure S25.** CryoDRGN analysis workflow for turnover condition. Junk particles and particles with poor β2 density were filtered through four rounds of cryoDRGN model training. Volumes along the first two principal components (Movie S1) of the latent space for the final cryoDRGN model are shown in the bottom.

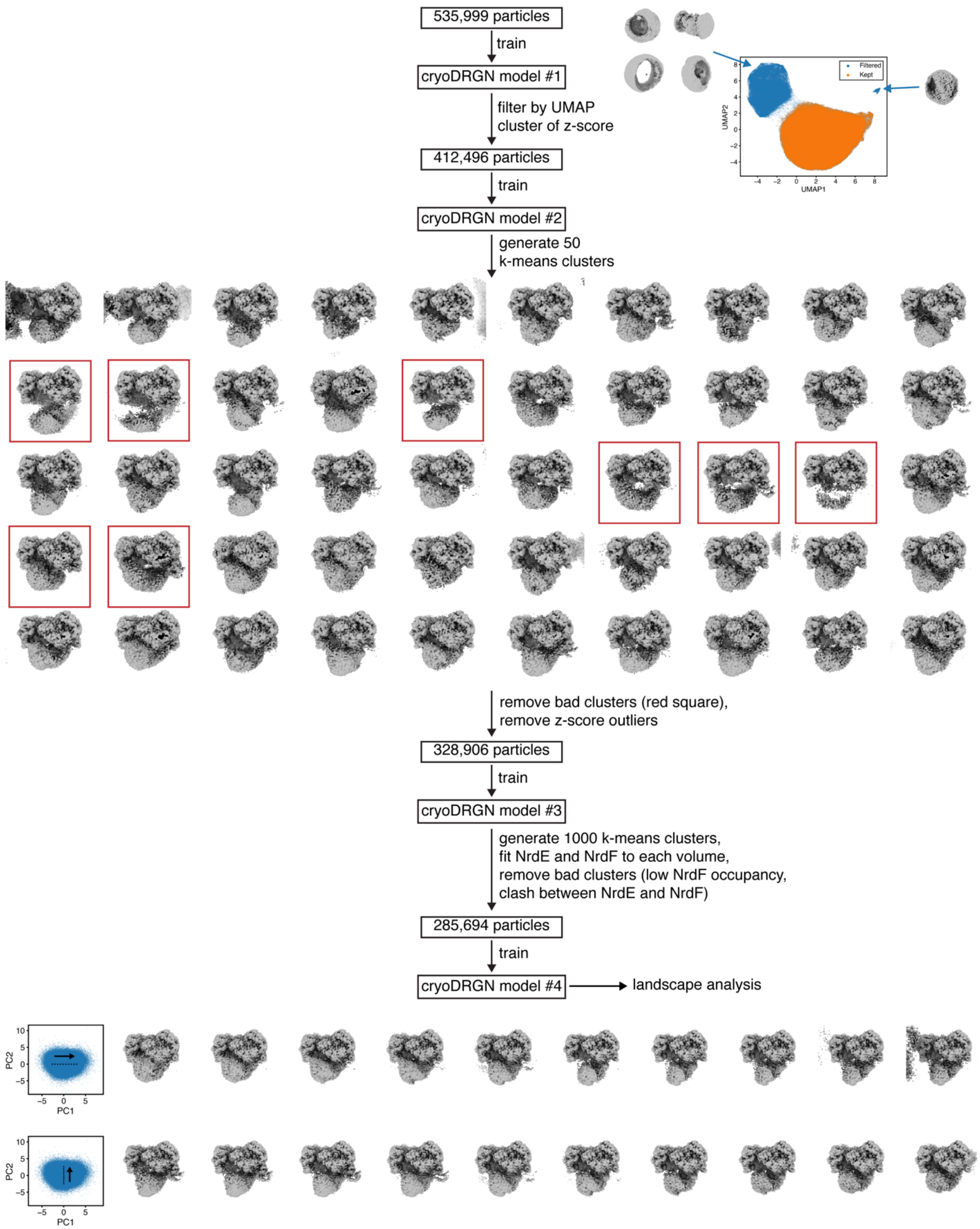


**Figure S26.** CryoDRGN analysis workflow for pre-turnover condition. Junk particles and particles with poor $\beta_2$ density were filtered through four rounds of cryoDRGN model training. Volumes along the first two principal components (Movie S2) of the latent space for the final cryoDRGN model are shown in the bottom.

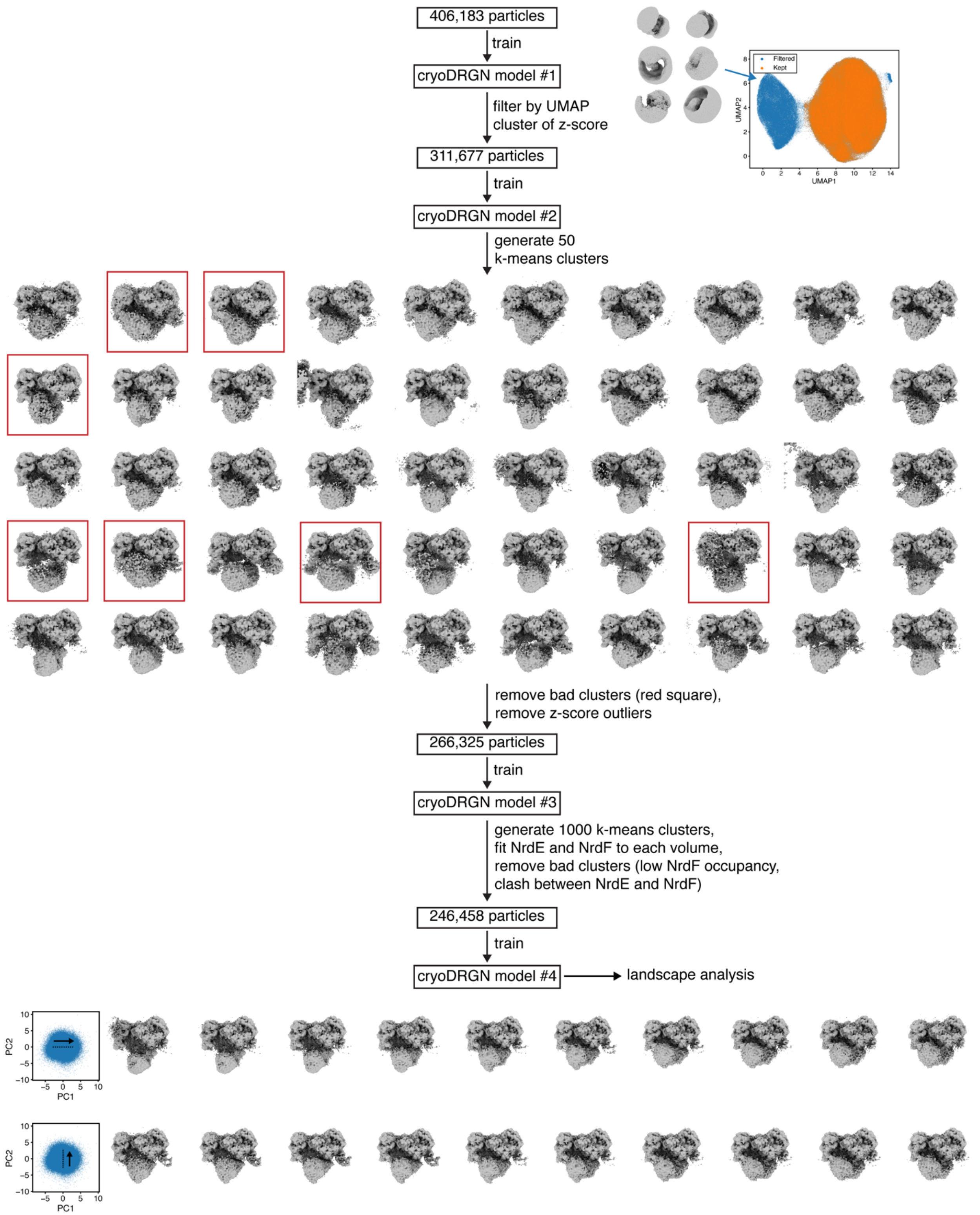


**Figure S27.** CryoDRGN analysis workflow for product condition. Junk particles and particles with poor $\beta_2$ density were filtered through four rounds of cryoDRGN model training. Volumes along the first two principal components (Movie S3) of the latent space for the final cryoDRGN model are shown in the bottom.

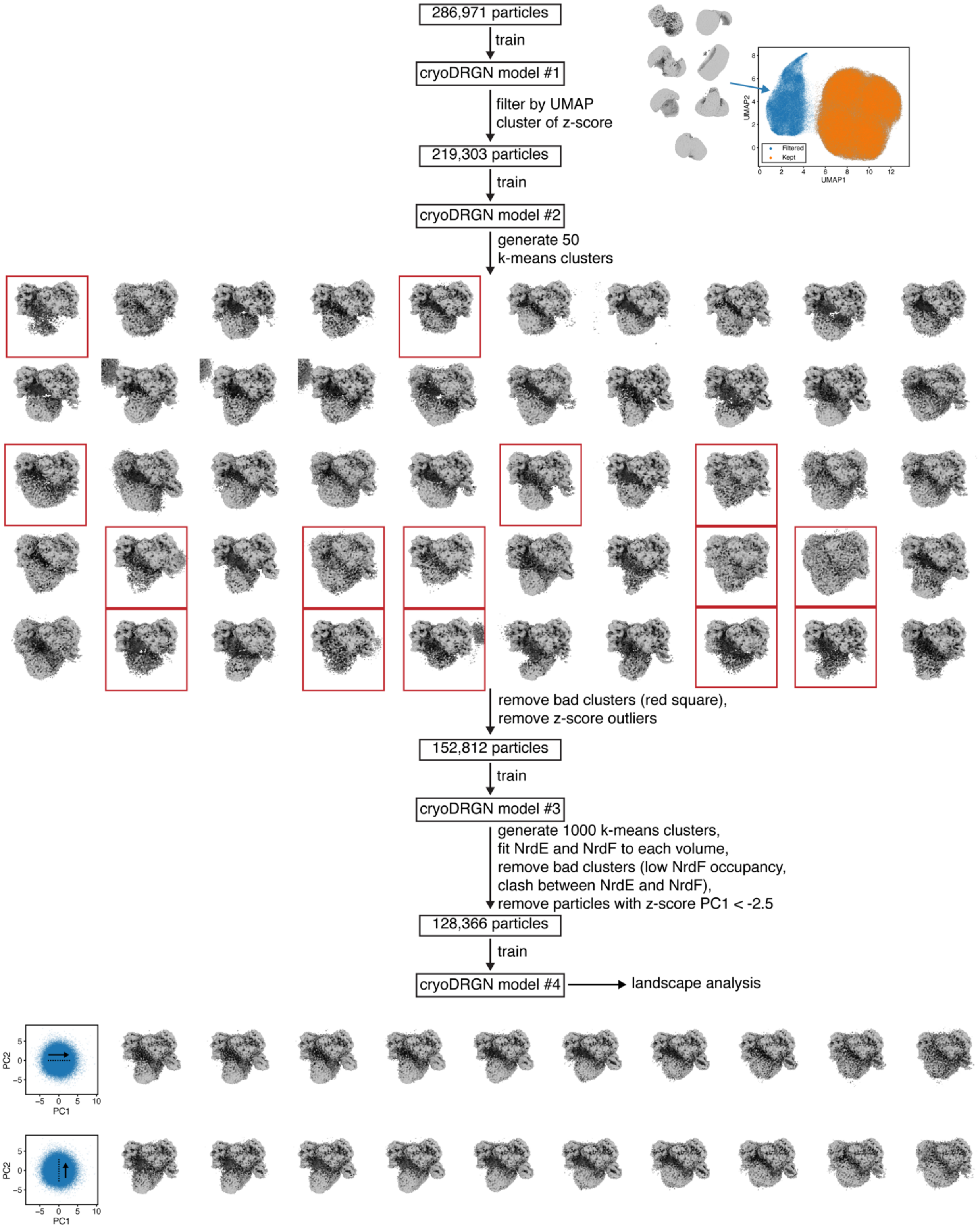


**Figure S28.** CryoDRGN analysis workflow for pre-reduction condition. Junk particles and particles with poor β2 density were filtered through four rounds of cryoDRGN model training. Volumes along the first two principal components (Movie S4) of the latent space for the final cryoDRGN model are shown in the bottom.

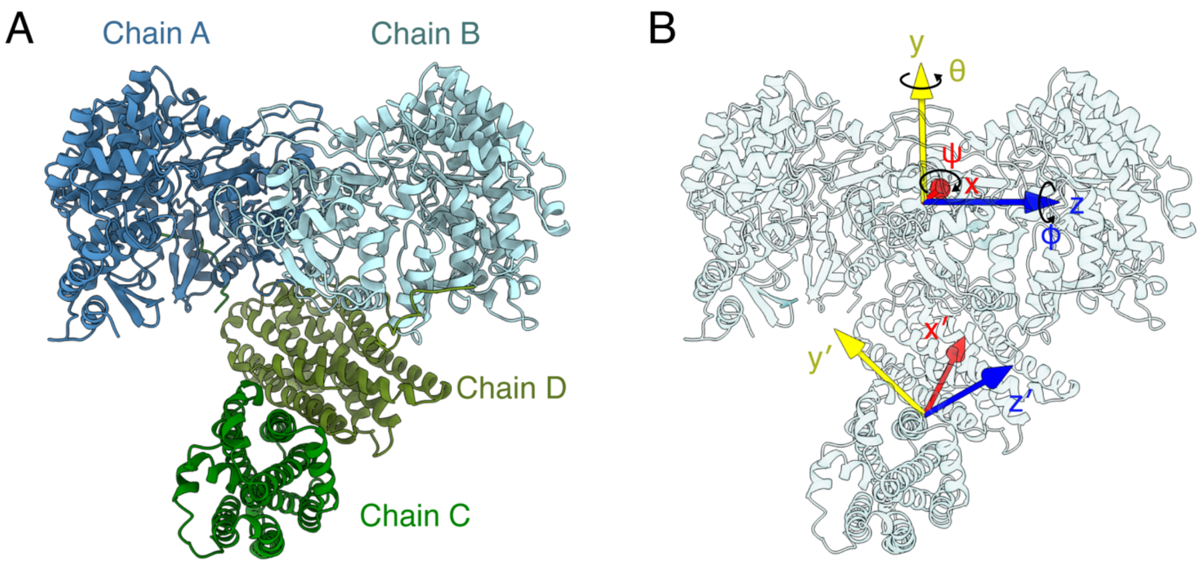


**Figure S29.** Parametrization of inter-subunit orientations in the $\alpha_2\beta_2$ complex for landscape analysis. (A) Chain IDs for the full-complex models. (B) Two coordinate systems defined for each model, with *x y z* being the first coordinate system based on $\alpha_2$ and *x′ y′ z′* being the second coordinate system based on $\beta_2$. Euler angles *ψ, θ, φ* correspond to extrinsic rotation about the *x y z* axes, respectively, needed to bring the two coordinate systems in alignment.

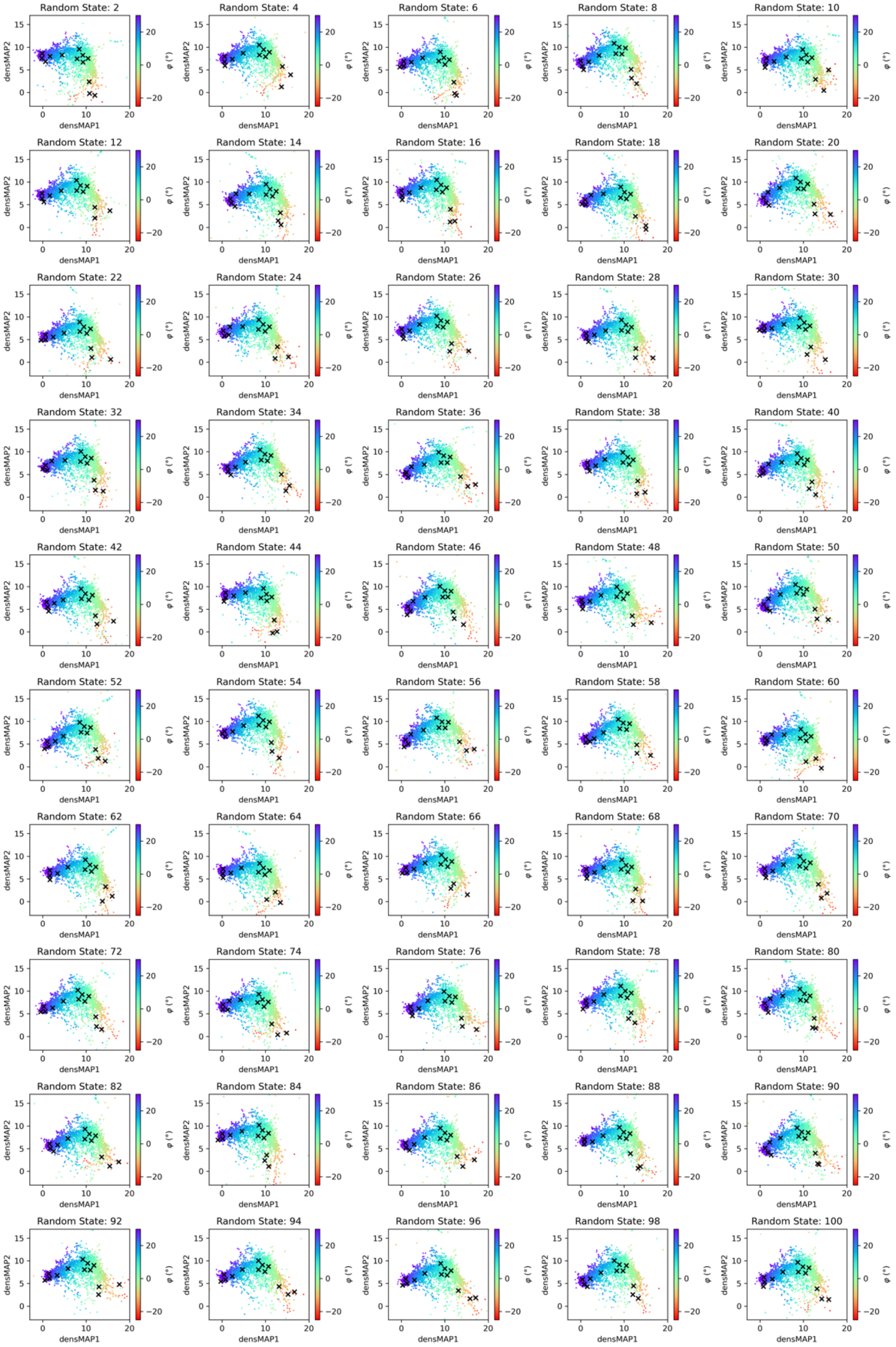


**Figure S30.** DensMAP 2D embeddings with different random seeds. The majority of the embedding is very stable, while the embedding around the PCET-like region shows slight variability in position within the lower right quadrant.

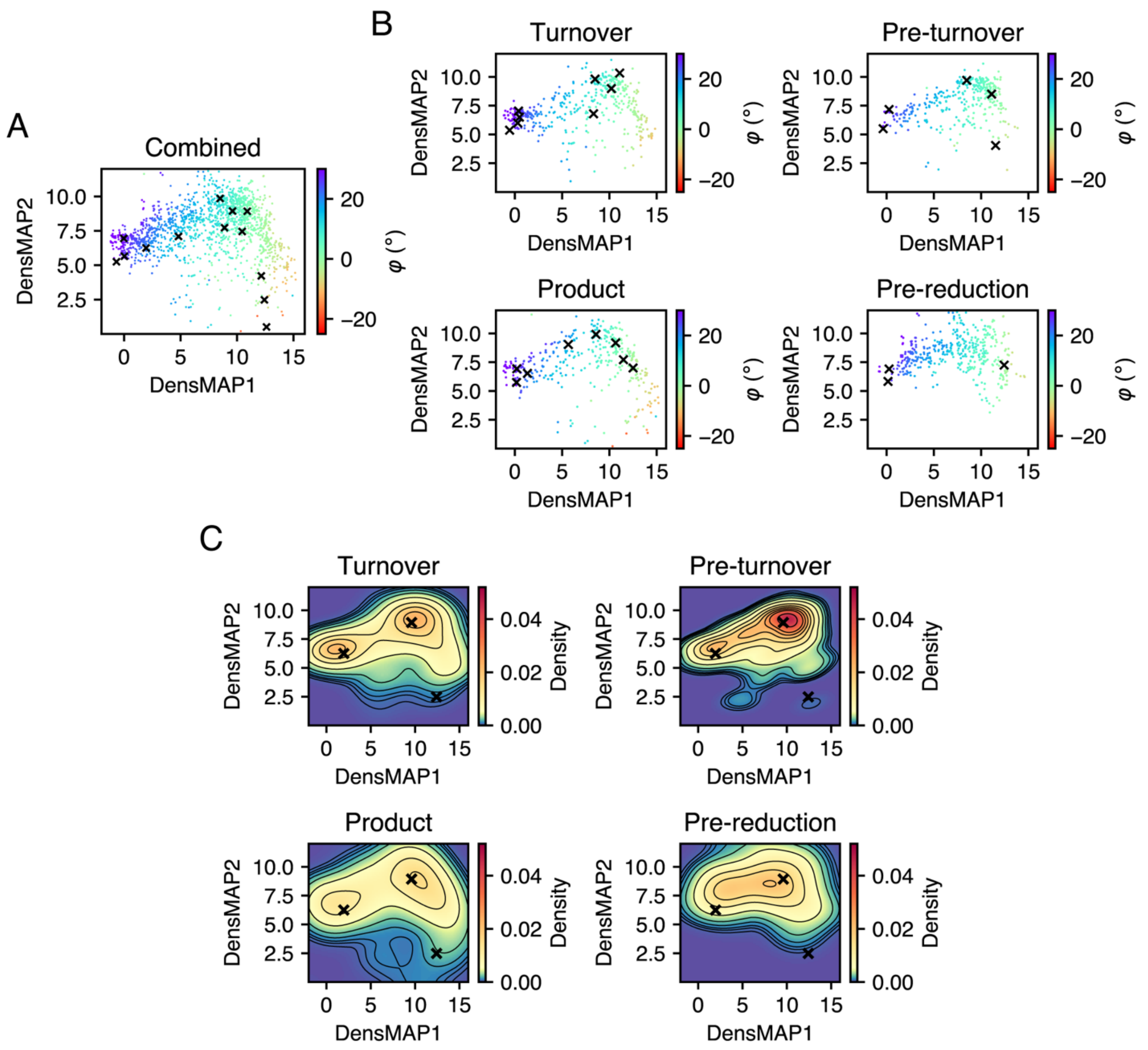


**Figure S31.** Landscape analysis repeated with a $k$-means clustering of $k$=500. (A) Combined 2D representation colored by Euler angle $\varphi$ and plotted with 3D classes from the combined refinement marked as crosses. (B) Individual 2D representations colored by Euler angle $\varphi$ and plotted with 3D classes from individual refinements marked as crosses. (C) Landscapes estimated from individual 2D representations, with the same three 3D classes marked as in Figure 4. That we obtain similar results with a larger $k$-means clustering of $k$=1000 (Figures 3, S8-9) indicates that the latent space was sufficiently sampled in our main analysis.

**Table S4.** Initial system setup for different conditions.

| | Oxidized $\alpha_2/\beta_2$ | Reduced $\alpha_2/\beta_2$ | Oxidized $\alpha_2/\beta_2$/TrxA |
|---|---|---|---|
| Box shape | Truncated octahedral | Truncated octahedral | Truncated octahedral |
| Box dimensions | 137.7Å, 137.7Å, 137.7Å 109.5°, 109.5°, 109.5° | 137.7Å, 137.7Å, 137.7Å 109.5°, 109.5°, 109.5° | 150.7Å, 150.7Å, 150.7Å 109.5°, 109.5°, 109.5° |
| Total atoms | 227774 | 227594 | 304473 |
| Total water molecules | 48367 | 48320 | 67122 |
| Salt concentration | 150 mM NaCl 15 mM $MgCl_2$ | 150 mM NaCl 15 mM $MgCl_2$ | 150 mM NaCl 15 mM $MgCl_2$ |

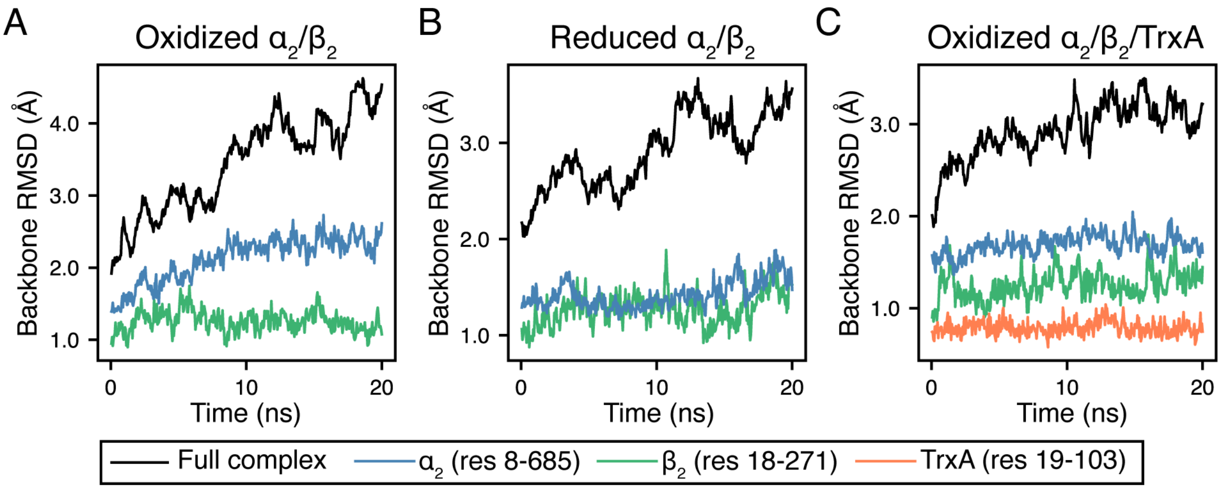


**Figure S32.** Backbone root-mean square deviation (RMSD) of entire complex or rigid parts of individual proteins during 20 ns unconstrained equilibration MD runs. (A-B) Oxidized and reduced $\alpha_2$ complexed with $\beta_2$ for F624 $\chi_1$ angle umbrella sampling. (C) Oxidized $\alpha_2$ complexed with $\beta_2$ and TrxA for 2D umbrella sampling of α tail. These results show that individual protein components were well-equilibrated after 10 ns in all runs, although the full complex exhibited larger deviations due to the flexible tails and drift in orientation between $\alpha_2$ and $\beta_2$.

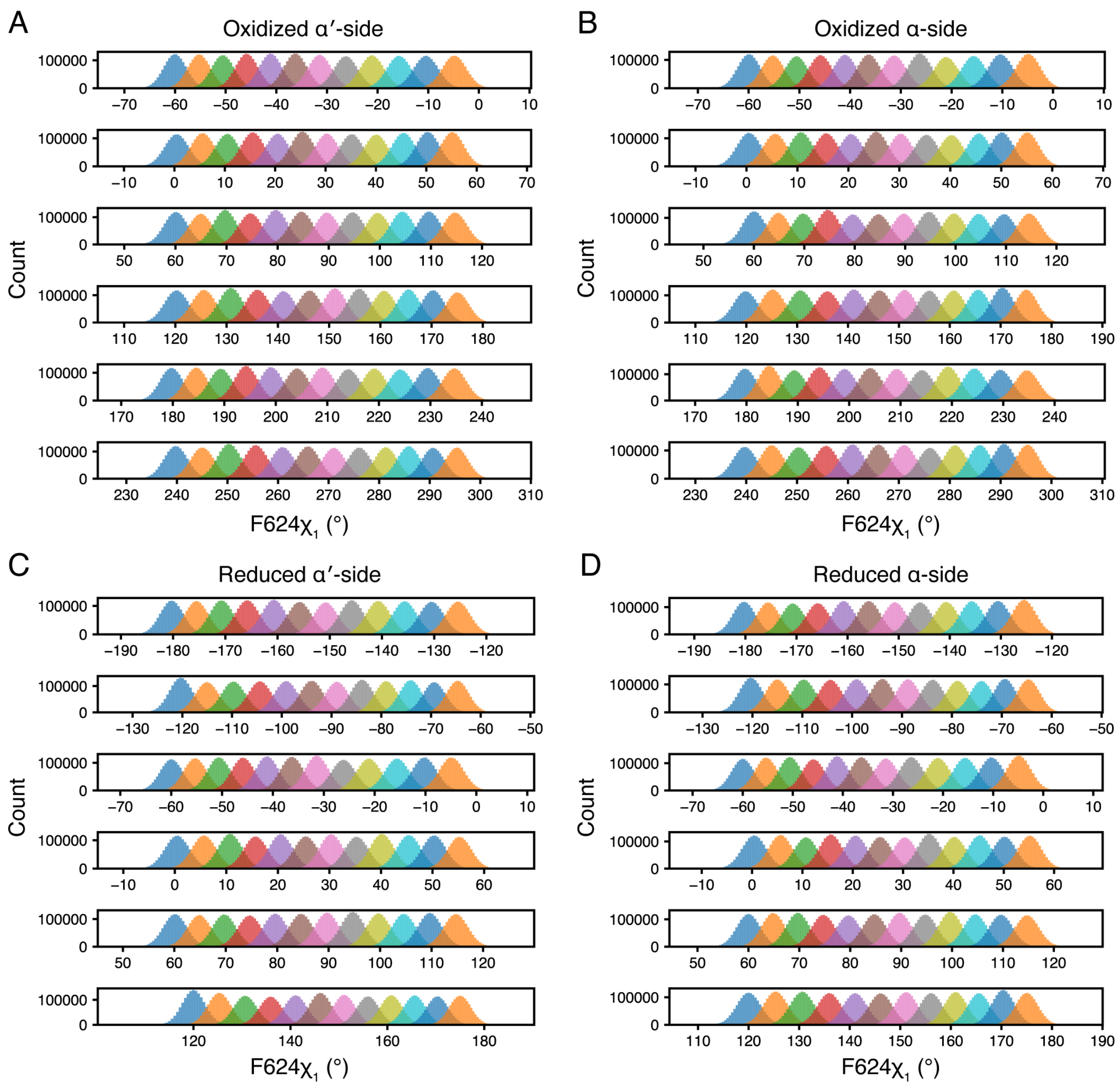


**Figure S33.** Histograms of sampled F624 $\chi_1$ angles in each simulation window for (A) α′ side of $\alpha_2$ with oxidized active site (B) α side of $\alpha_2$ with oxidized active site (C) α′ side of $\alpha_2$ with reduced active site (D) α side of $\alpha_2$ with reduced active site. Histograms show good overlap between adjacent windows, indicating sufficient sampling.

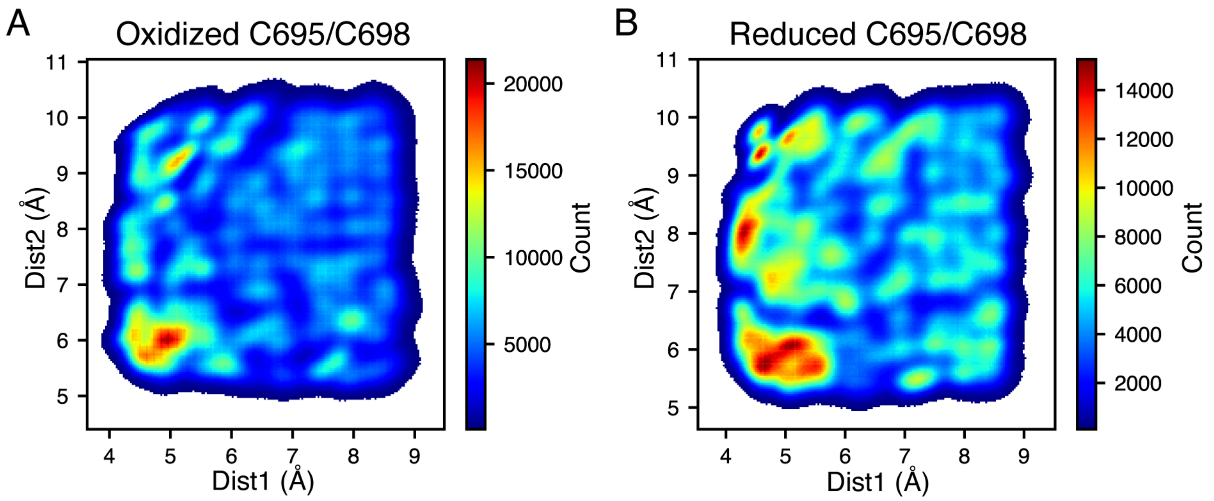


**Figure S34.** 2D histograms of sampled distances in 2D umbrella sampling for (A) oxidized and (B) reduced C695/C698 pair on the α′ C-terminus. Both histograms show sufficient sampling across the two distances.

**Supplementary Movies**

**Movie S1.** Volumes along the first two principal components PC1 (left) and PC2 (right) for the final cryoDRGN model of the turnover condition. 10 volumes were uniformly sampled along each principal component between the point corresponding to the 5th percentile of all points and the point corresponding to the 95th percentile (Figure S25). The movie steps through volumes starting from the 5th percentile point to the 95th percentile point of each principal component before returning.

**Movie S2.** Volumes along the first two principal components PC1 (left) and PC2 (right) for the final cryoDRGN model of the pre-turnover condition. 10 volumes were uniformly sampled along each principal component between the point corresponding to the 5th percentile of all points and the point corresponding to the 95th percentile (Figure S26). The movie steps through volumes starting from the 5th percentile point to the 95th percentile point of each principal component before returning.

**Movie S3.** Volumes along the first two principal components PC1 (left) and PC2 (right) for the final cryoDRGN model of the product condition. 10 volumes were uniformly sampled along each principal component between the point corresponding to the 5th percentile of all points and the point corresponding to the 95th percentile (Figure S27). The movie steps through volumes starting from the 5th percentile point to the 95th percentile point of each principal component before returning.

**Movie S4.** Volumes along the first two principal components PC1 (left) and PC2 (right) for the final cryoDRGN model of the pre-reduction condition. 10 volumes were uniformly sampled along each principal component between the point corresponding to the 5th percentile of all points and the point corresponding to the 95th percentile (Figure S28). The movie steps through volumes starting from the 5th percentile point to the 95th percentile point of each principal component before returning.
